# Bioinformatics and next generation data analysis reveals the potential role of inflammation in sepsis and its associated complications

**DOI:** 10.1101/2023.08.02.551653

**Authors:** Basavaraj Vastrad, Chanabasayya Vastrad

## Abstract

Sepsis is the leading systemic inflammatory response syndrome in worldwide, yet relatively little is known about the genes and signaling pathways involved in sepsis progression. The current investigation aimed to elucidate potential key candidate genes and pathways in sepsis and its associated complications. Next generation sequencing (NGS) dataset (GSE185263) was downloaded from the Gene Expression Omnibus (GEO) database, which included data from 348 sepsis samples and 44 normal control samples. Differentially expressed genes (DEGs) were identified using t-tests in the DESeq2 R package. Next, we made use of the g:Profiler to analyze gene ontology (GO) and REACTOME pathway. Then protein-protein interaction (PPI) of these DEGs was visualized by Cytoscape with Search Tool for the Retrieval of Interacting Genes (STRING). Furthermore, we constructed miRNA-hub gene regulatory network and TF-hub gene regulatory network among hub genes utilizing miRNet and NetworkAnalyst online databases tool and Cytoscape software. Finally, we performed receiver operating characteristic (ROC) curve analysis of hub genes through the pROC package in R statistical software. In total, 958 DEGs were identified, of which 479 were up regulated and 479 were down regulated. GO and REACTOME results showed that DEGs mainly enriched in regulation of cellular process, response to stimulus, extracellular matrix organization and immune system. The hub genes of PRKN, KIT, FGFR2, GATA3, ERBB3, CDK1, PPARG, H2BC5, H4C4 and CDC20 might be associated with sepsis and its associated complications. Predicted miRNAs (e.g., hsa-mir-548ad-5p and hsa-mir-2113) and TFs (e.g., YAP1 and TBX5) were found to be significantly correlated with sepsis and its associated complications. In conclusion, the DEGs, relative pathways, hub genes, miRNA and TFs identified in the current investigation might help in understanding of the molecular mechanisms underlying sepsis and its associated complications progression and provide potential molecular targets and biomarkers for sepsis and its associated complications.

## Introduction

Sepsis is the most important systemic inflammatory response syndrome caused by a dysregulated host response to microbial infection [1]. Sepsis was estimated to affect over 49 million people worldwide and resulted in approximately 11 million deaths in 2017 [2]. Sepsis becomes the chief cause of the life-threatening organ dysfunction [3]. Sepsis is broadly characterized as a cascade of immune system activation and widespread inflammation throughout the body [4]. The crucial features of sepsis include infection [5], systemic inflammation [6], altered body temperature [7], altered heart function [8], altered lung function [9], altered brain function [10], altered kidney function [11], altered liver function [12] and altered blood pressure [13]. Septic shock characterized by extremely low blood pressure (hypotension) and inadequate blood flow to vital organs. Septic shock can lead to multiple organ failure [14] and has a high mortality rate [15]. Sepsis is often considered one of the leading causes of death globally, especially in intensive care units (ICUs) and hospitals [16]. Sepsis can affect individuals of all ages, from newborns to the elderly [17]. Sepsis poses a significant burden on healthcare systems worldwide [18]. Sepsis is a particularly challenging problem in developing countries due to limited access to healthcare facilities, diagnostic tools, and resources. Therefore, an in-depth understanding of molecular mechanism of sepsis is great significance.

Sepsis is a complex problem thought to result from genetic [19] factors. However, there is no satisfactory treatment for sepsis. Therefore, there is an urgent need to better understand the molecular pathogenesis and genetic modulators of sepsis to develop effective therapies. Many studies focused on the identification of sepsis susceptibility genes and signaling pathways, which were correlated with the pathogenesis. Genes include APOL1 [20], serum mannose-binding lectin (MBL) [21], IL-6, IL-10 and IL-17 [22], NOS2 [23], and TLR4 and TNF-alpha (TNFA) [24] were responsible for sepsis. Signaling pathways include TLR4 signaling pathway [25], JNK1/Sirt1/FoxO3a signaling pathway [26], TGFBR2/Smad signaling pathway [27], CXCL12 signaling pathway [28] and sphingosine 1-phosphate receptor 2 signaling pathway [29] were linked with progression of sepsis. Nevertheless, the molecular mechanisms of sepsis remain to be fully understood.

Despite a many genes and signaling pathways in the advancement and progression of sepsis have been widely investigated, the molecular mechanisms underlying sepsis are still being unraveled. Currently, the next generation sequencing (NGS) data analysis of gene expression, coupled with bioinformatics tools, becomes promising for investigating the novel genes in the initiation and evolution of various diseases [30].

Bioinformatics analysis is a well-orchestrated tool for screening sepsis specific genes and prognosis-relevant biomarkers, which can contribute to the development of sepsis treatment [31]. At present investigation, NGS data (GSE185263) [32] downloaded from Gene Expression Omnibus (GEO) [https://www.ncbi.nlm.nih.gov/geo/] [33] database can be used to detect genes transcription expression levels and to provide the technical support for monitoring mRNA expression and cell function prediction. Afterwards, gene ontology (GO) terms and REACTOME pathways associated with DEGs were explored to elucidate the gene enrichment in sepsis. Moreover, protein–protein interaction (PPI) networks, modules and the hub genes were identified according to degree, betweenness, stress and closeness parameters. A miRNA-hub gene regulatory network and TF-hub gene regulatory network constructed to determine the miRNA and TFs were associated with sepsis. The expression levels of the identified hub genes were validated based on receiver operating characteristic (ROC) curve analysis. Our results may provide potential biomarker candidates for clinical diagnosis and therapy of sepsis.

## Materials and Methods

### Next generation sequencing data source

We downloaded the NGS dataset GSE185263 as blood immune expression profiles of patients with sepsis, deposited by Baghela et al [32] from the GEO database. The dataset was performed on GPL16791 Illumina HiSeq 2500 (Homo sapiens) expression NGS platform. The NGS data for GSE185263 contained 392 samples, including 348 sepsis samples and 44 normal control samples. No ethics approval and patients’ informed consent were needed for this current investigation.

### Identification of DEGs

DESeq2 package in R software [34] was adopted to identify the DEGs between sepsis and normal control. The log-fold change (FC) in expression and adjusted P-values (adj. P) were determined. The adj. P using the Benjamini-Hochberg method with default values was applied to correct the potential false-positive results [35]. Genes that met the specific cut-off criteria of adj. P<0.05, and |logFC|> 1.3175 were regarded as up regulated genes and |logFC|< −2.137 were regarded as down regulated genes. The ggplot2 and gplot packages in R software was used to generate volcano plots and heatmaps.

### GO and pathway enrichment analyses of DEGs

To comprehensively investigate the biological meaning behind DEGs intersection, g:Profiler (http://biit.cs.ut.ee/gprofiler/) [36] was used to obtain the set of functional annotation. GO (http://www.geneontology.org) [37] is an extensively used tool for annotating genes with potential functions, such as biological pathways (BP), cellular components (CC) and molecular function (MF). REACTOME pathway (https://reactome.org/) [38] enrichment analysis is a practical resource for analytical study of gene functions and associated high-level genome functional information. p < 0.05 was regarded as the threshold value.

### Construction of the PPI network and module analysis

The Search Tool for the Retrieval of Interacting Genes (STRING) (http://string-db.org) [39] is a public database resource designed to analyze PPI information. In the current investigation, the STRING online tool was utilized to generate the full STRING network containing all DEGs. Subsequently, Cytoscape software (V3.10.0; http://cytoscape.org/) [40] was used to visualize and analyze biological networks. In this experiment, the genes with the highest degree [41], betweenness [42], stress [43] and closeness [44] values were considered as hub genes. Finally, the Cytoscape plug-in PEWCC [45] was used to identify the key modules from PPI network.

### Construction of the miRNA-hub gene regulatory network

In the disease state, gene expression is affected by miRNA through the posttranscriptional control. In this investigation, the miRNet database (https://www.mirnet.ca/) [46] is used to search for miRNAs related to hub genes. miRNet database is a publicly available comprehensive resource, a comprehensive miRNA-hub gene database, including predicted and experimentally verified miRNA-hub gene interaction pairs. The miRNA binding sites on the full-length sequence of the gene are not only recorded but also compared with the prediction binding information collection of the existing 14 miRNA-hub gene prediction programs (TarBase, miRTarBase, miRecords, miRanda (S mansoni only), miR2Disease, HMDD, PhenomiR, SM2miR, PharmacomiR, EpimiR, starBase, TransmiR, ADmiRE, and TAM 2.0). Subsequently, Cytoscape software (V3.10.0; http://cytoscape.org/) [40] was used to visualize and analyze biological networks.

### Construction of the TF-hub gene regulatory network

In the disease state, gene expression is affected by TF through the posttranscriptional control. In this investigation, the NetworkAnalyst database (https://www.networkanalyst.ca/) [47] is used to search for TFs related to hub genes. NetworkAnalyst database is a publicly available comprehensive resource, a comprehensive TF-hub gene database, including predicted and experimentally verified TF-hub gene interaction pairs. The TF binding sites on the full-length sequence of the gene are not only recorded but also compared with the prediction binding information collection of the existing ChEA. Subsequently, Cytoscape software (V3.10.0; http://cytoscape.org/) [40] was used to visualize and analyze biological networks.

### Receiver operating characteristic curve (ROC) analysis

ROC analysis was performed to predict the diagnostic effectiveness of biomarkers by pROC package in R statistical software [48]. The area under the ROC curve (AUC) value was utilized to determine the diagnostic effectiveness in discriminating sepsis from normal control samples. The higher the values of AUC, the greater the diagnostic efficiency of hub genes.

## Results

### Identification of DEGs

We identified 958 DEGs in sepsis patients, with a total of 479 up regulated genes and 479 down regulated genes compared with healthy controls (Table 1). We drafted a volcano plot of DEGs (Fig. 1) and a heat map of the differentially expressed genes (Fig. 2).

**Fig. 1.**
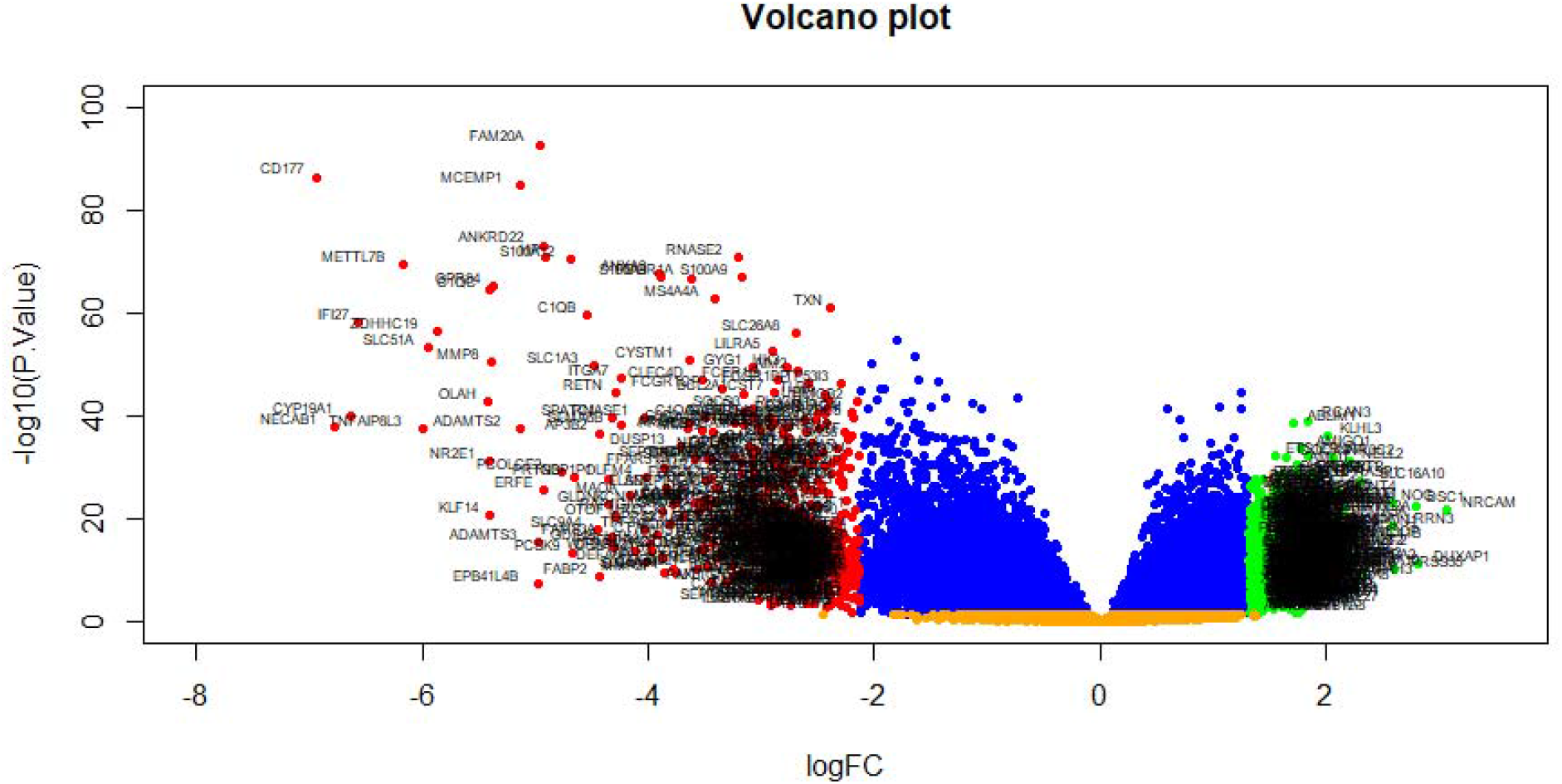
Volcano plot of differentially expressed genes. Genes with a significant change of more than two-fold were selected. Green dot represented up regulated significant genes and red dot represented down regulated significant genes.

**Fig. 2.**
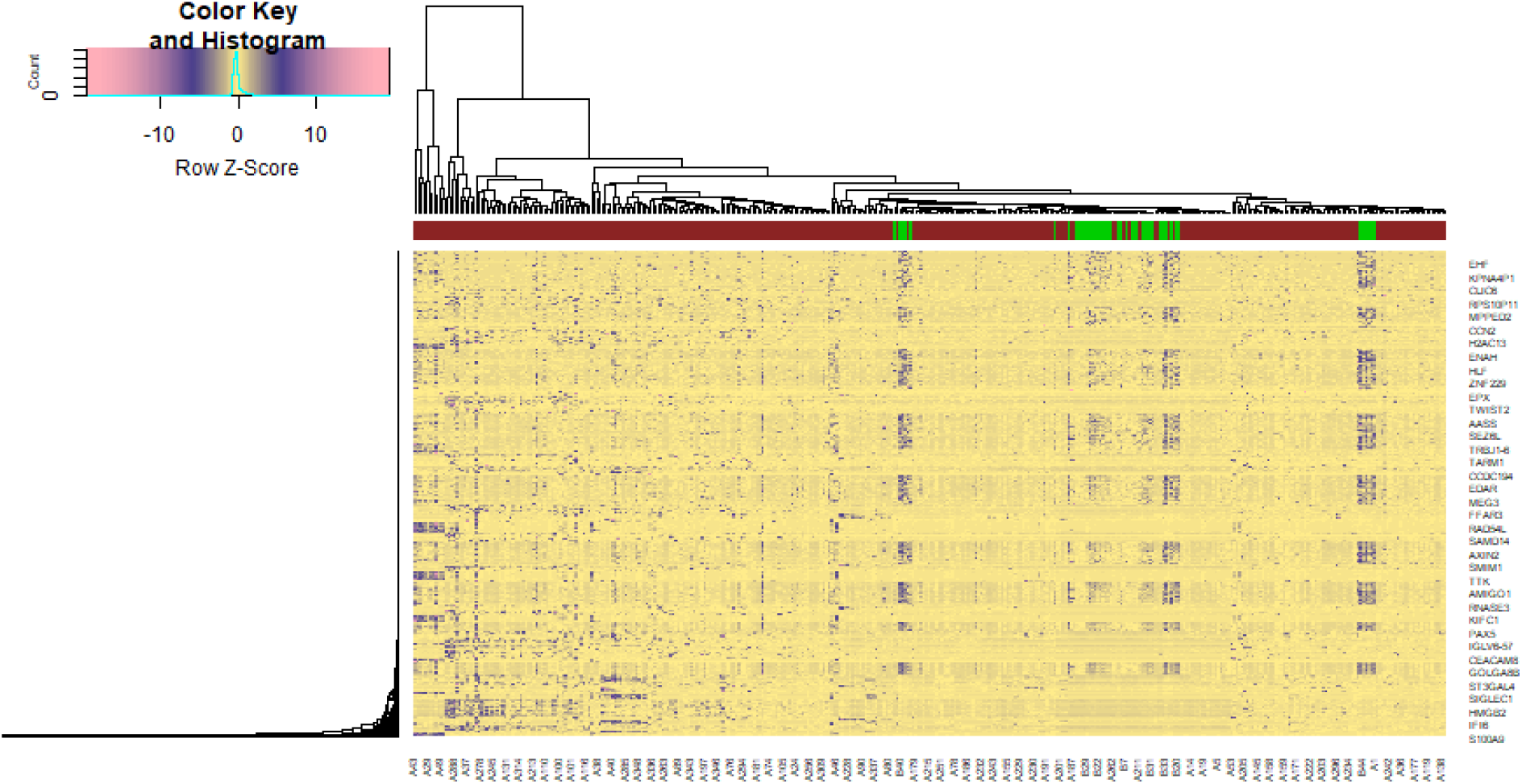
Heat map of differentially expressed genes. Legend on the top left indicate log fold change of genes. (A1 – A44 = normal control samples; B1 – B 348 = sepsis samples)

**Table 1.**
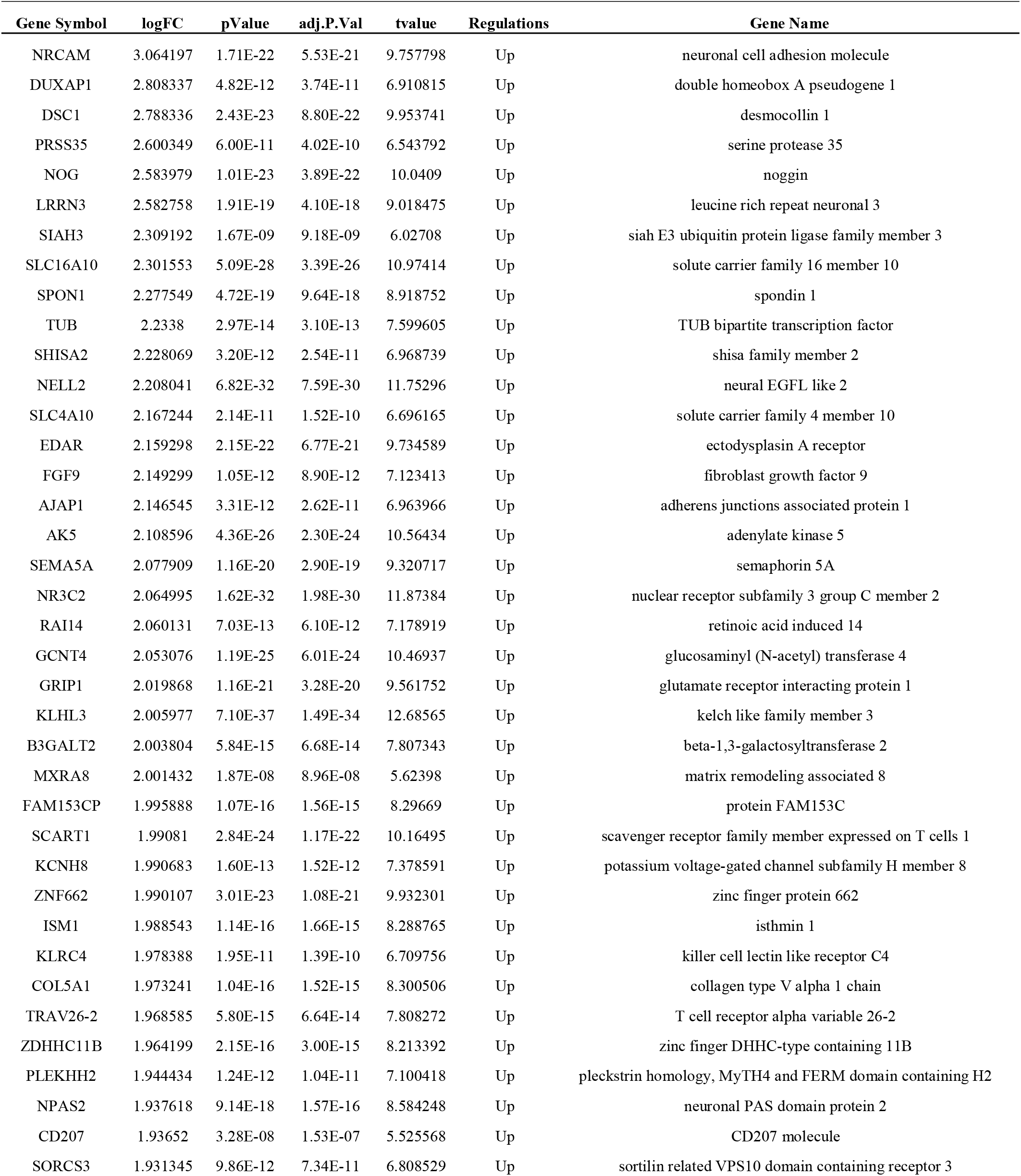
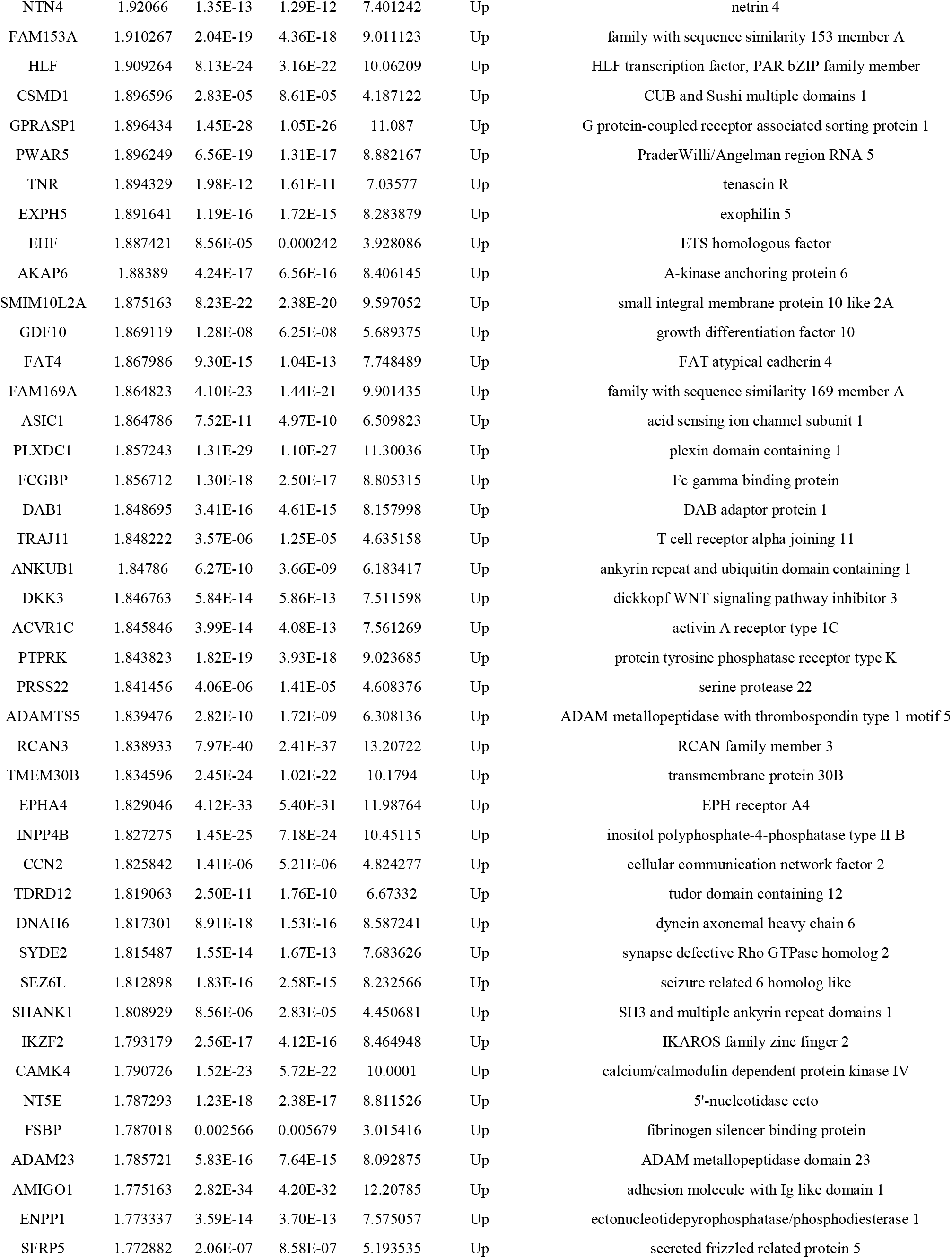
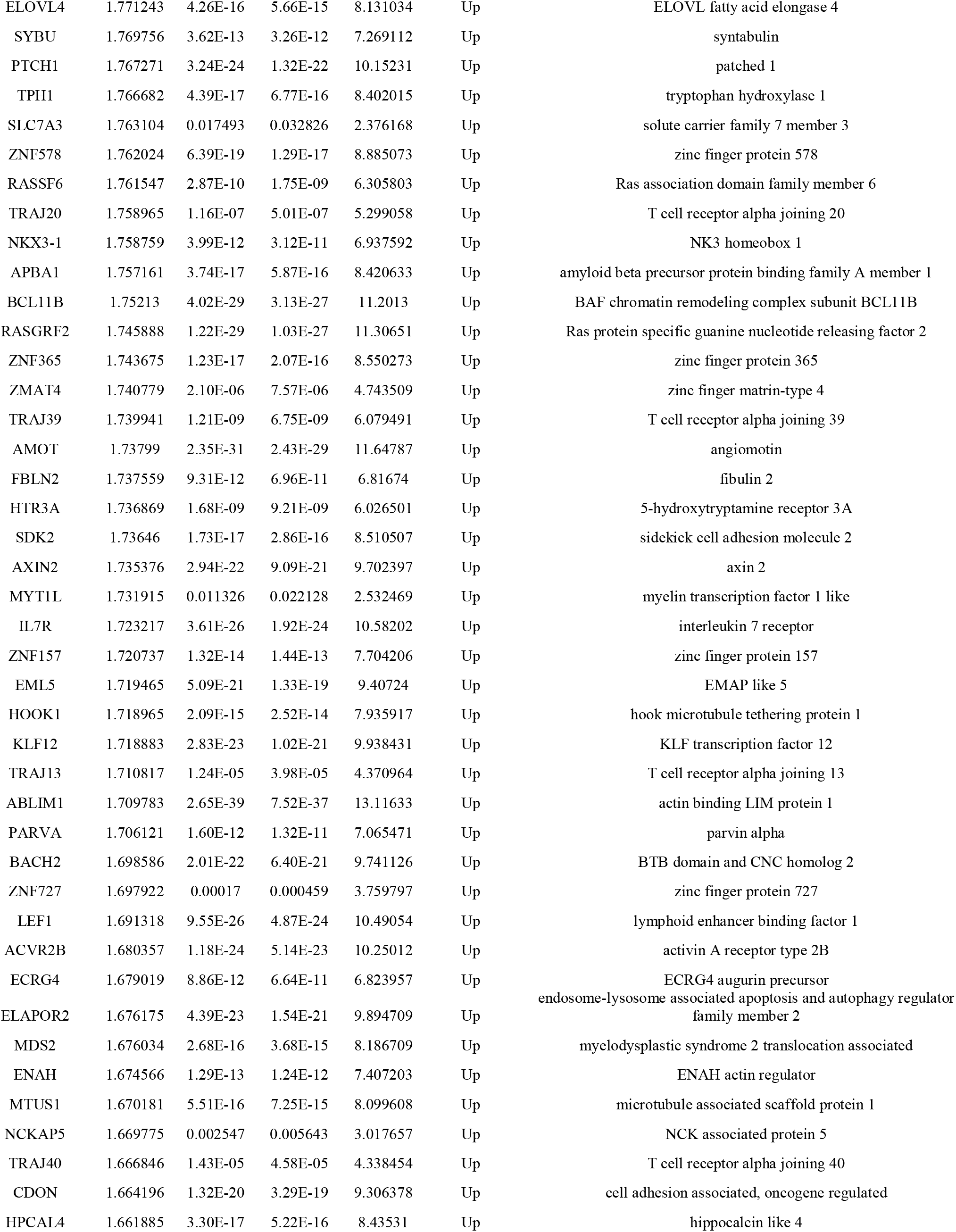
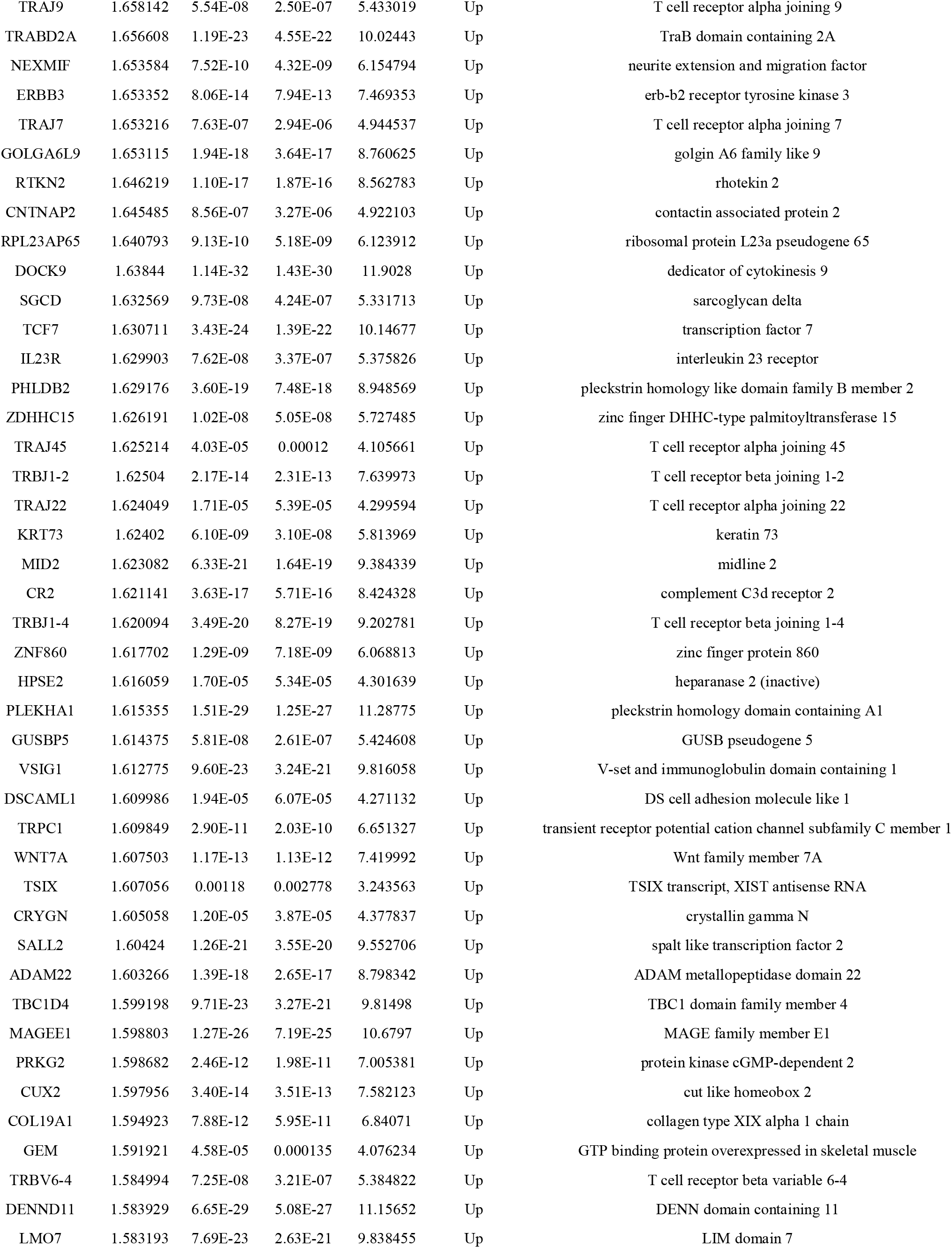
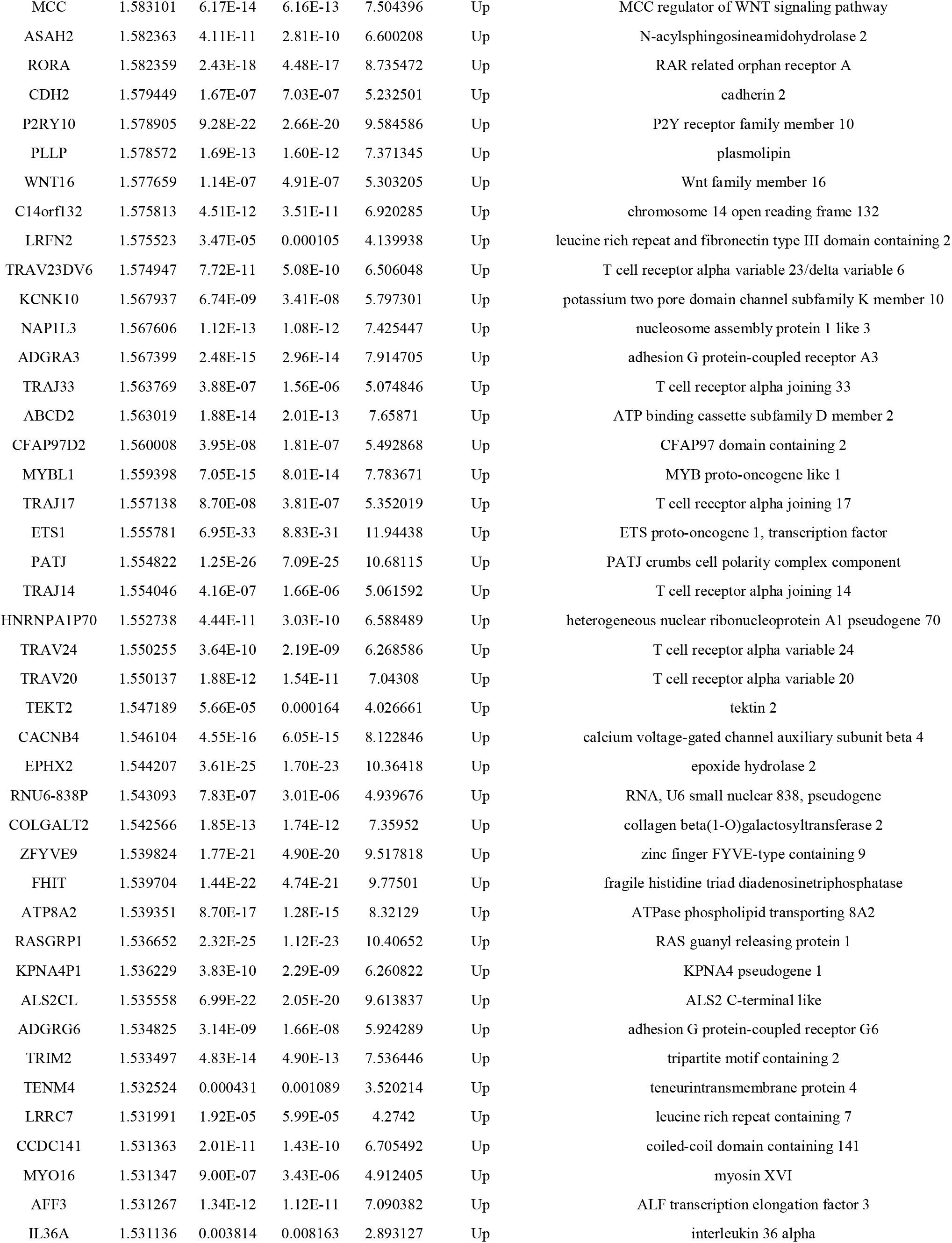
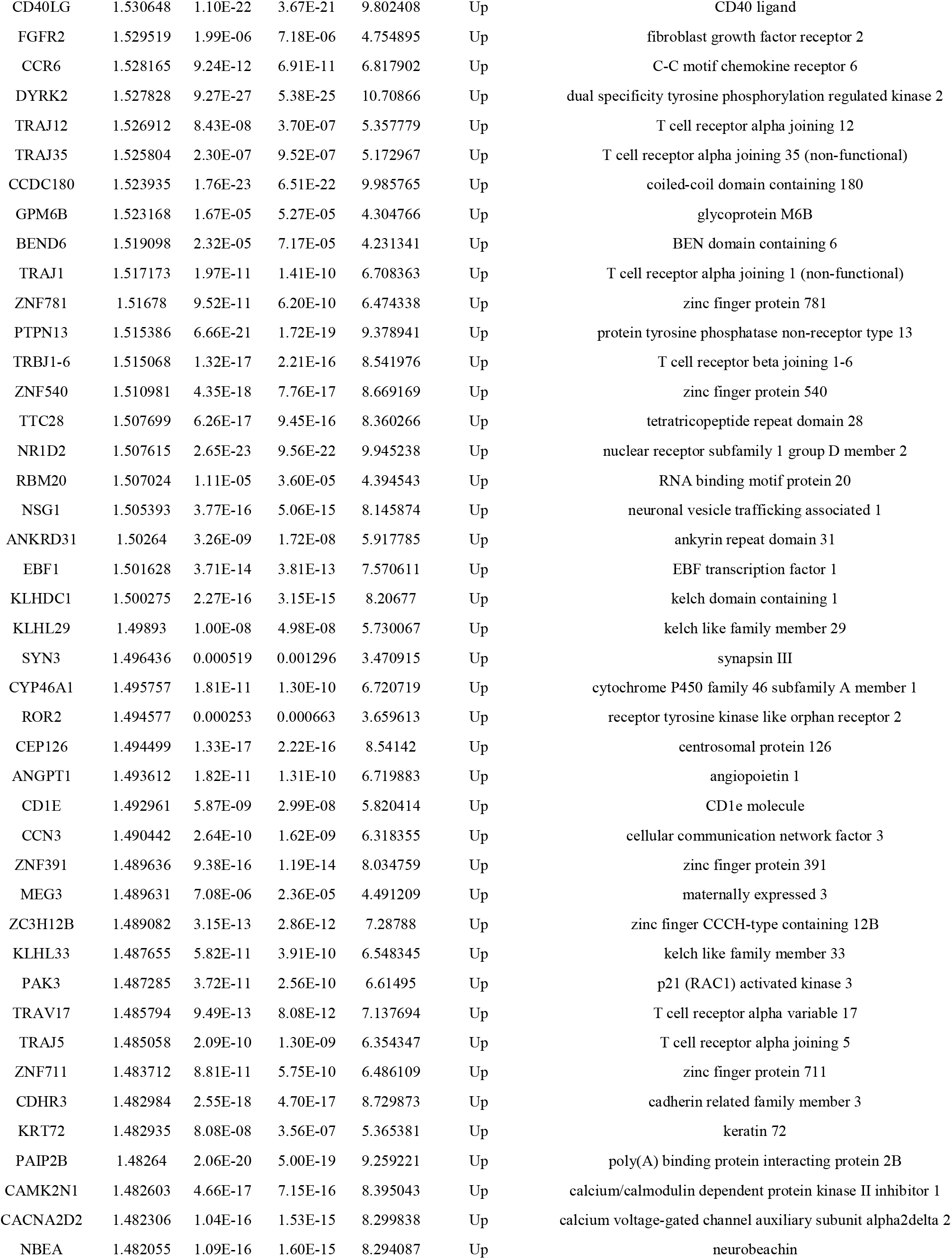
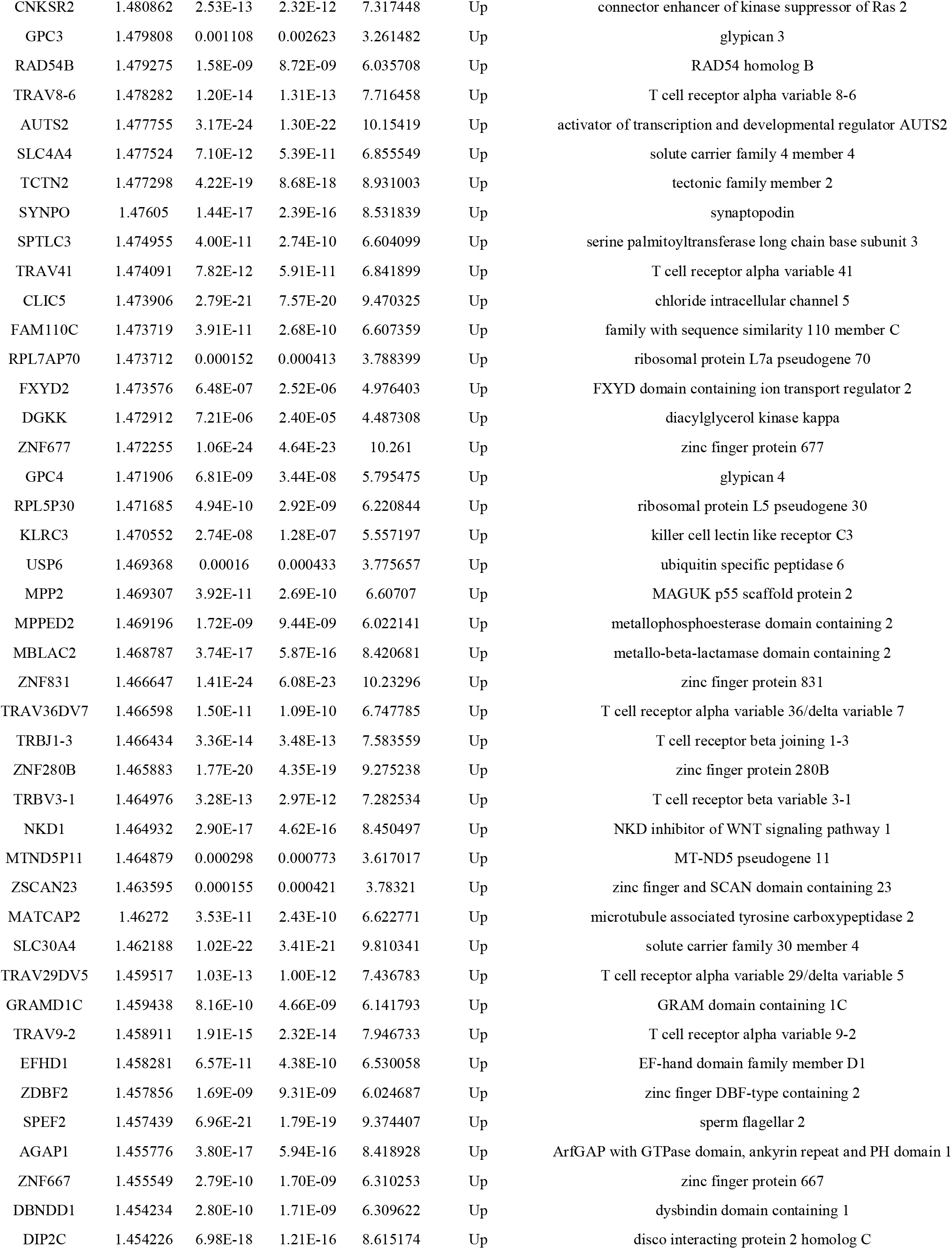
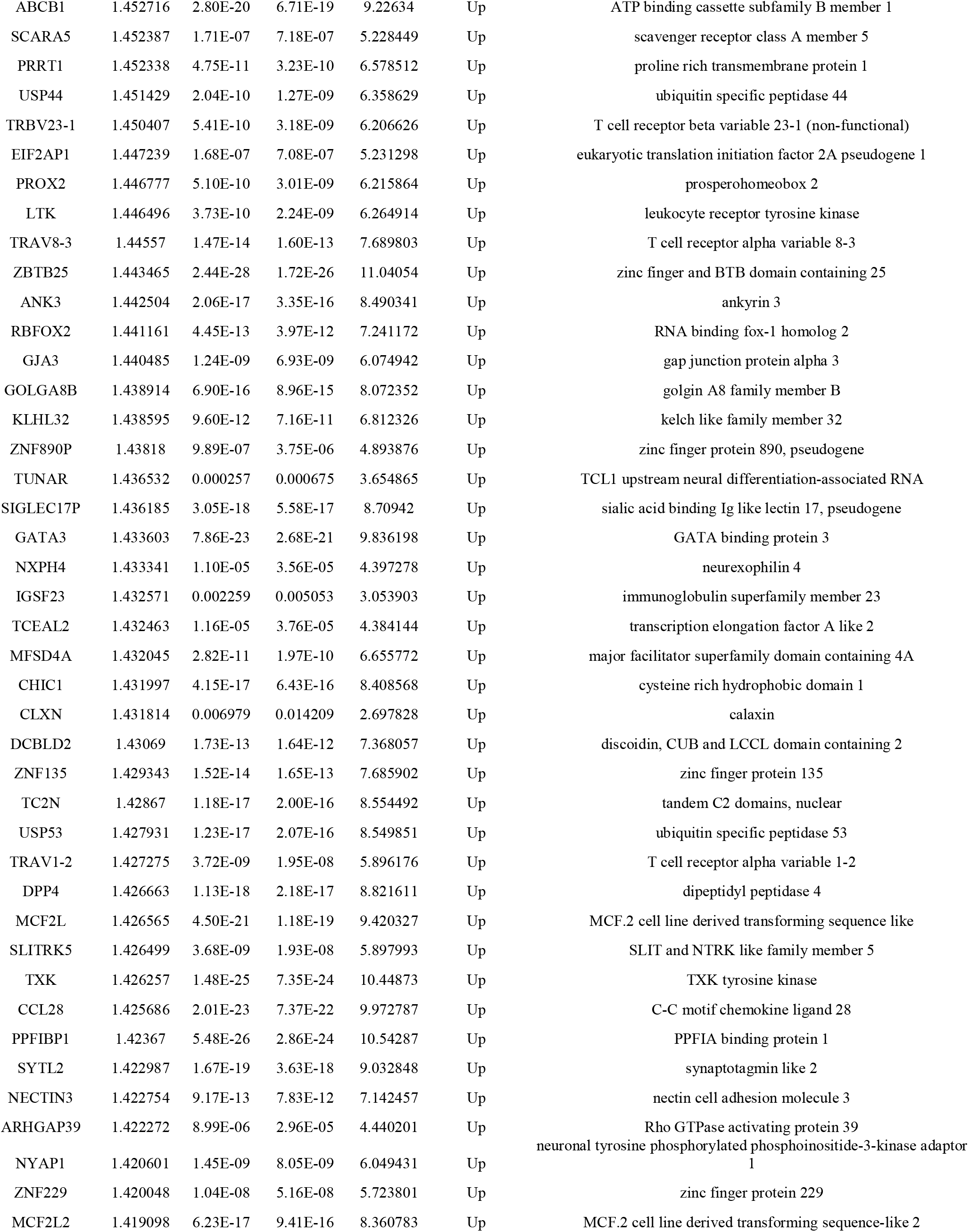
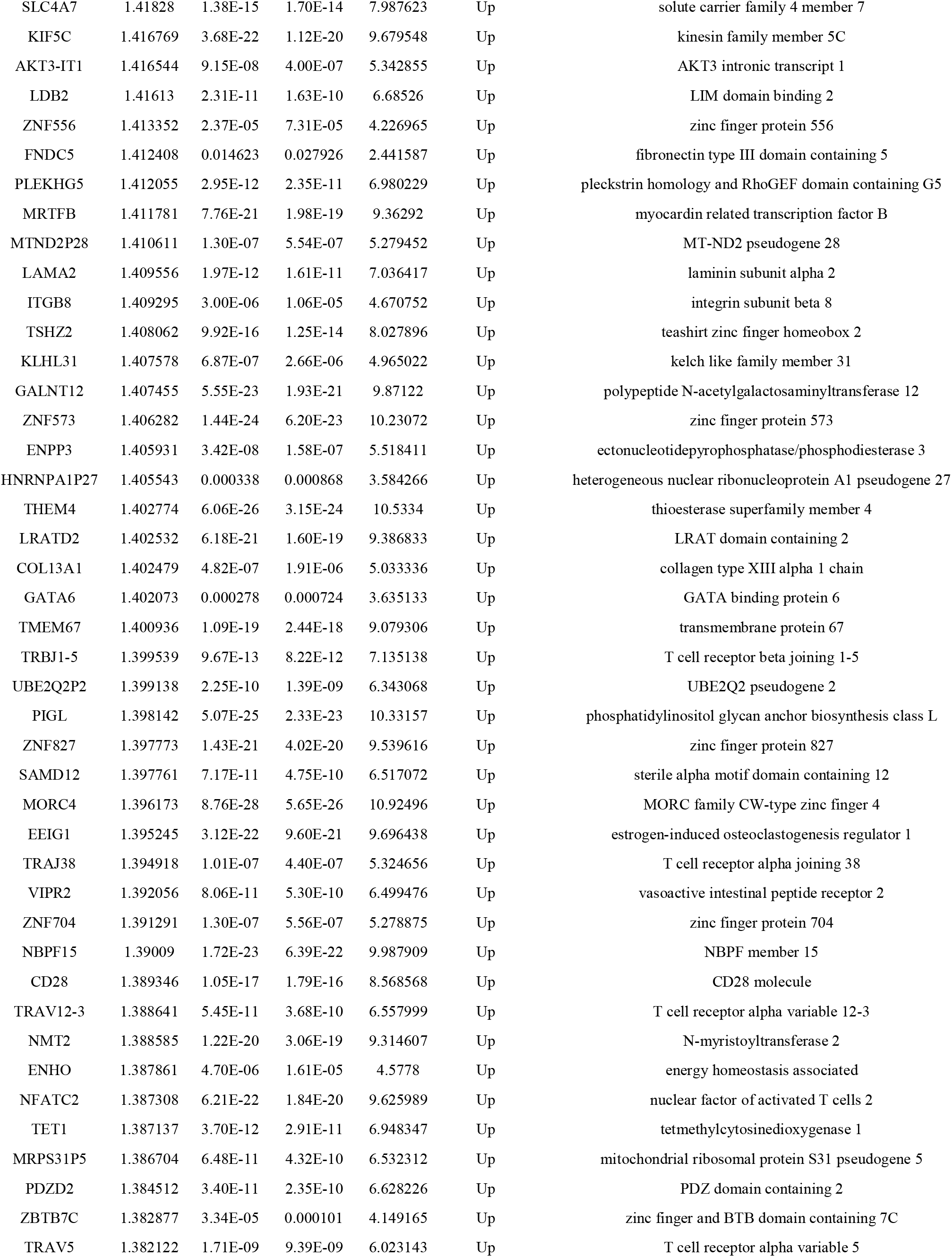
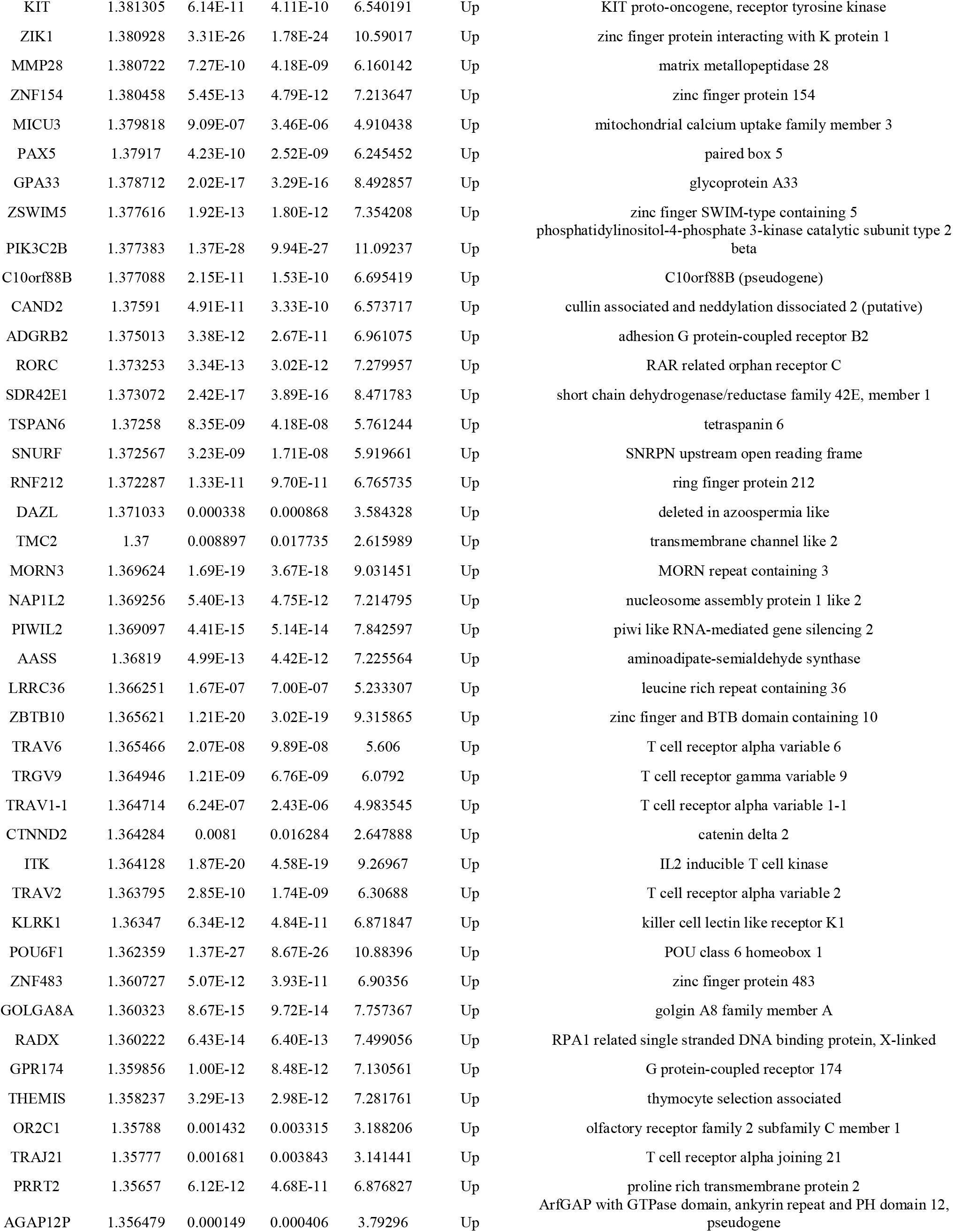
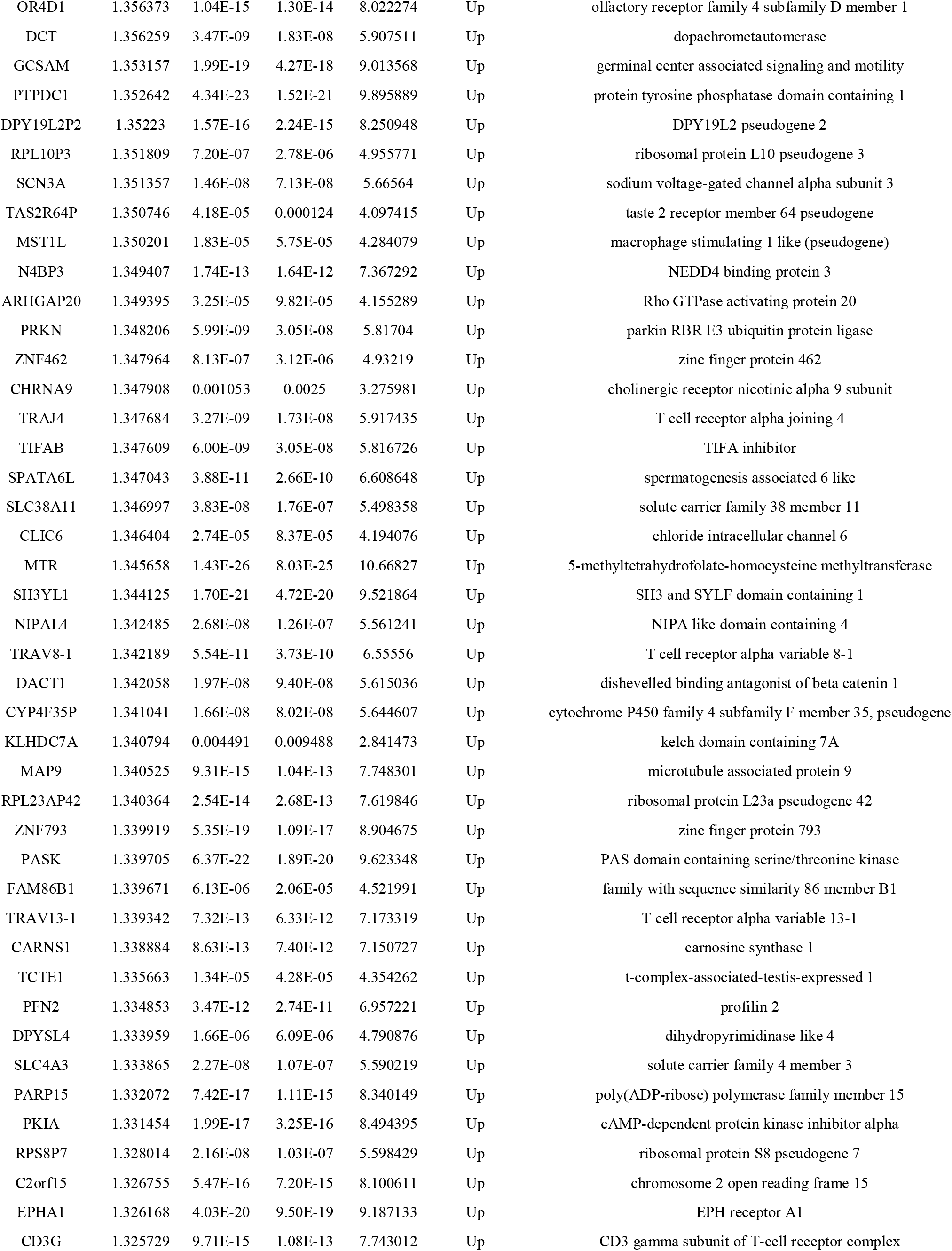
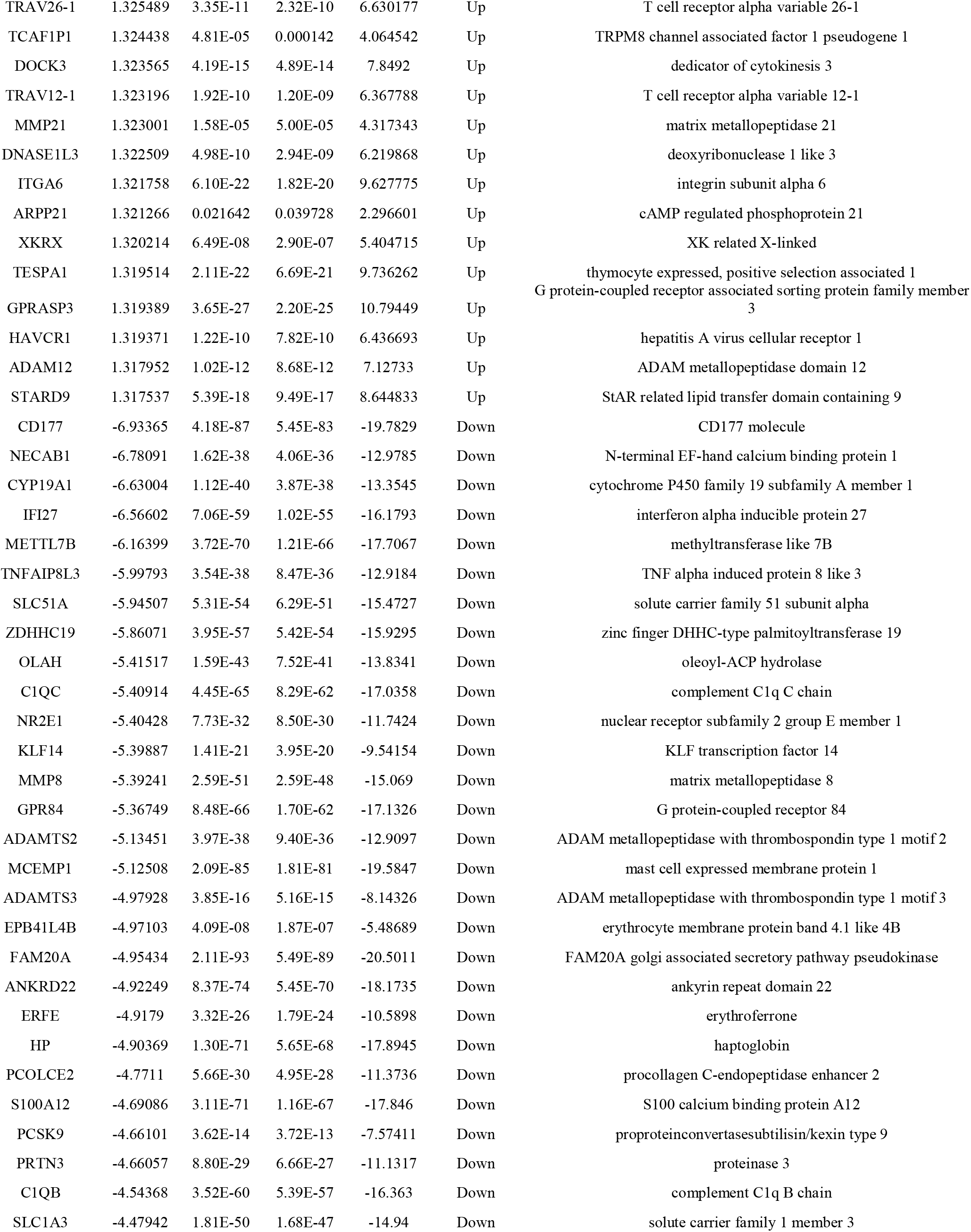
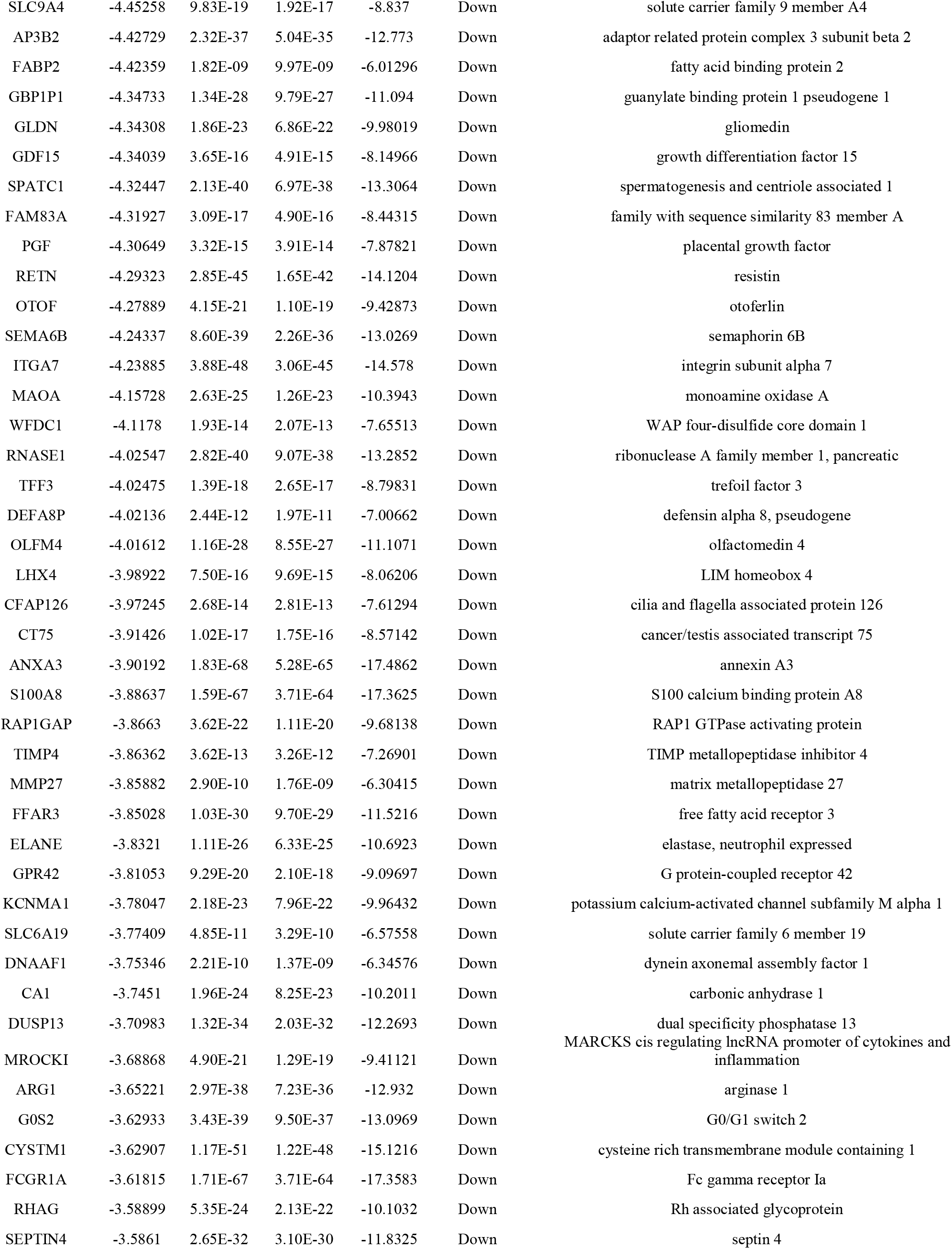
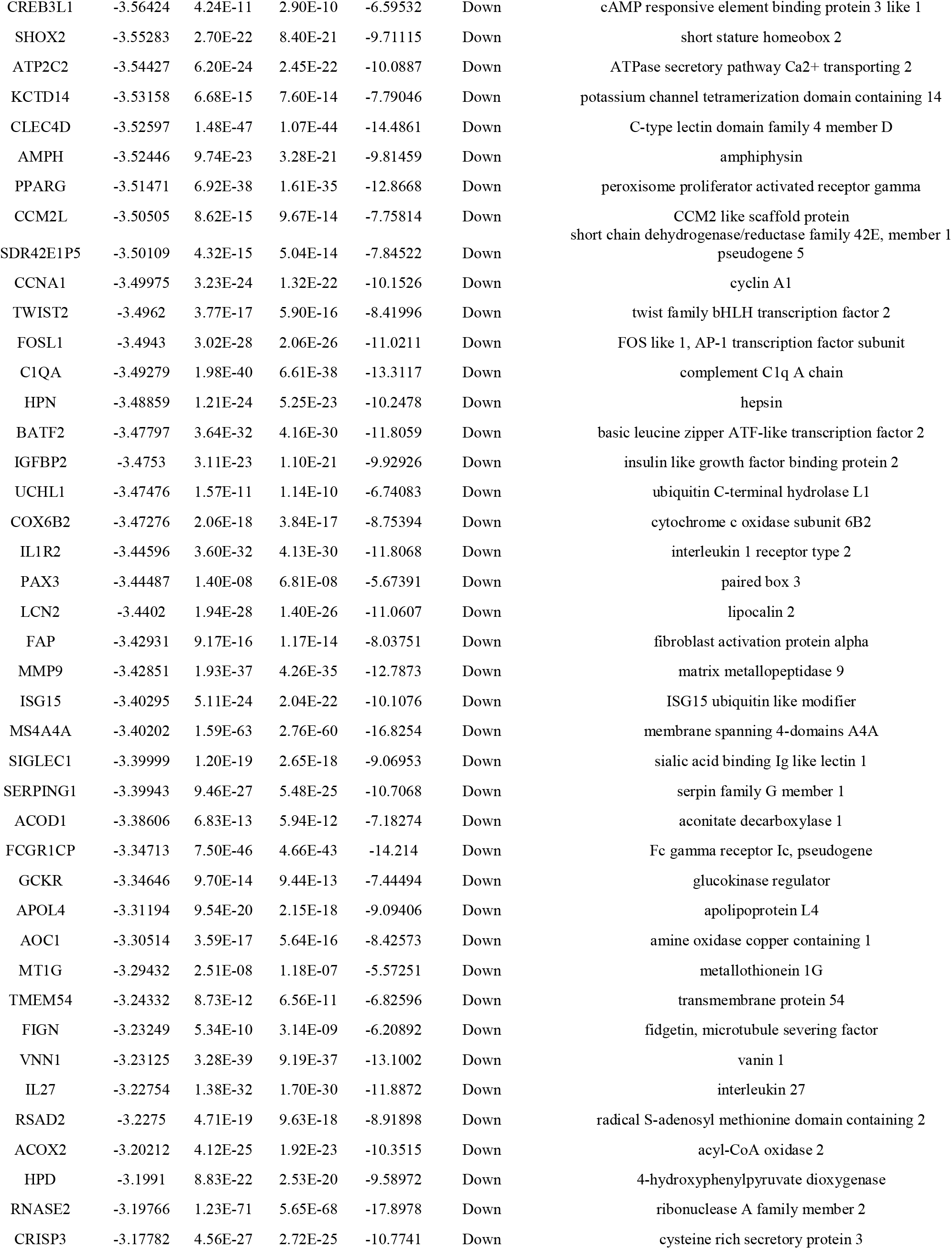
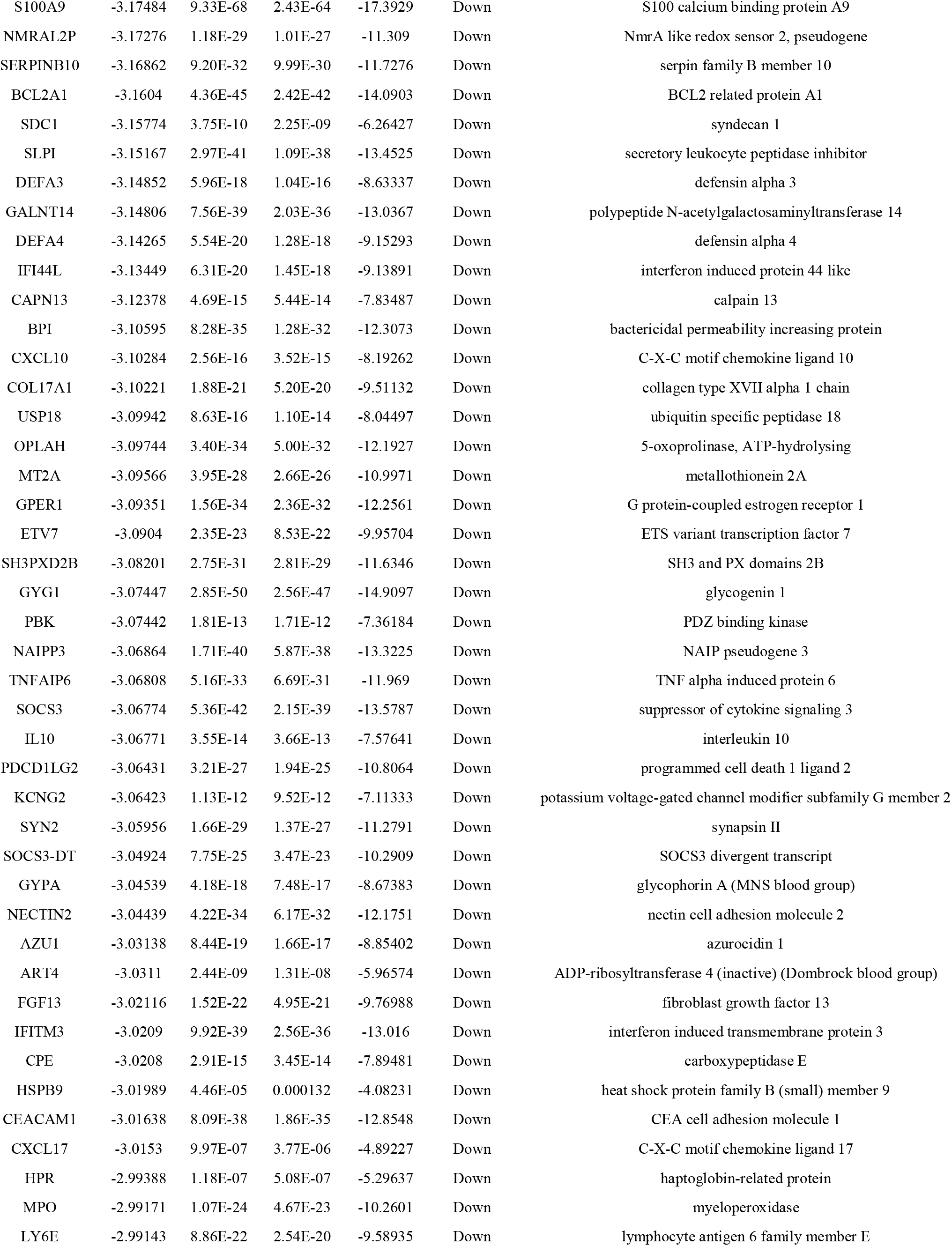
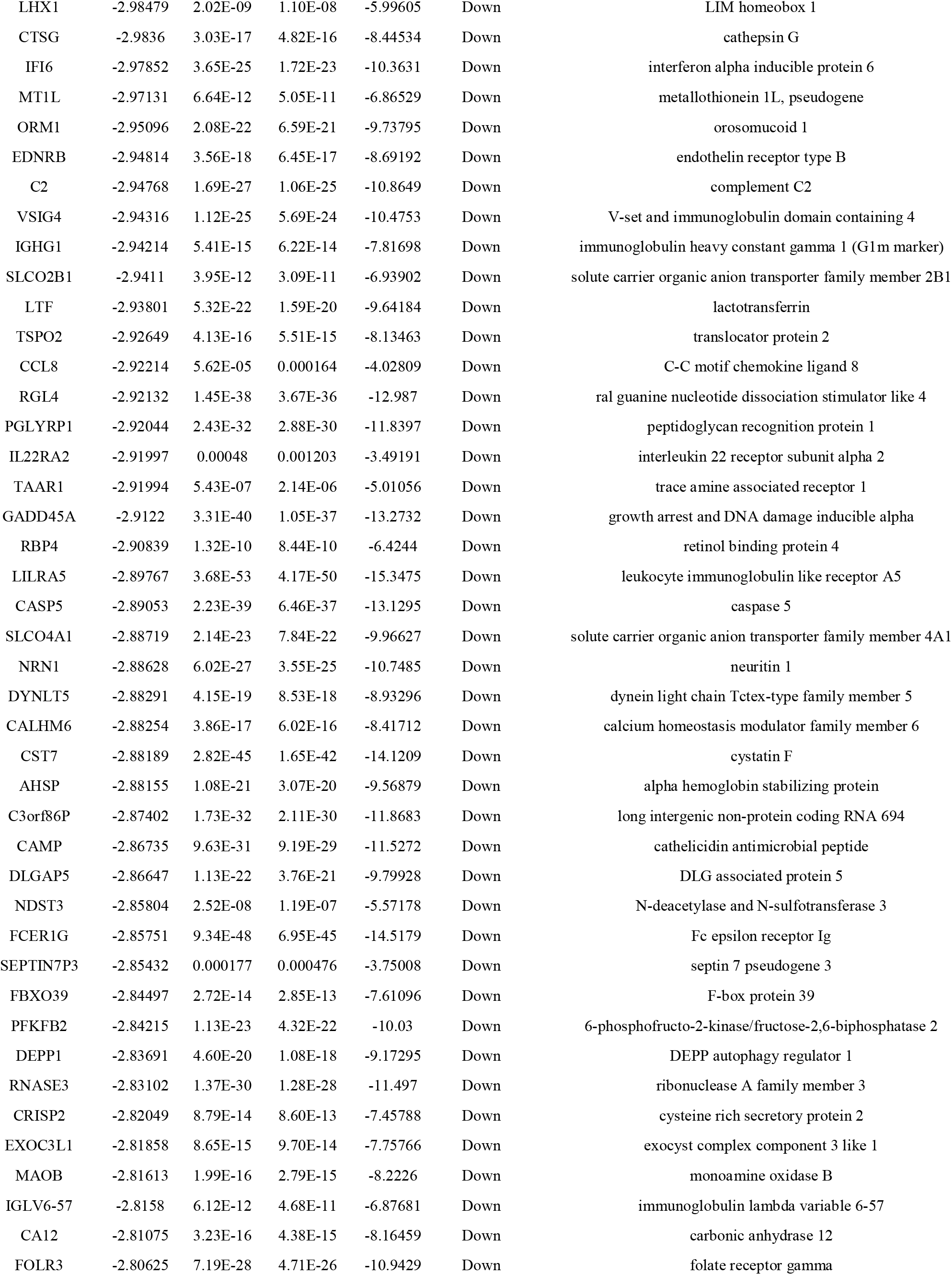
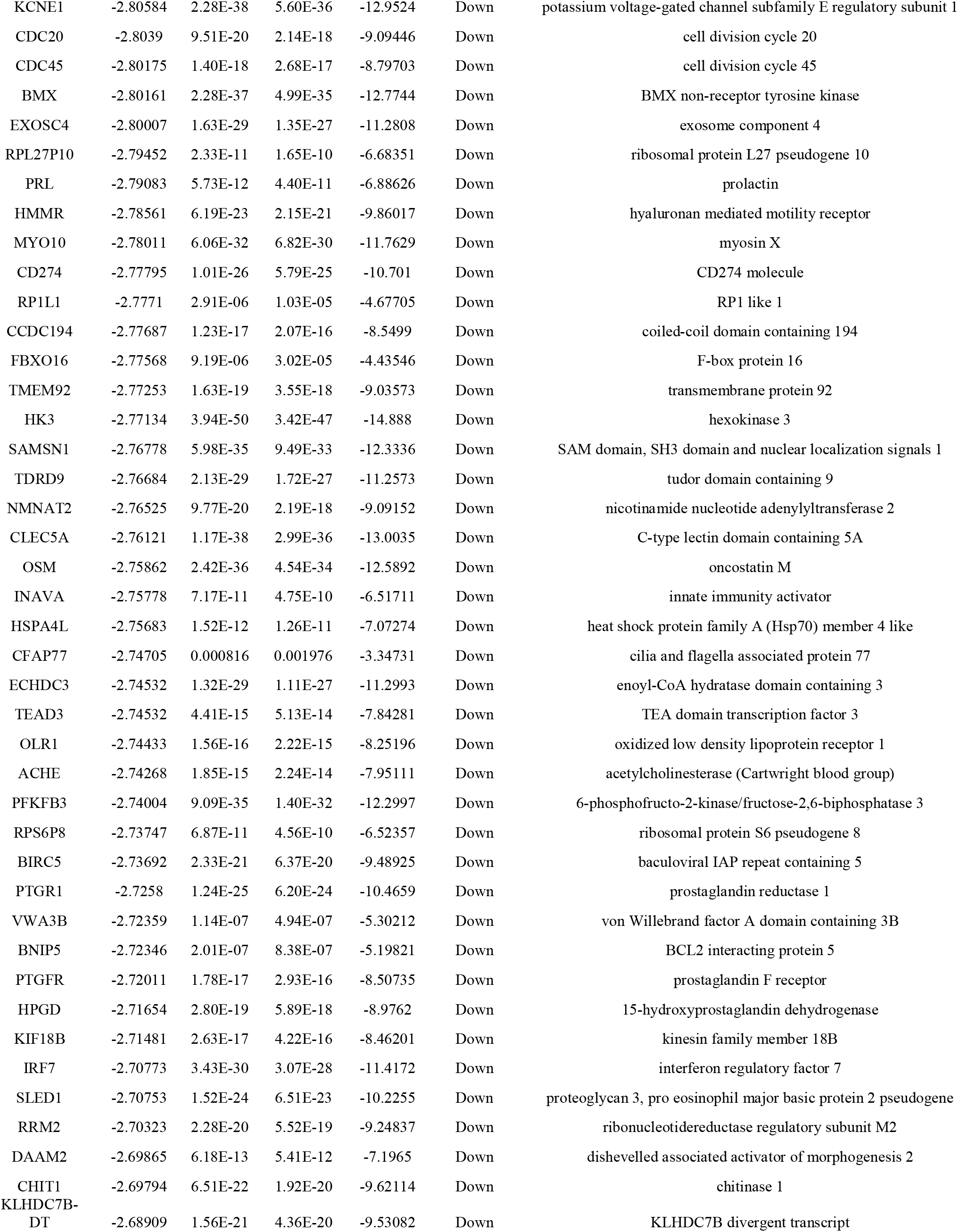
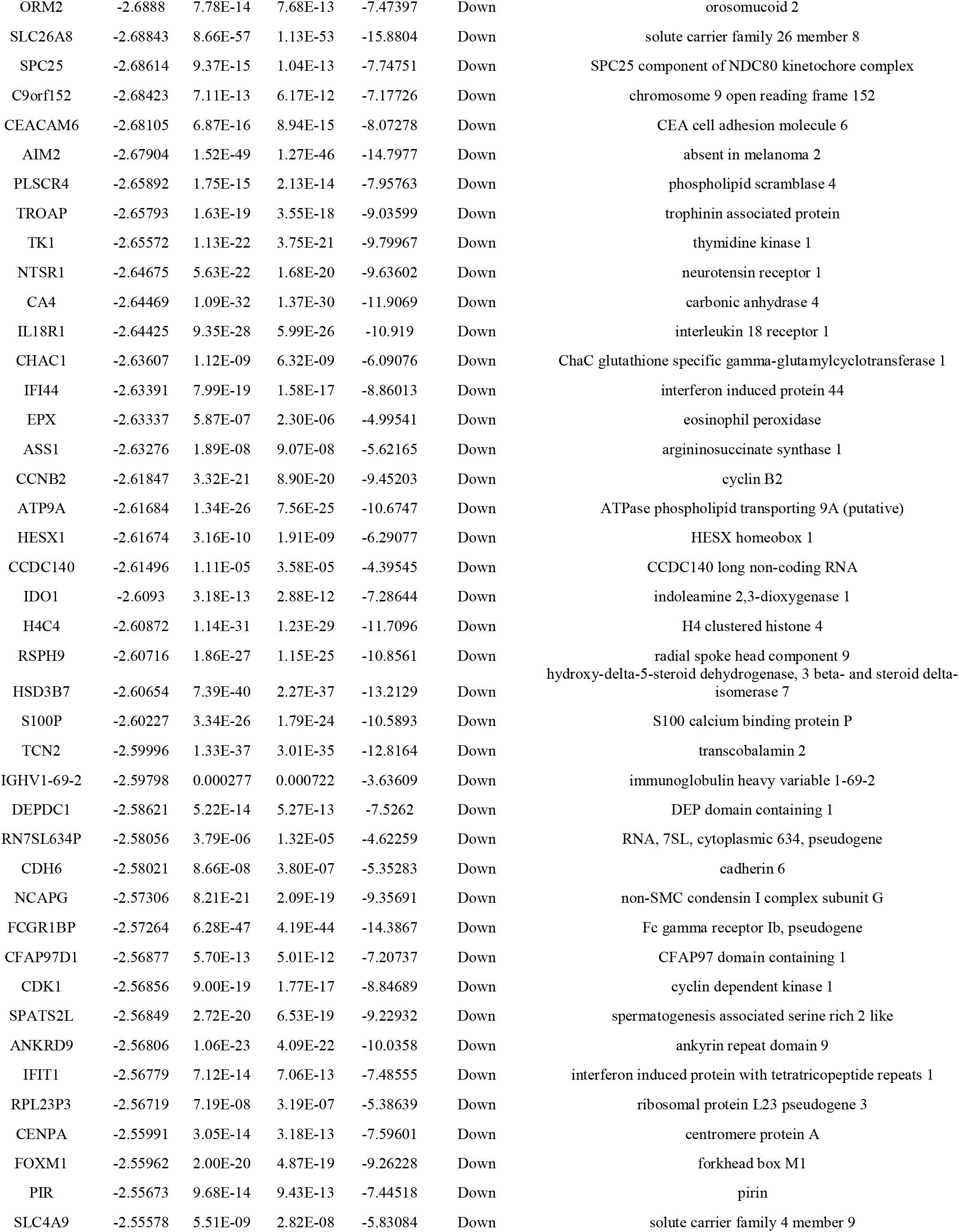
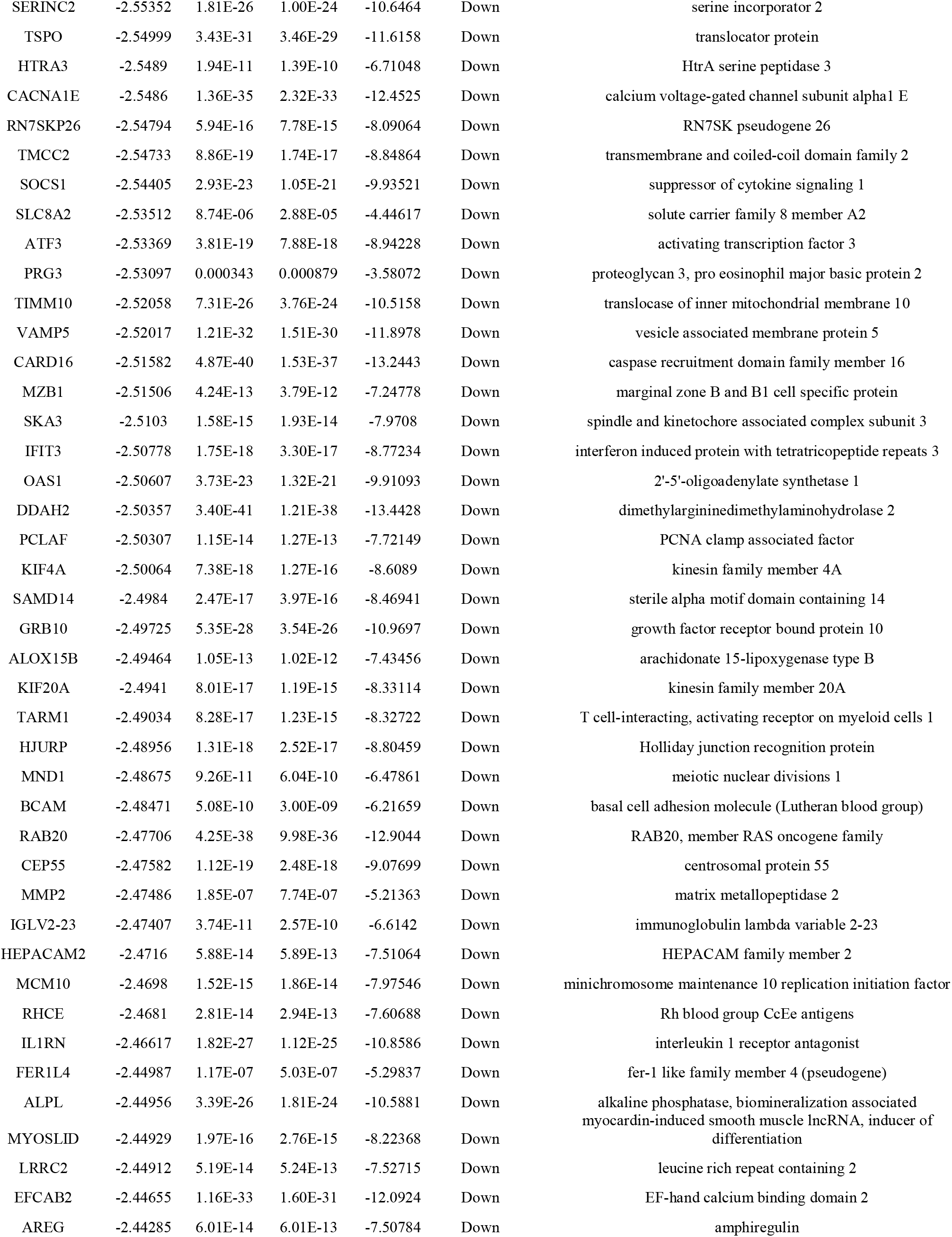
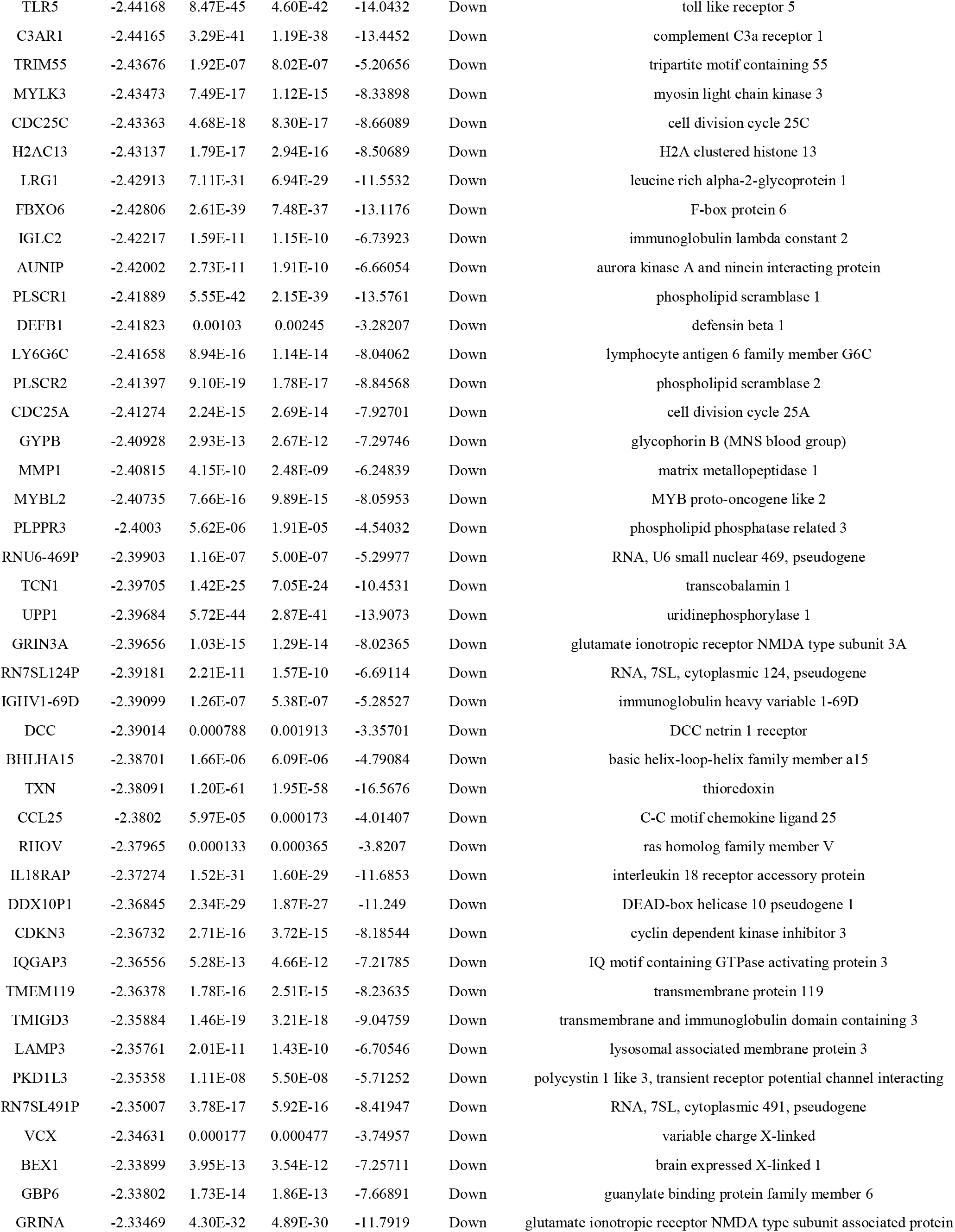
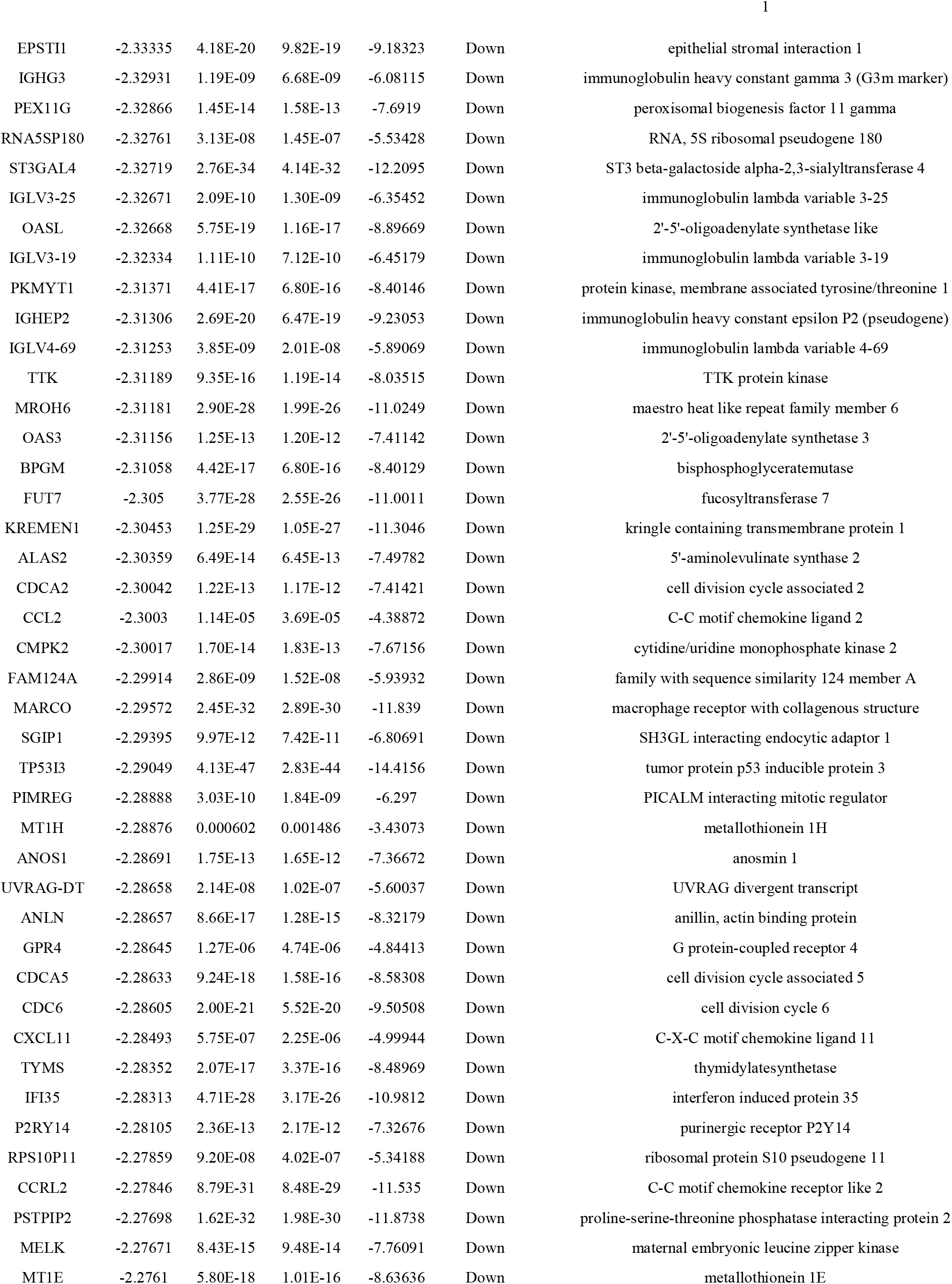
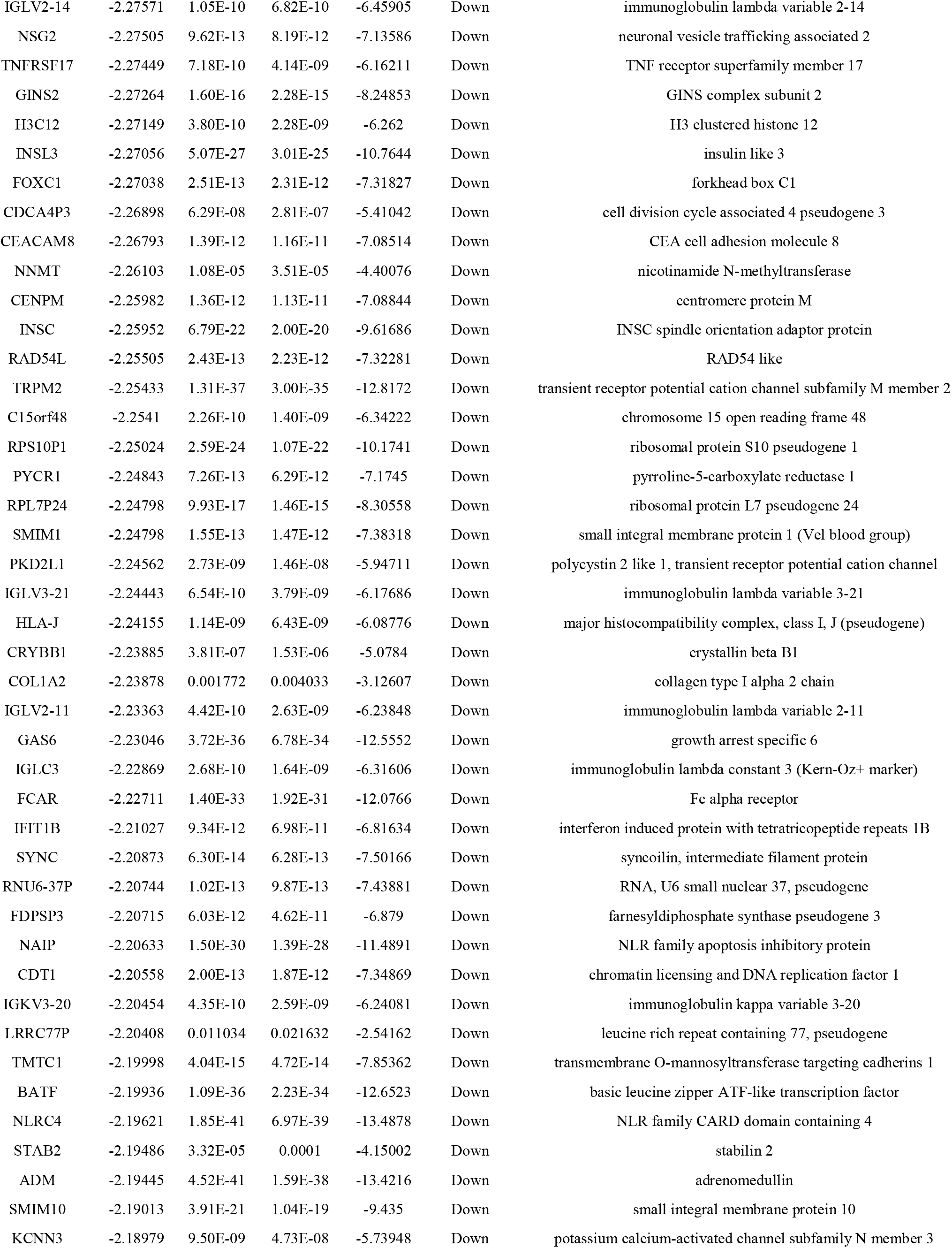
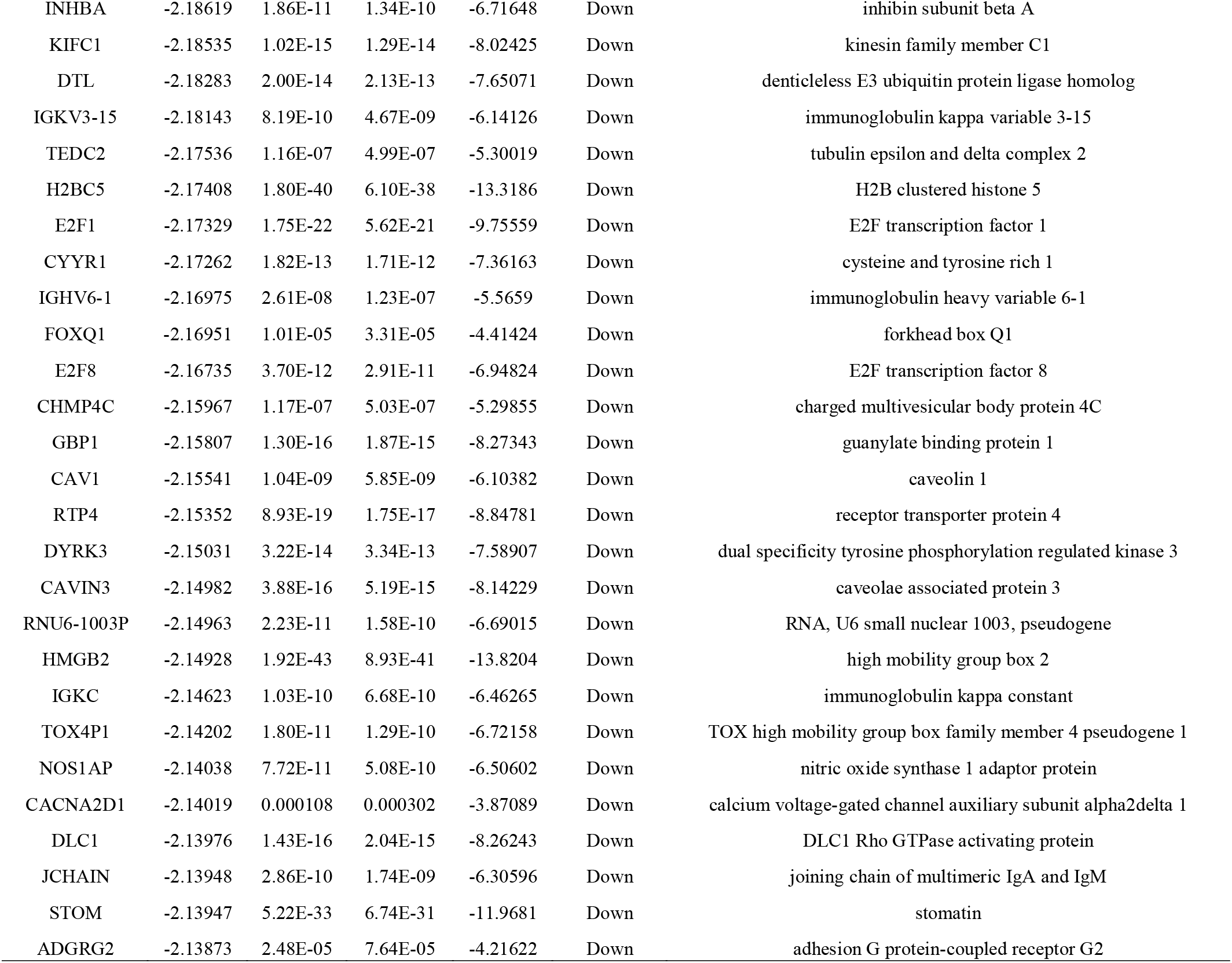
The statistical metrics for key differentially expressed genes (DEGs)

### GO and pathway enrichment analyses of DEGs

GO and REACTOME enrichment analysis of genes between sepsis and normal control groups were performed. In GO BP analysis, genes were primarily rich in regulation of cellular process, regulation of biological process, response to stimulus and signaling (Table 2). In GO CC analysis, genes were mainly rich in cell periphery, membrane, extracellular region and cytoplasm (Table 2). In GO MF analysis, genes were mainly rich in metal ion binding, signaling receptor activity, identical protein binding and signaling receptor binding (Table 2). In REACTOME pathway enrichment analysis, genes were rich in extracellular matrix organization, developmental biology, immune system and neutrophil degranulation (Table 3).

**Table 2.**
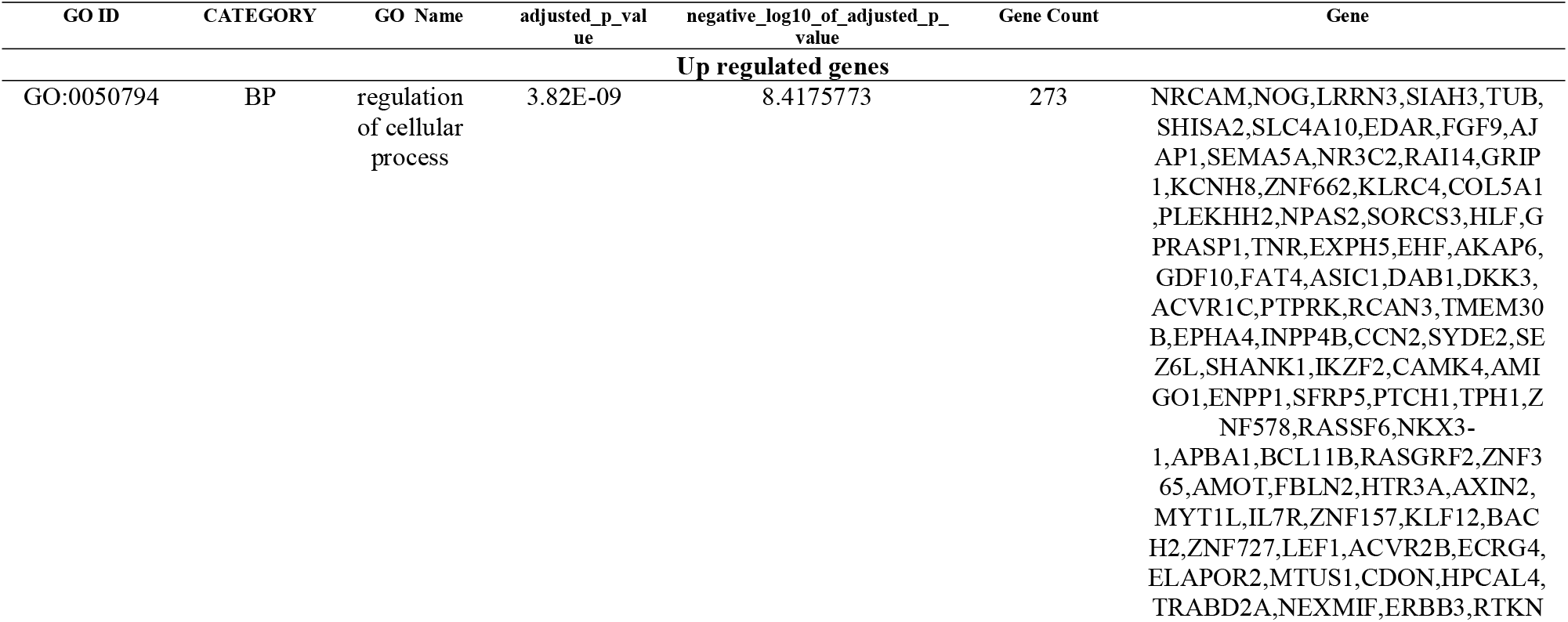
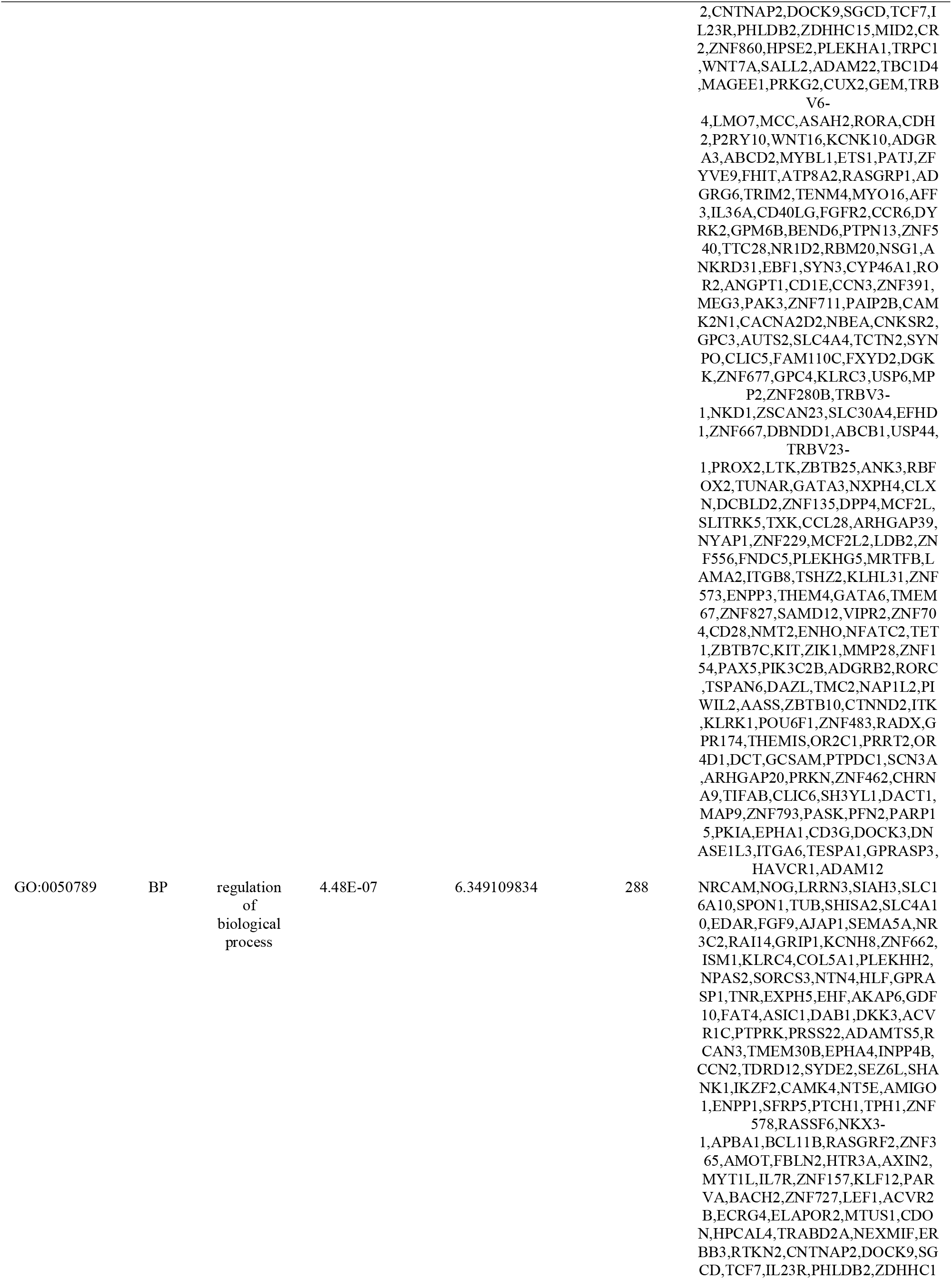
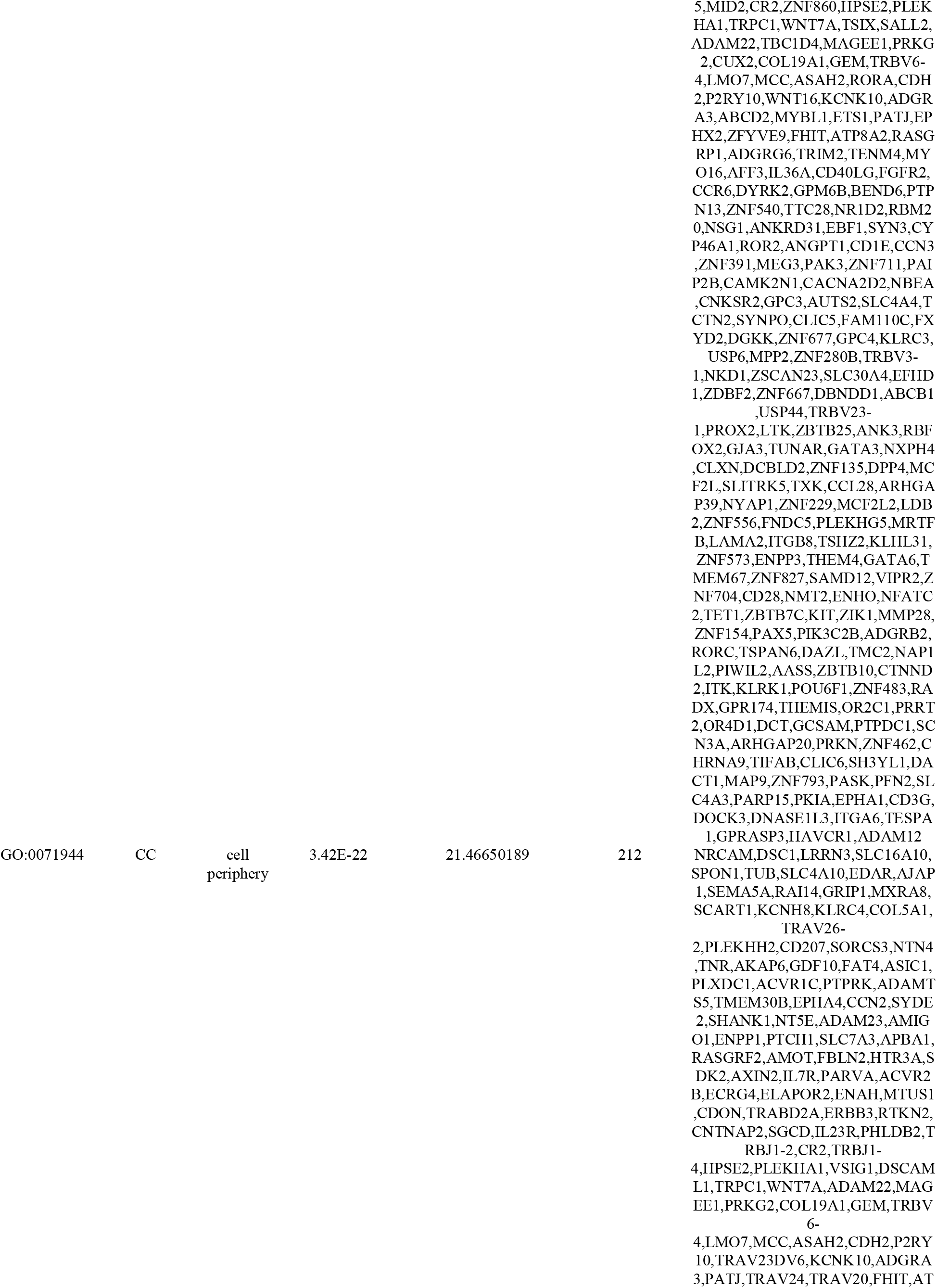
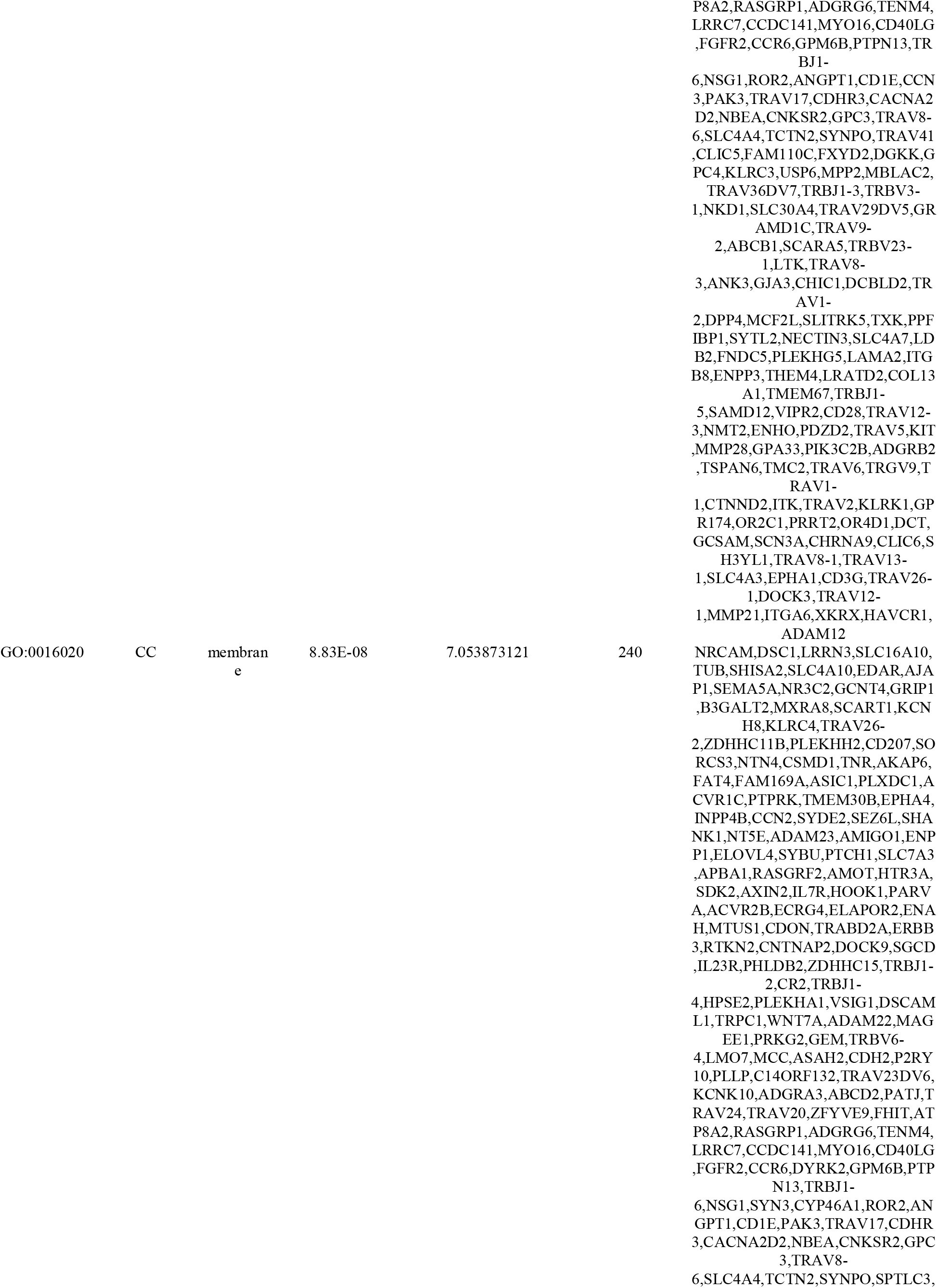
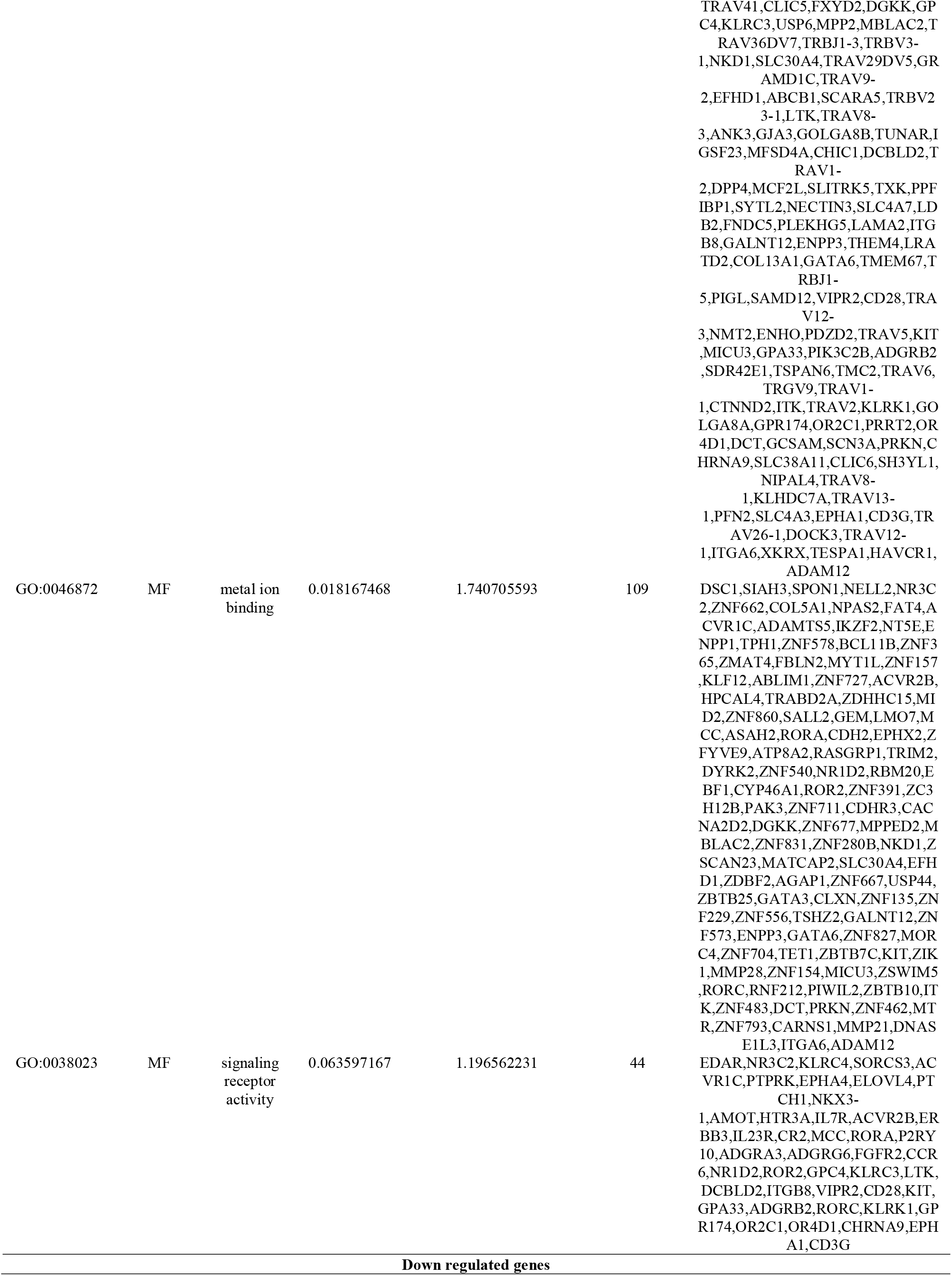
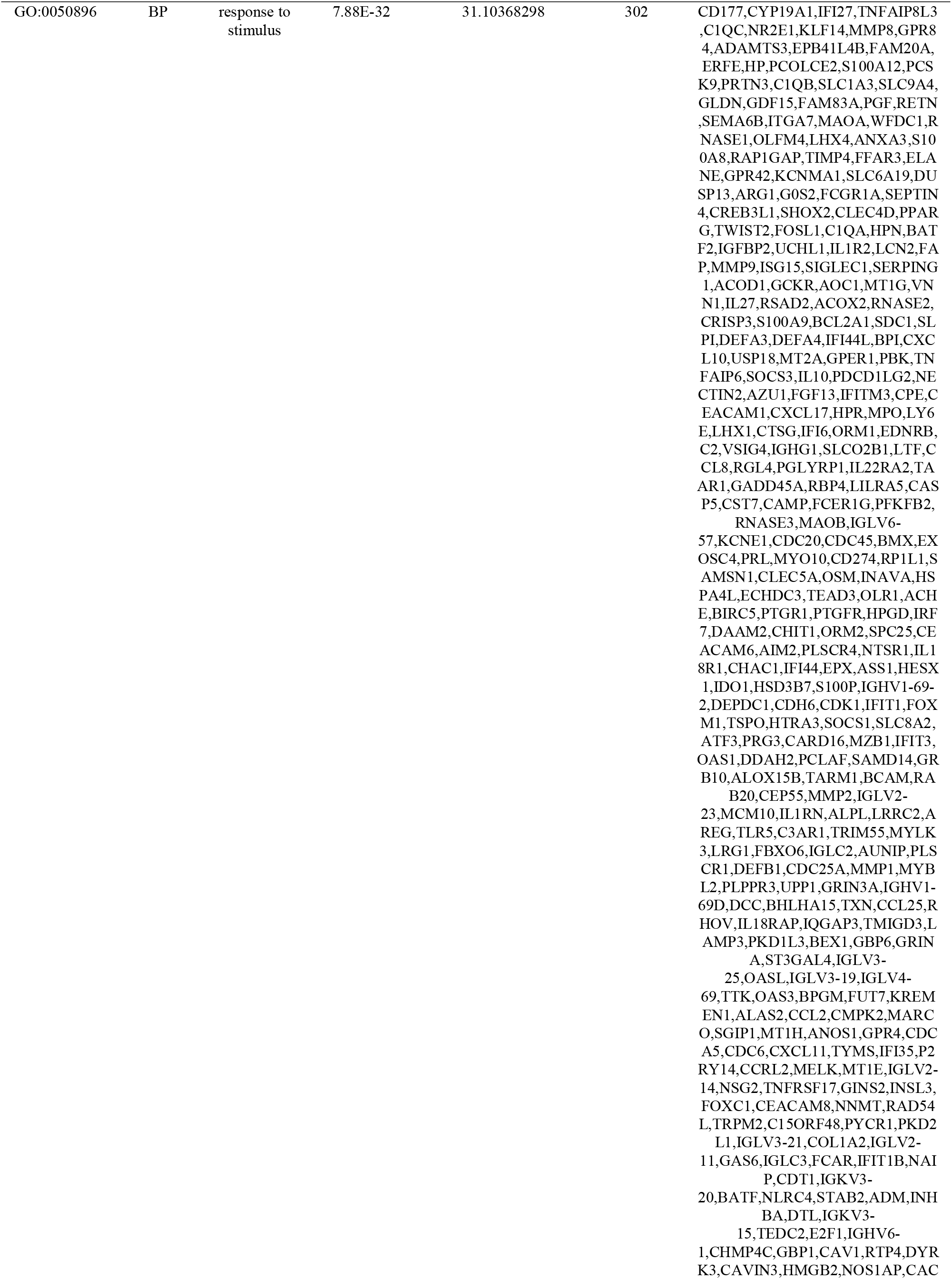
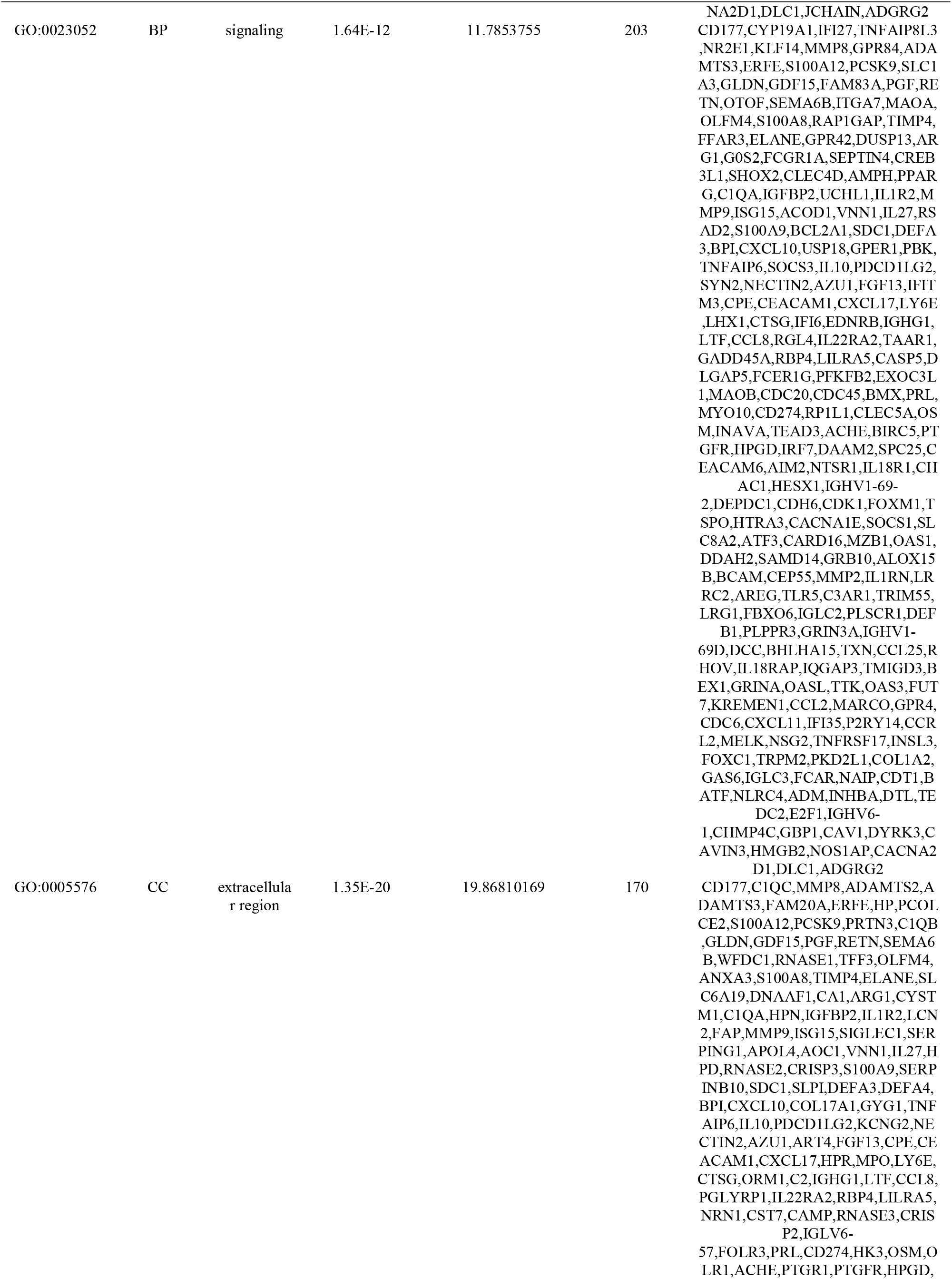
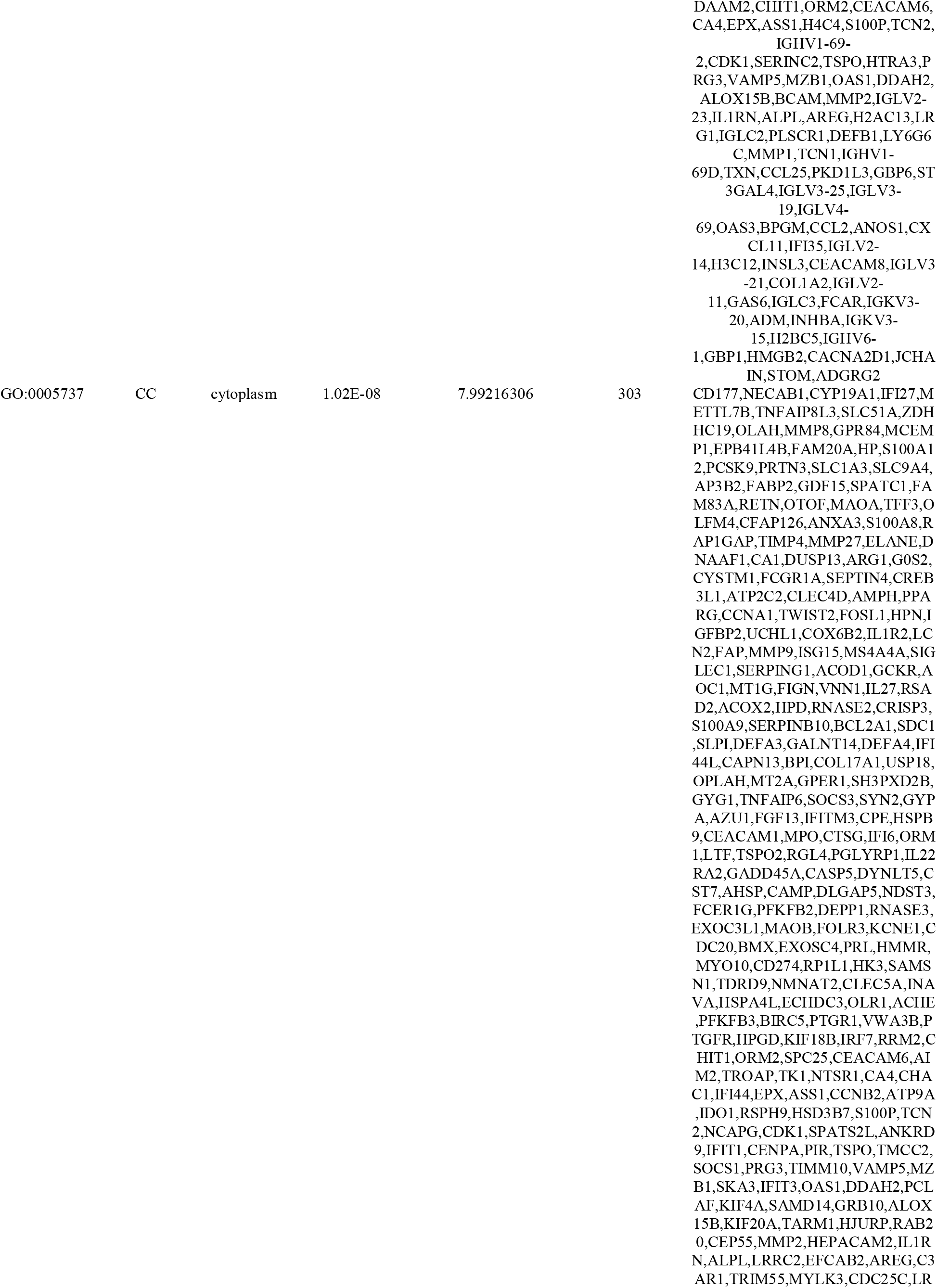
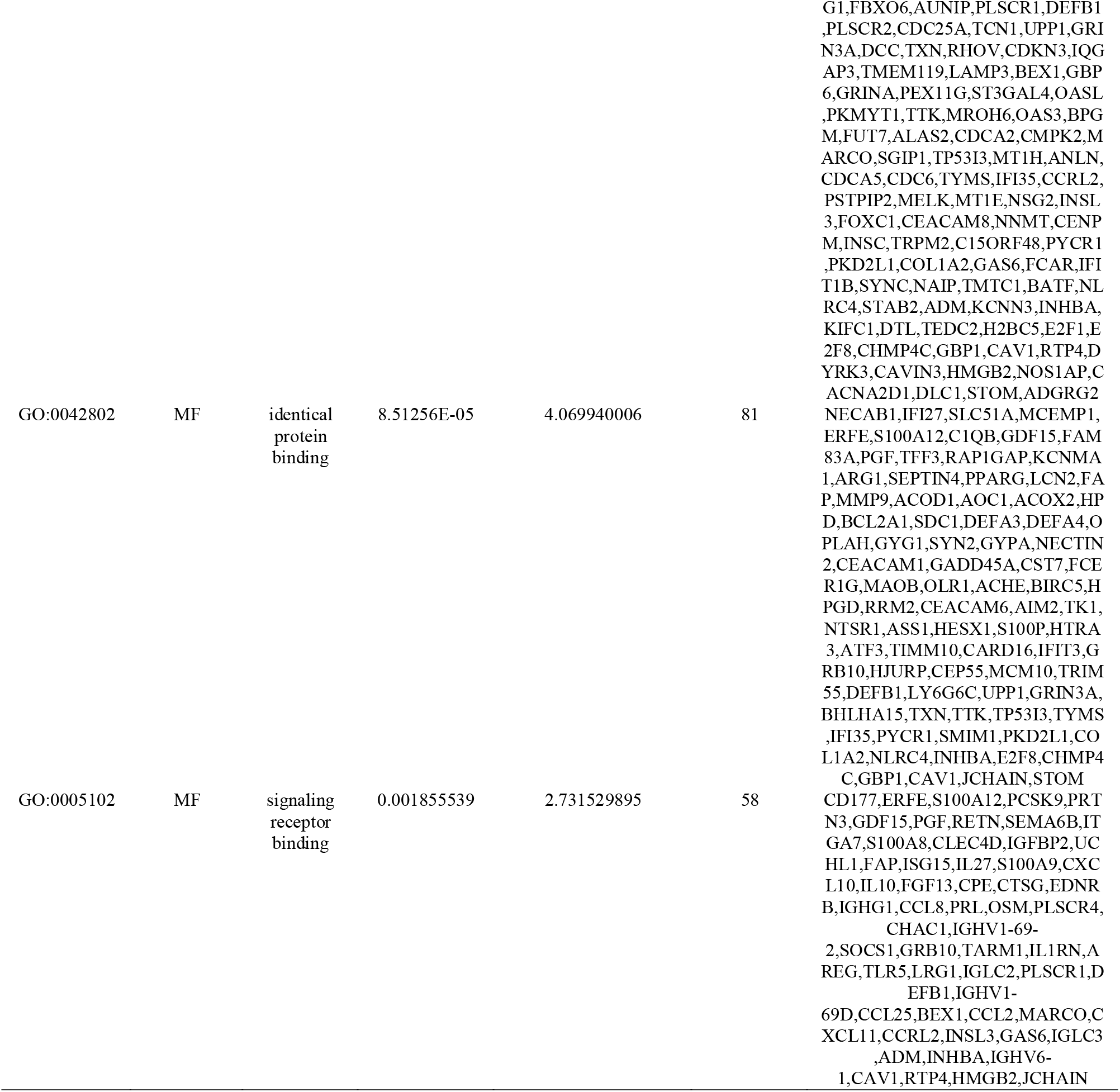
The enriched GO terms of the up and down regulated differentially expressed genes.

**Table 3.**
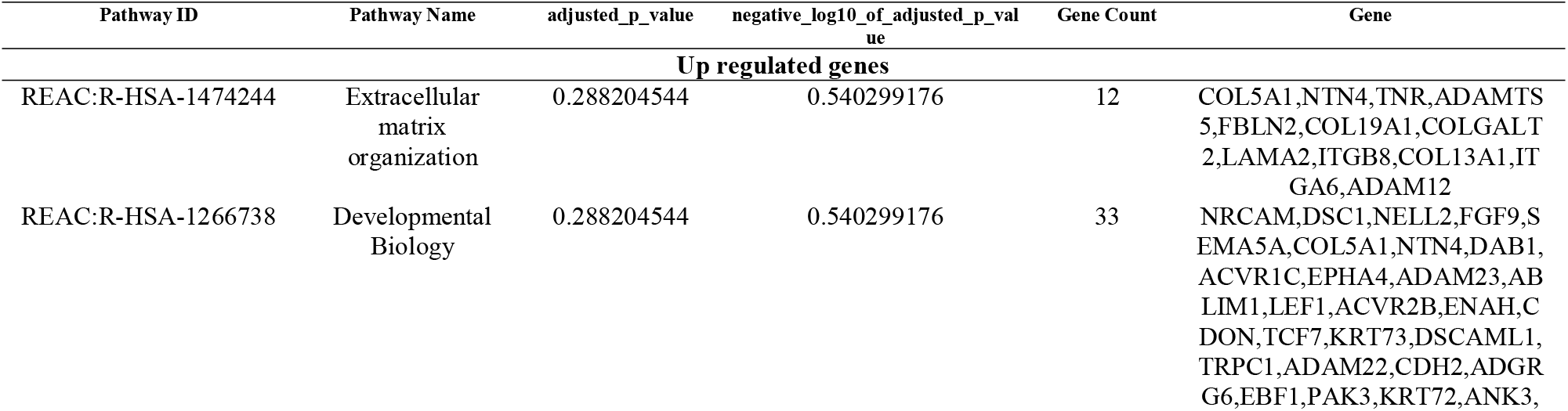
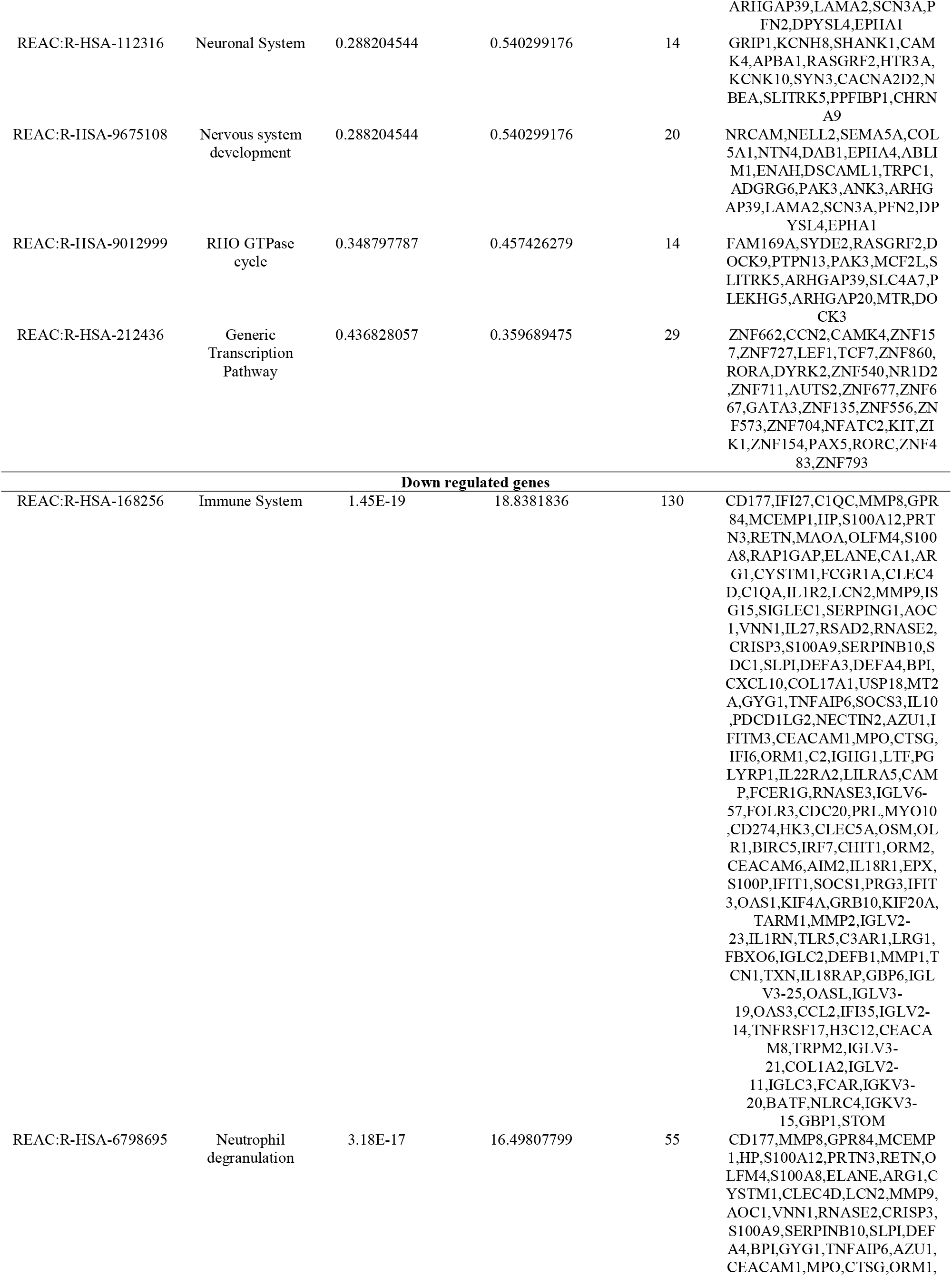
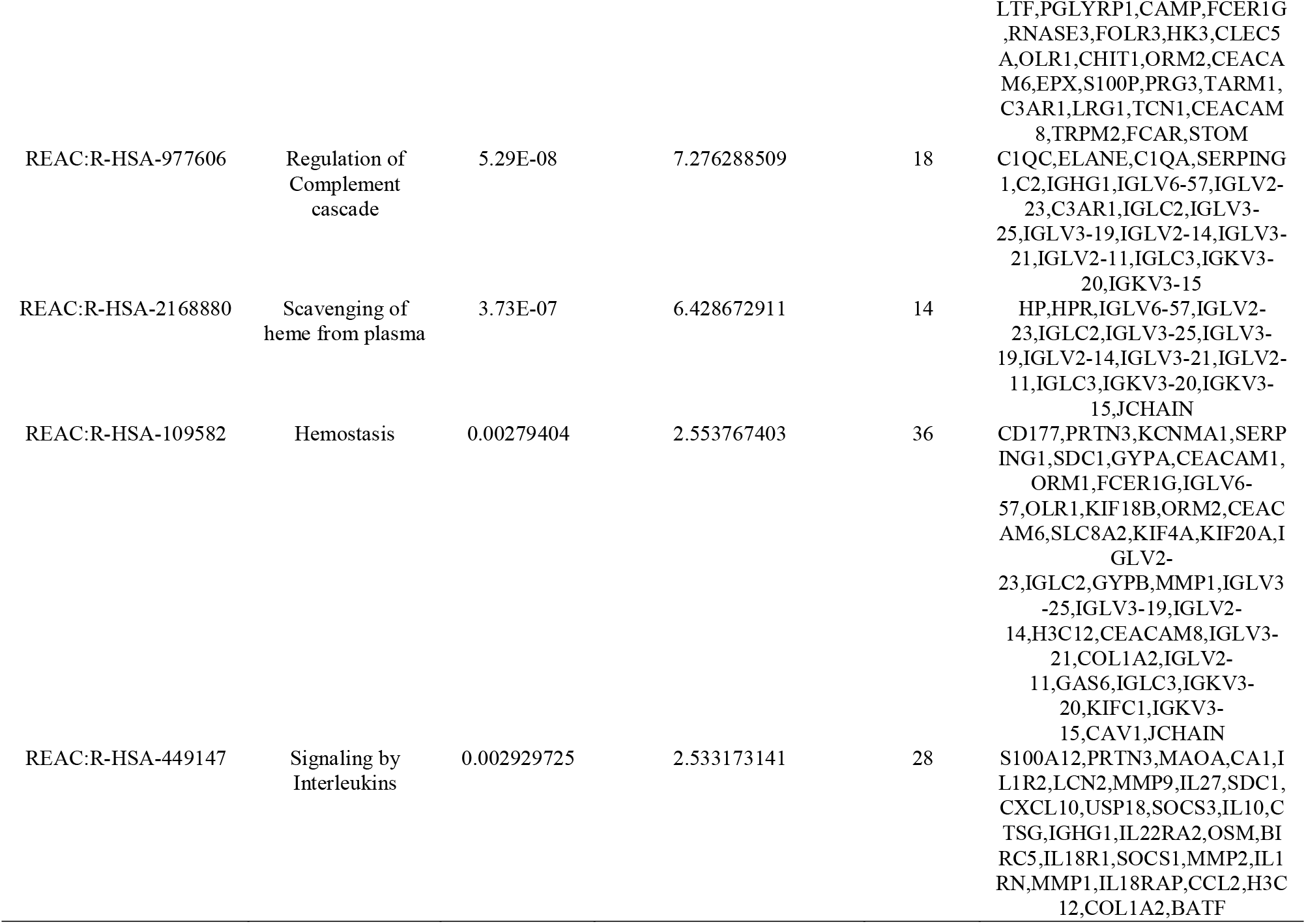
The enriched pathway terms of the up and down regulated differentially expressed genes.

### Construction of the PPI network and module analysis

The STRING database was applied to determine the PPI networks with DEGs, which were constructed via Cytoscape software, respectively (Fig. 3). Generally, the PPI network covered 4336 nodes and 10427 edges. Network Analyzer plugin of Cytoscape was used to rank the top nodes in the above PPI networks according to 4 topological analysis methods, including degree, betweenness, stress and closeness parameters. The hub genes according to the 4 methods were PRKN, KIT, FGFR2, GATA3, ERBB3, CDK1, PPARG, H2BC5, H4C4 and CDC20 (Table 4). Furthermore, PEWCC application results were indicative of two significant modules. Module 1 and module 2 were significant modules in the PPI network. A total of 20 nodes and 42 edges were included in module 1 (Fig. 4A), mainly involved in regulation of cellular process, generic transcription pathway and regulation of biological process and a total of 22 nodes and 53 edges were included in module 2 (Fig. 4B), which was associated with response to stimulus, cytoplasm and signaling.

**Fig. 3.**
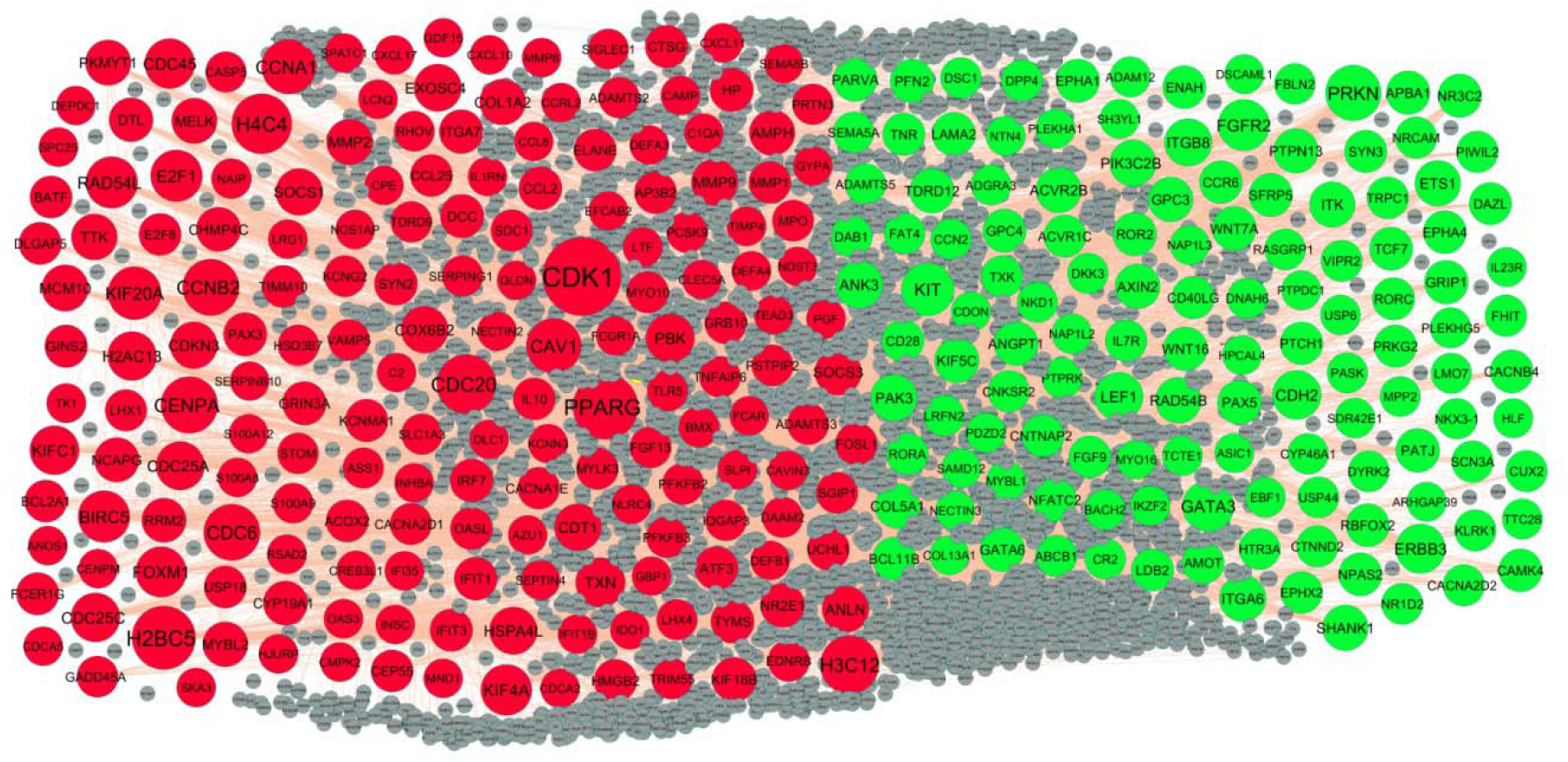
PPI network of DEGs. Up regulated genes are marked in parrot green; down regulated genes are marked in red

**Fig. 4.**
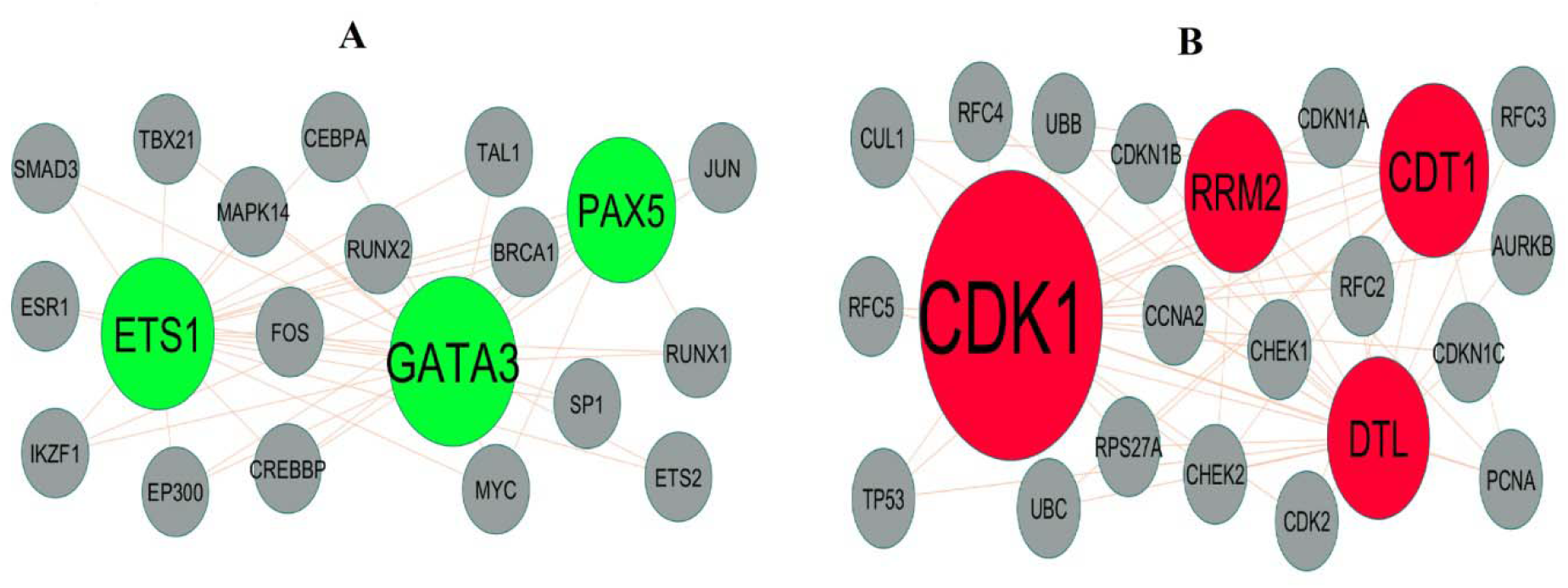
Modules selected from the PPI network. (A) The most significant module was obtained from PPI network with 20 nodes and 42 edges for up regulated genes (B) The most significant module was obtained from PPI network with 22 nodes and 53 edges for down regulated genes. Up regulated genes are marked in parrot green; down regulated genes are marked in red

**Table 4.**
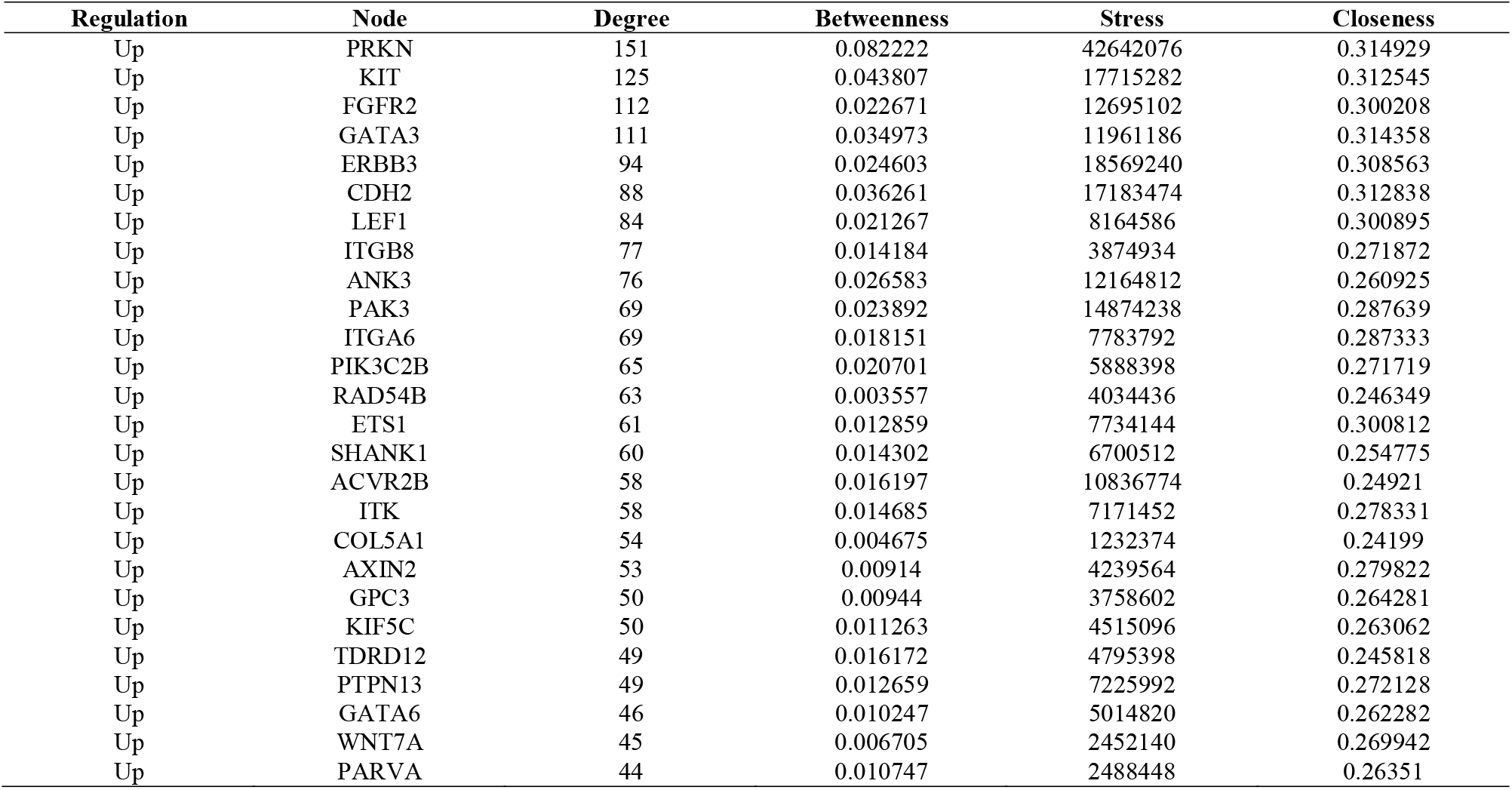
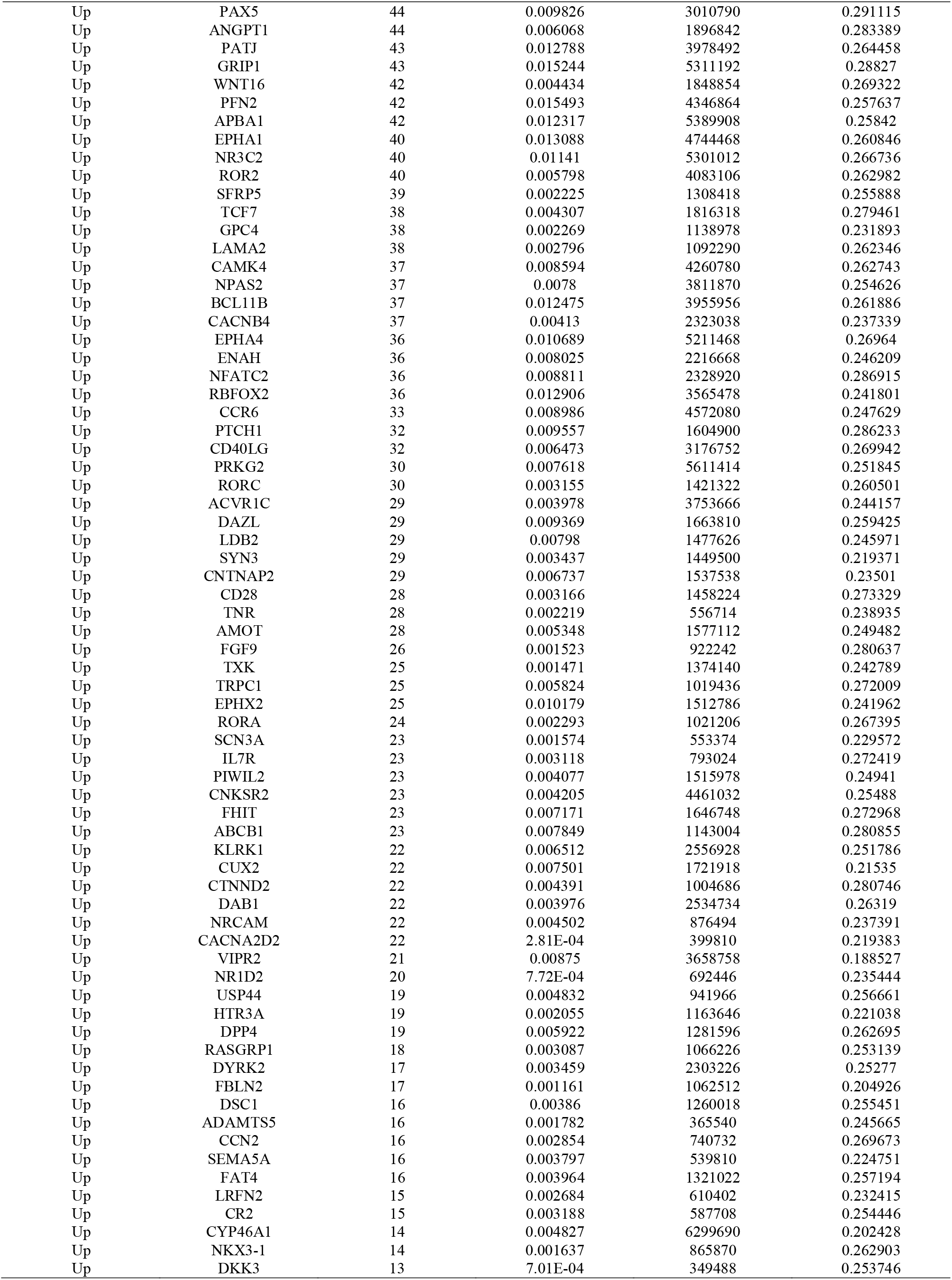
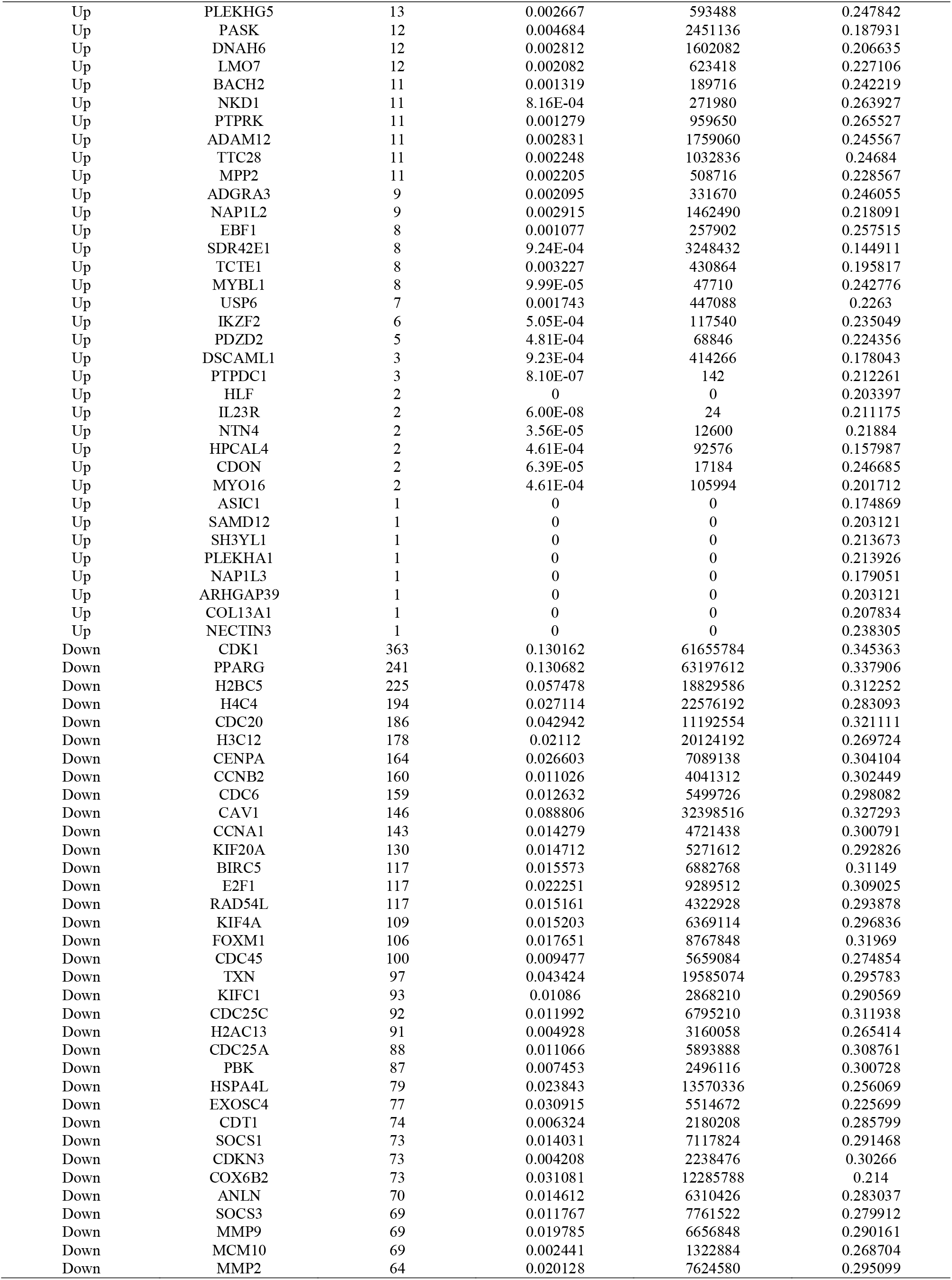
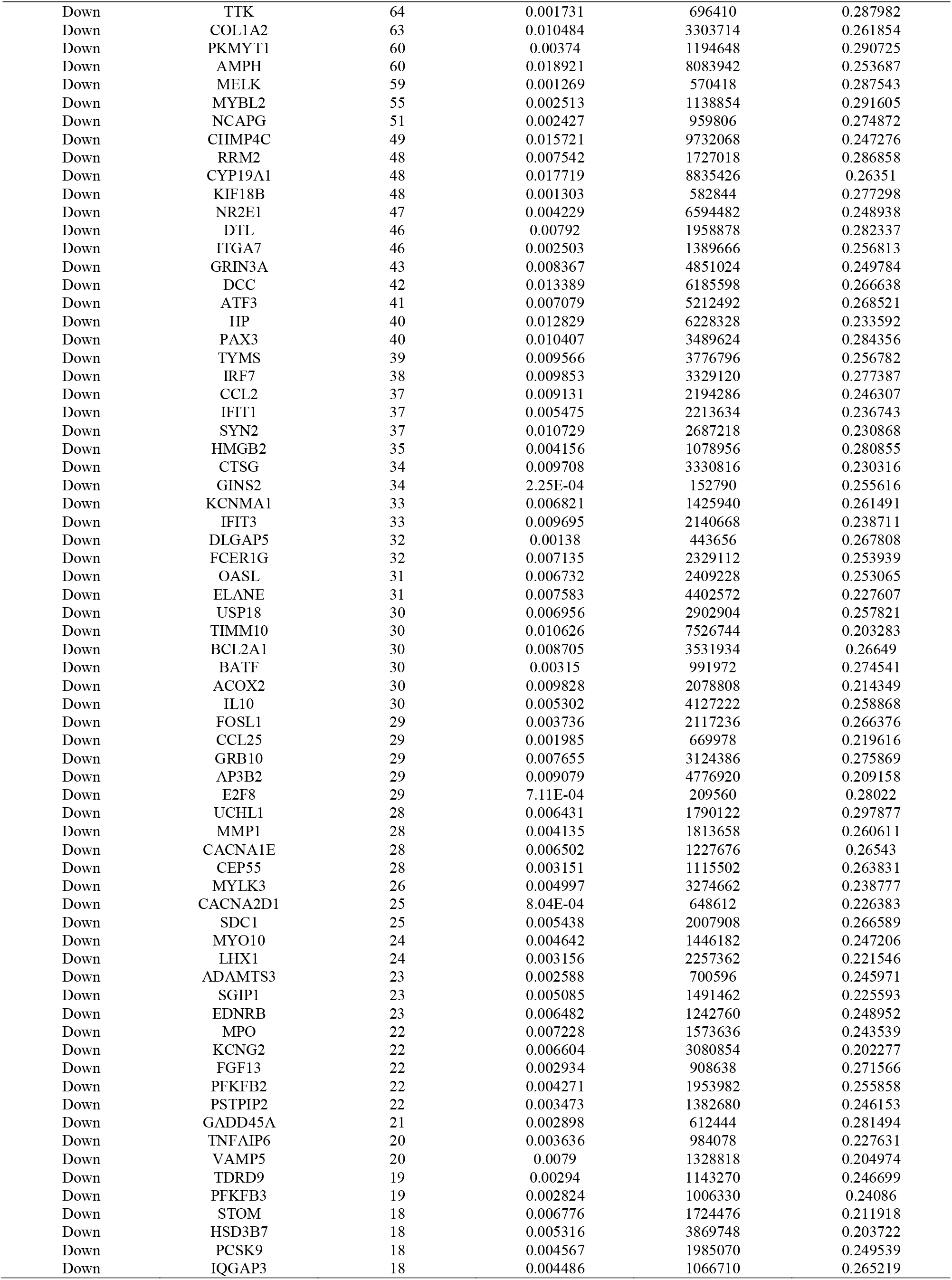
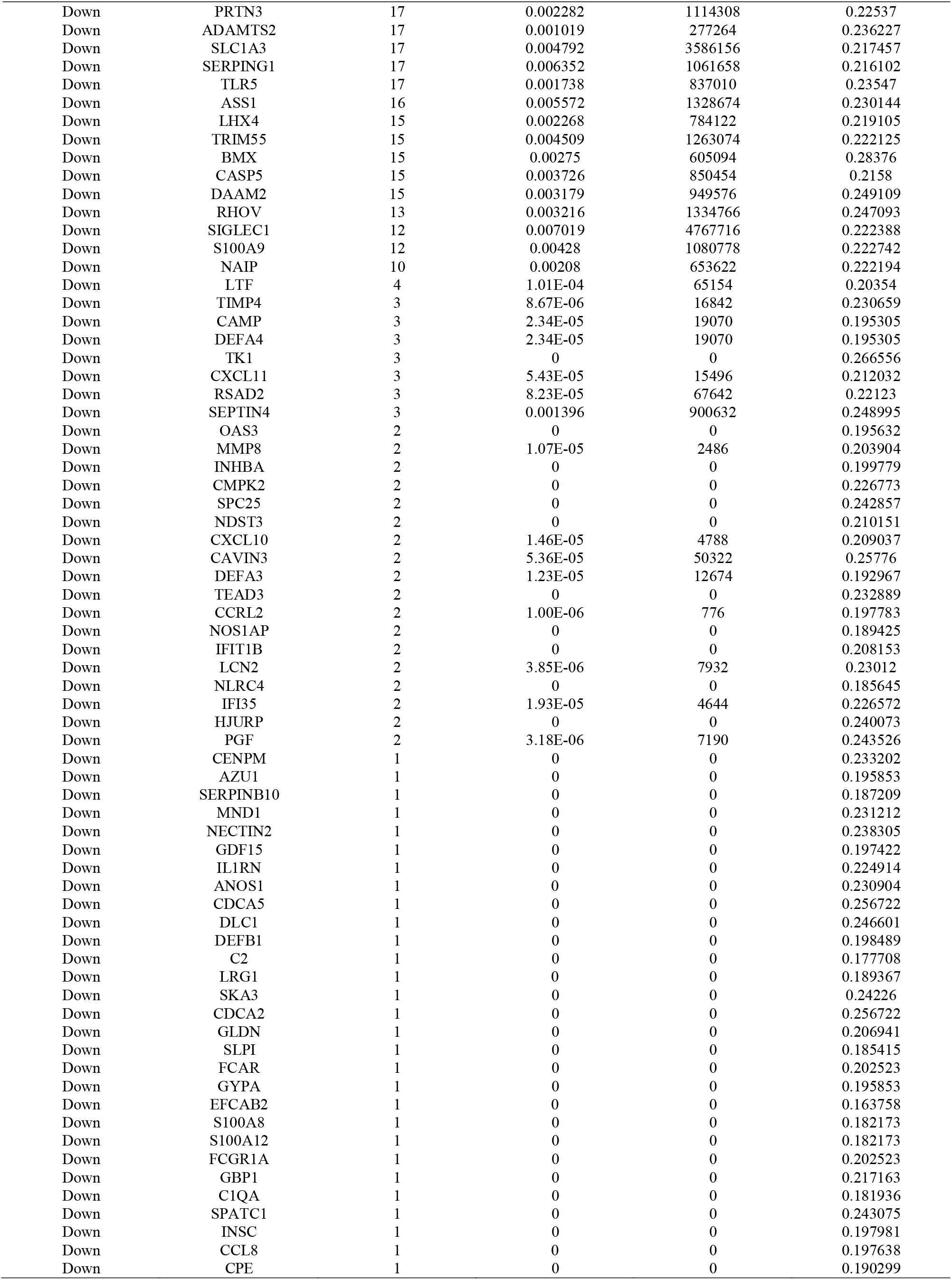
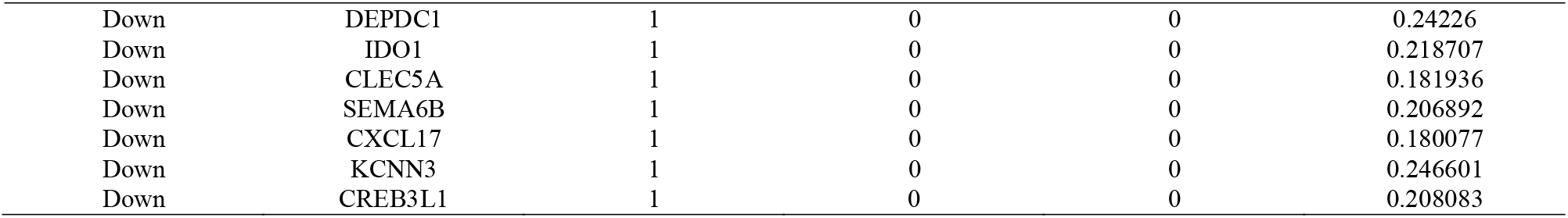
Topology table for up and down regulated genes.

### Construction of the miRNA-hub gene regulatory network

The miRNA-hub gene regulatory network of the hub genes was constructed, which contained 2411 nodes (miRNA: 2107; hub gene: 304) and 13734 edges (Fig.5). ITGB8 that was modulated by 146 miRNAs (ex: hsa-mir-548ad-5p); ETS1 that was modulated by 122 miRNAs (ex: hsa-mir-182-5p); PIK3C2B that was modulated by 94 miRNAs (ex: hsa-mir-3679-3p); ITGA6 that was modulated by 70 miRNAs (ex: hsa-mir-3145-3p); PAK3 that was modulated by 59 miRNAs (ex: hsa-mir-2116-3p); BIRC5 that was modulated by 135 miRNAs (ex: hsa-mir-2113); E2F1 that was modulated by 121 miRNAs (ex: hsa-mir-223-3p); CAV1 that was modulated by 115 miRNAs (ex: hsa-mir-652-3p); CDK1 that was modulated by 109 miRNAs (ex: hsa-mir-548ay-3p); CDC6 that was modulated by 71 miRNAs (ex: hsa-mir-106b-5p) (Table 5).

**Fig. 5.**
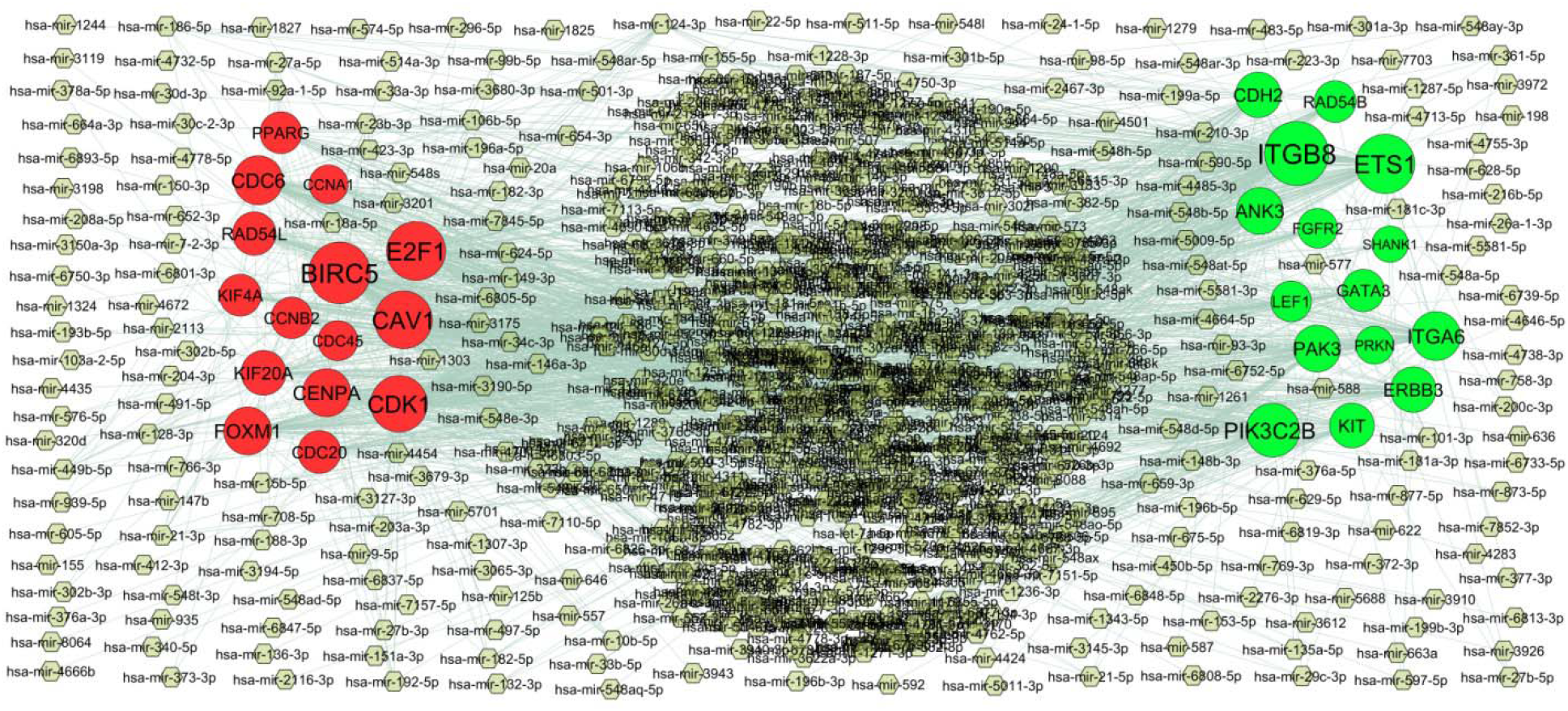
Hub gene - miRNA regulatory network. The pista green color diamond nodes represent the key miRNAs; up regulated genes are marked in green; down regulated genes are marked in red.

**Table 5.** MiRNA - hub gene and TF – hub gene topology table.

| Regulation | Hub Genes | Degree | MicroRNA | Regulation | Hub Genes | Degree | TF |
| --- | --- | --- | --- | --- | --- | --- | --- |
| Up | ITGB8 | 146 | hsa-mir-548ad-5p | Up | LEF1 | 49 | YAP1 |
| Up | ETS1 | 122 | hsa-mir-182-5p | Up | ANK3 | 49 | BCL3 |
| Up | PIK3C2B | 94 | hsa-mir-3679-3p | Up | FGFR2 | 47 | PHC1 |
| Up | ITGA6 | 70 | hsa-mir-3145-3p | Up | ETS1 | 46 | ATF3 |
| Up | PAK3 | 59 | hsa-mir-2116-3p | Up | GATA3 | 45 | RUNX2 |
| Up | ANK3 | 59 | hsa-mir-15b-5p | Up | CDH2 | 38 | MEF2A |
| Up | ERBB3 | 53 | hsa-mir-200c-3p | Up | KIT | 34 | GATA1 |
| Up | CDH2 | 52 | hsa-mir-1244 | Up | PIK3C2B | 33 | GFI1B |
| Up | KIT | 48 | hsa-mir-23b-3p | Up | ITGA6 | 33 | TFAP2C |
| Up | GATA3 | 36 | hsa-mir-124-3p | Up | RAD54B | 29 | SREBF2 |
| Up | RAD54B | 33 | hsa-mir-181a-3p | Up | ITGB8 | 29 | SOX2 |
| Up | LEF1 | 28 | hsa-mir-147b | Up | ERBB3 | 25 | NFIB |
| Up | FGFR2 | 23 | hsa-mir-3194-5p | Up | ACVR2B | 14 | REST |
| Up | PRKN | 14 | hsa-mir-193b-5p | Up | PAK3 | 12 | DNAJC2 |
| Up | SHANK1 | 6 | hsa-mir-491-5p | Up | SHANK1 | 8 | SMAD4 |
| Down | BIRC5 | 135 | hsa-mir-2113 | Down | CDC6 | 40 | TBX5 |
| Down | E2F1 | 121 | hsa-mir-223-3p | Down | PPARG | 39 | CUX1 |
| Down | CAV1 | 115 | hsa-mir-652-3p | Down | CDC45 | 36 | ESRRB |
| Down | CDK1 | 109 | hsa-mir-548ay-3p | Down | CAV1 | 36 | TET1 |
| Down | CDC6 | 71 | hsa-mir-106b-5p | Down | E2F1 | 35 | KIF20A |
| Down | CENPA | 66 | hsa-mir-9-5p | Down | CENPA | 32 | ASH2L |
| Down | FOXM1 | 65 | hsa-mir-98-5p | Down | BIRC5 | 27 | ESR1 |
| Down | KIF20A | 49 | hsa-mir-10b-5p | Down | KIF20A | 27 | CHD1 |
| Down | RAD54L | 42 | hsa-mir-21-3p | Down | RAD54L | 21 | POU5F1 |
| Down | CDC20 | 39 | hsa-mir-577 | Down | CDC20 | 21 | XRN2 |
| Down | PPARG | 34 | hsa-mir-548d-5p | Down | CCNB2 | 21 | OLIG2 |
| Down | KIF4A | 34 | hsa-mir-449b-5p | Down | CDK1 | 19 | NUCKS1 |
| Down | CCNB2 | 32 | hsa-mir-514a-3p | Down | FOXM1 | 19 | MYCN |
| Down | CDC45 | 25 | hsa-mir-203a-3p | Down | CCNA1 | 14 | STAT3 |
| Down | CCNA1 | 19 | hsa-mir-101-3p | Down | KIF4A | 10 | PBX1 |

### Construction of the TF-hub gene regulatory network

The TF-hub gene regulatory network of the hub genes was constructed, which contained 495 nodes (TF: 195; Hub gene: 300) and 6972 edges (Fig.6). LEF1 that was modulated by 49 TFs (ex: YAP1); ANK3 that was modulated by 49 TFs (ex: BCL3); FGFR2 that was modulated by 47 TFs (ex: PHC1); ETS1 that was modulated by 46 TFs (ex: ATF3); GATA3 that was modulated by 45 TFs (ex: RUNX2); CDC6 that was modulated by 40 TFs (ex: TBX5); PPARG that was modulated by 39 TFs (ex: CUX1); CDC45 that was modulated by 36 TFs (ex: ESRRB); CAV1 that was modulated by 36 TFs (ex: TET1); E2F1 that was modulated by 35 TFs (ex: KIF20A) (Table 5).

**Fig. 6.**
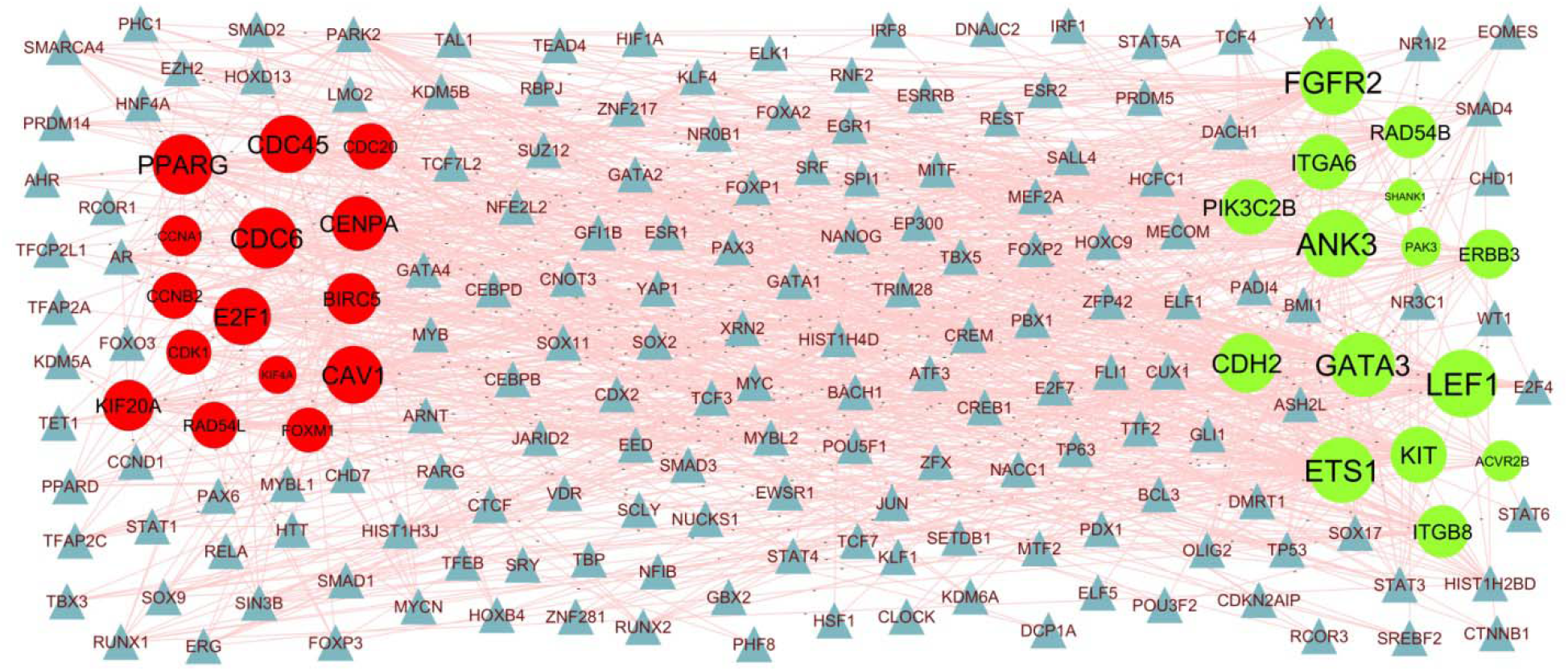
Hub gene - TF regulatory network. The blue color triangle nodes represent the key TFs; up regulated genes are marked in green; down regulated genes are marked in red.

### Receiver operating characteristic curve (ROC) analysis

To validate the diagnostic ability of the hub genes obtained from the above mentioned analysis, ROC curves were constructed and the AUC value was utilized to determine the diagnostic effectiveness in distinguishing sepsis from normal control sample. Fig.7 shows the AUC for PRKN was 0.910, AUC for KIT was 0.914, AUC for FGFR2 was 0.886, AUC for GATA3 was 0.924, AUC for ERBB3 was 0.938, AUC for CDK1 was 0.933, AUC for PPARG was 0.890, AUC for H2BC5 was 0.905, AUC for H4C4 was 0.900 and AUC for CDC20 was 0.919, indicating that the hub genes had high diagnostic ability.

**Fig. 7.**
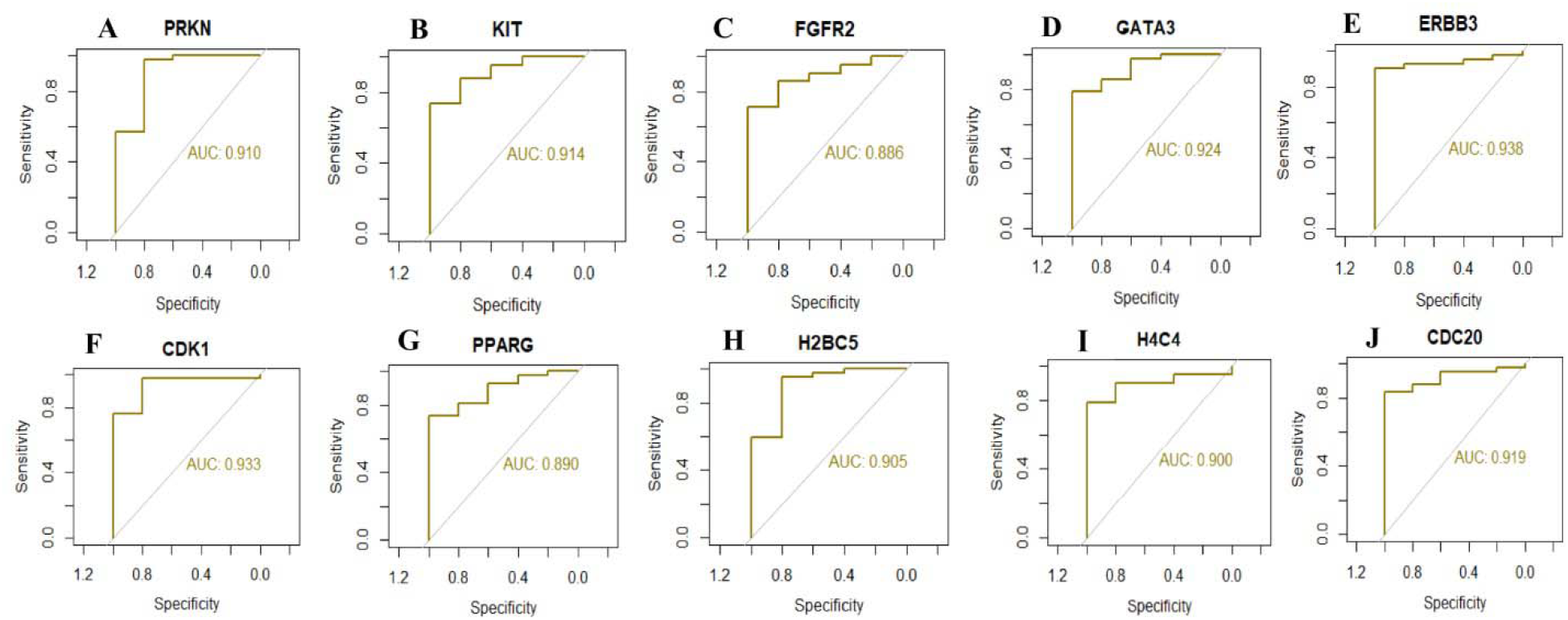
ROC curve analyses of hub genes. A) PRKN B) KIT C) FGFR2 D) GATA3 E) ERBB3 F) CDK1 G) PPARG H) H2BC5 I) H4C4 J) CDC20

## Discussion

Sepsis is a highly prevalent disease around world, which remains a leading cause of death [49]. Therefore, timely and appropriate diagnosis of sepsis is essential to improve the prognosis. Hence, we aimed to identify novel diagnostic biomarkers to screen for patients with sepsis and reveal the role of systemic inflammatory response in sepsis.

In this investigation, we collected NGS data sets from the GEO database, and a total of 958 DEGs, including 479 up regulated genes and 479 down regulated genes, were found. Previous studies have demonstrated that NRCAM (neuronal cell adhesion molecule) [50], NOG (noggin) [51], LRRN3 [52], SPON1 [53], NECAB1 [54] and CYP19A1 [55] are linked with the development mechanisms of brain dysfunction. NOG (noggin) [56] and CD177 [57] participates in pathogenic processes of liver dysfunction. SIAH3 [58] and CD177 [59] are involved in the development and progression of kidney dysfunction. CD177 [60] and METTL7B [61] were found to promote the sepsis. CD177 [62] plays an important role in regulating the septic shock. IFI27 [63] plays an important role in the microbial infections. METTL7B [61] has been known to be involved in inflammation. C1QC [64] has a significant prognostic potential in heart dysfunction. These genes might be served as biomarkers for sepsis diagnosis and prognosis.

GO and REACTOME enrichment analyses were performed to explore interactions among the DEGs. Signaling pathways include neuronal system [65], nervous system development [66], immune system [67], neutrophil degranulation [68], regulation of complement cascade [69], scavenging of heme from plasma [70] and hemostasis [71] are involved in sepsis. Studies had shown that SLC4A10 [72], FGF9 [73], AJAP1 [74], NR3C2 [75], SORCS3 [76], TNR (tenascin R) [77], ASIC1 [78], DAB1 [79], DKK3 [80], EPHA4 [81], CCN2 [82], CAMK4 [83], PTCH1 [84], BCL11B [85], AXIN2 [86], ECRG4 [87], NEXMIF (neurite extension and migration factor) [88], ERBB3 [89], IL23R [90], HPSE2 [91], TRPC1 [92], WNT7A [93], ADAM22 [94], RORA (RAR related orphan receptor A) [95], KCNK10 [96], ETS1 [97], RASGRP1 [98], TENM4 [99], CCR6 [100], CYP46A1 [101], ANGPT1 [102], ZNF391 [103], MEG3 [104], CACNA2D2 [105], NBEA (neurobeachin) [106], CNKSR2 [107], SLC4A4 [108], TCTN2 [109], ZNF667 [110], ABCB1 [111], RBFOX2 [112], GATA3 [113], DPP4 [114], SLITRK5 [115], TXK (TXK tyrosine kinase) [116], CCL28 [117], FNDC5 [118], ITGB8 [119], GATA6 [120], VIPR2 [121], CD28 [122], NFATC2 [123], TET1 [124], PIWIL2 [125], PRRT2 [126], SCN3A [127], PRKN (parkin RBR E3 ubiquitin protein ligase) [128], EPHA1 [129], ADAM12 [130], ADAMTS5 [131], NT5E [132], ADAM23 [133], SLC7A3 [134], PPFIBP1 [135], NECTIN3 [136], PDZD2 [137], EPHX2 [138], MTR (5-methyltetrahydrofolate-homocysteine methyltransferase) [139], NR2E1 [140], KLF14 [141], GPR84 [142], HP (haptoglobin) [143], S100A12 [144], PCSK9 [145], C1QB [146], GDF15 [147], FAM83A [148], RNASE1 [149], ANXA3 [150], S100A8 [151], ARG1 [152], SHOX2 [153], FOSL1 [154], C1QA [155], IGFBP2 [156], UCHL1 [157], IL1R2 [158], LCN2 [159], MMP9 [160], ISG15 [161], SIGLEC1 [162], GCKR (glucokinase regulator) [163], IL27 [164], RSAD2 [165], S100A9 [166], SDC1 [167], CXCL10 [168], USP18 [169], GPER1 [170], SOCS3 [171], IL10 [172], FGF13 [173], IFITM3 [174], CPE (carboxypeptidase E) [175], CEACAM1 [176], MPO (myeloperoxidase) [177], CTSG (cathepsin G) [178], VSIG4 [179], LTF (lactotransferrin) [180], TAAR1 [181], GADD45A [182], RBP4 [183], MAOB (monoamine oxidase B) [184], PRL (prolactin) [185], OSM (oncostatin M) [186], ACHE (acetylcholinesterase) [187], BIRC5 [188], IRF7 [189], CHIT1 [190], AIM2 [191], NTSR1 [192], HSD3B7 [193], CDH6 [194], TSPO (translocator protein) [195], SOCS1 [196], ATF3 [197], IFIT3 [198], OAS1 [199], GRB10 [200], RAB20 [201], MMP2 [202], TLR5 [203], C3AR1 [204], LRG1 [205], FBXO6 [206], MMP1 [207], GRIN3A [208], DCC (DCC netrin 1 receptor) [209], OASL (2’-5’-oligoadenylate synthetase like) [210], OAS3 [211], KREMEN1 [212], CCL2 [213], CMPK2 [214], GPR4 [215], IFI35 [216], FOXC1 [217], NNMT (nicotinamide N-methyltransferase) [218], TRPM2 [219], COL1A2 [220], GAS6 [221], NLRC4 [222], ADM (adrenomedullin) [223], E2F1 [224], CAV1 [225], RTP4 [226], CAVIN3 [227], HMGB2 [228], NOS1AP [229], CACNA2D1 [230], TFF3 [231], HPD (4-hydroxyphenylpyruvate dioxygenase) [232], NRN1 [233], CA4 [234], SERINC2 [235], MCEMP1 [236] and SYN2 [237] were associated with brain dysfunction. Studies have found that FGF9 [238], DKK3 [239], CAMK4 [240], SFRP5 [241], BCL11B [242], TCF7 [243], CDH2 [244], MYBL1 [245], RASGRP1 [246], MEG3 [247], DGKK (diacylglycerol kinase kappa) [248], DPP4 [249], CD28 [250], ITK (IL2 inducible T cell kinase) [251], GPR174 [252], HAVCR1 [253], KLF14 [254], MMP8 [255], HP (haptoglobin) [256], S100A12 [257], PCSK9 [258], GDF15 [259], PGF (placental growth factor) [260], RETN (resistin) [261], MAOA (monoamine oxidase A) [262], RNASE1 [263], OLFM4 [264], ANXA3 [265], S100A8 [266], ARG1 [267], FOSL1 [268], IL1R2 [269], LCN2 [270], MMP9 [271], VNN1 [272], IL27 [273], S100A9 [274], BCL2A1 [275], SDC1 [276], DEFA3 [277], BPI (bactericidal permeability increasing protein) [278], CXCL10 [279], USP18 [280], GPER1 [281], SOCS3 [282], IL10 [283], NECTIN2 [284], IFITM3 [285], CEACAM1 [286], MPO (myeloperoxidase) [287], C2 [288], VSIG4 [289], LTF (lactotransferrin) [290], LILRA5 [291], RNASE3 [292], PRL (prolactin) [293], SAMSN1 [294], OSM (oncostatin M) [295], ACHE (acetylcholinesterase) [296], CHIT1 [297], FOXM1 [298], TSPO (translocator protein) [299], SOCS1 [300], ATF3 [301], DDAH2 [302], MMP2 [303], IL1RN [304], TLR5 [305], LRG1 [205], DEFB1 [306], CCL25 [307], CCL2 [308], FOXC1 [217], TRPM2 [219], GAS6 [309], ADM (adrenomedullin) [310], E2F1 [311], CAV1 [312], TFF3 [313], SERINC2 [314], STOM (stomatin) [315] and MCEMP1 [316] are altered expression in sepsis. FGF9 [317], NR3C2 [318], DAB1 [319], CCN2 [320], CAMK4 [321], NKX3-1 [322], BCL11B [323], AMOT (angiomotin) [324], IL7R [325], ECRG4 [326], TRABD2A [327], TCF7 [328], IL23R [329], CR2 [330], TRPC1 [331], RORA (RAR related orphan receptor A) [332], ETS1 [333], CD40LG [334], CCR6 [335], ROR2 [336], MEG3 [337], ABCB1 [338], USP44 [339], ZBTB25 [340], GATA3 [341], DPP4 [342], CCL28 [343], CD28 [344], MMP28 [345], PIK3C2B [346], TSPAN6 [347], ITK (IL2 inducible T cell kinase) [348], THEMIS (thymocyte selection associated) [349], HAVCR1 [350], MXRA8 [351], CDHR3 [352], TRAV1-2 [353], MMP8 [354], GPR84 [355], HP (haptoglobin) [356], S100A12 [357], PCSK9 [358], PRTN3 [359], GDF15 [360], RETN (resistin) [361], WFDC1 [362], RNASE1 [363], OLFM4 [364], S100A8 [365], ELANE (elastase, neutrophil expressed) [366], ARG1 [367], TWIST2 [368], FOSL1 [369], C1QA [370], BATF2 [371], LCN2 [372], FAP (fibroblast activation protein alpha) [373], MMP9 [160], ISG15 [374], SIGLEC1 [375], SERPING1 [376], ACOD1 [377], IL27 [378], RNASE2 [379], S100A9 [380], SDC1 [381], DEFA3 [382], DEFA4 [383], IFI44L [384], BPI (bactericidal permeability increasing protein) [385], CXCL10 [386], USP18 [387], SOCS3 [388], IL10 [389], IFITM3 [390], CEACAM1 [391], CXCL17 [392], HPR (haptoglobin-related protein) [393], MPO (myeloperoxidase) [394], LY6E [395], CTSG (cathepsin G) [396], IFI6 [397], ORM1 [398], VSIG4 [399], LTF (lactotransferrin) [400], CCL8 [401], PGLYRP1 [402], RBP4 [403], CAMP (cathelicidin antimicrobial peptide) [404], RNASE3 [405], CLEC5A [406], OSM (oncostatin M) [407], BIRC5 [408], IRF7 [409], CHIT1 [410], IFI44 [411], CEACAM6 [412], AIM2 [413], IL18R1 [414], CHAC1 [415], EPX (eosinophil peroxidase) [416], IDO1 [397], S100P [417], CDK1 [418], IFIT1 [419], FOXM1 [420], TSPO (translocator protein) [421], SOCS1 [388], ATF3 [422], IFIT3 [198], OAS1 [199], TARM1 [423], RAB20 [424], MMP2 [425], IL1RN [426], AREG (amphiregulin) [427], TLR5 [428], PLSCR1 [429], DEFB1 [430], MMP1 [431], IL18RAP [432], LAMP3 [433], OASL (2’-5’-oligoadenylate synthetase like) [434], TTK (TTK protein kinase) [435], OAS3 [211], BPGM (bisphosphoglyceratemutase) [436], KREMEN1 [437], CCL2 [438], CMPK2 [439], MARCO (macrophage receptor with collagenous structure) [440], GPR4 [441], CDC6 [442], CXCL11 [443], IFI35 [216], CCRL2 [444], TRPM2 [445], GAS6 [446], NAIP (NLR family apoptosis inhibitory protein) [447], BATF (basic leucine zipper ATF-like transcription factor) [448], NLRC4 [449], ADM (adrenomedullin) [450], CHMP4C [451], GBP1 [452], CAV1 [453], RTP4 [226], CAVIN3 [454], HMGB2 [455], TFF3 [456], MCEMP1 [457], TK1 [458] and KIF20A [459] plays an indispensable role in microbial infections. Studies have shown that FGF9 [460], GRIP1 [461], TNR (tenascin R) [462], ASIC1 [463], DKK3 [464], EPHA4 [465], CCN2 [466], CAMK4 [467], SFRP5 [468], PTCH1 [469], NKX3-1 [470], ZNF365 [471], AMOT (angiomotin) [472], IL7R [473], BACH2 [474], LEF1 [475], ECRG4 [476], ERBB3 [477], TCF7 [478], IL23R [479], TRPC1 [480], WNT7A [481], RORA (RAR related orphan receptor A) [332], WNT16 [482], ABCD2 [483], ETS1 [484], RASGRP1 [485], FGFR2 [486], CCR6 [487], EBF1 [488], ROR2 [489], CCN3 [490], MEG3 [491], FXYD2 [492], ZNF667 [493], ABCB1 [494], USP44 [495], GATA3 [496], DPP4 [497], TXK (TXK tyrosine kinase) [498], FNDC5 [499], ITGB8 [119], THEM4 [500], GATA6 [501], CD28 [502], NFATC2 [503], TET1 [504], MMP28 [505], ITK (IL2 inducible T cell kinase [506], GPR174 [507], THEMIS (thymocyte selection associated) [508], TIFAB (TIFA inhibitor) [509], ITGA6 [510], ADAM12 [511], SCART1 [512], NTN4 [513], ADAMTS5 [514], NT5E [132], SCARA5 [515], PPFIBP1 [516], EPHX2 [517], MPPED2 [518], MORC4 [519], MICU3 [520], NR2E1 [521], KLF14 [522], MMP8 [523], GPR84 [524], ERFE (erythroferrone) [525], HP (haptoglobin) [526], S100A12 [527], PCSK9 [528], GDF15 [529], RETN (resistin) [361], MAOA (monoamine oxidase A) [530], WFDC1 [531], OLFM4 [532], S100A8 [266], TIMP4 [533], ELANE (elastase, neutrophil expressed) [366], ARG1 [152], TWIST2 [368], FOSL1 [154], IGFBP2 [156], UCHL1 [534], IL1R2 [535], LCN2 [536], FAP (fibroblast activation protein alpha) [537], MMP9 [538], ISG15 [539], ACOD1 [377], GCKR (glucokinase regulator) [540], MT1G [541], VNN1 [542], IL27 [543], RNASE2 [544], CRISP3 [545], S100A9 [546], BCL2A1 [547], SDC1 [548], DEFA3 [549], BPI (bactericidal permeability increasing protein) [550], CXCL10 [551], USP18 [387], MT2A [552], GPER1 [553], TNFAIP6 [554], SOCS3 [555], IL10 [556], FGF13 [557], IFITM3 [558], CPE (carboxypeptidase E) [559], CEACAM1 [560], CXCL17 [561], MPO (myeloperoxidase) [562], LY6E [563], CTSG (cathepsin G) [564], EDNRB (endothelin receptor type B) [565], VSIG4 [566], LTF (lactotransferrin) [567], CCL8 [568], PGLYRP1 [569], TAAR1 [570], GADD45A [571], RBP4 [572], CAMP (cathelicidin antimicrobial peptide) [573], RNASE3 [574], MAOB (monoamine oxidase B) [575], CDC20 [576], BMX (BMX non-receptor tyrosine kinase) [577], PRL (prolactin) [185], MYO10 [578], CLEC5A [579], OSM (oncostatin M) [580], OLR1 [581], ACHE (acetylcholinesterase) [296], BIRC5 [582], IRF7 [583], CHIT1 [410], CEACAM6 [584], AIM2 [413], NTSR1 [585], IL18R1 [586], CHAC1 [587], EPX (eosinophil peroxidase) [588], ASS1 [589], IDO1 [590], S100P [417], CDH6 [591], IFIT1 [592], FOXM1 [593], TSPO (translocator protein) [594], HTRA3 [595], SOCS1 [596], ATF3 [597], MZB1 [598], IFIT3 [599], GRB10 [600], ALOX15B [601], TARM1 [602], RAB20 [603], MMP2 [604], IL1RN [605], AREG (amphiregulin) [606], TLR5 [428], LRG1 [607], DEFB1 [608], CDC25A [609], CCL25 [307], IL18RAP [432], KREMEN1 [610], CCL2 [438], MARCO (macrophage receptor with collagenous structure) [611], GPR4 [215], CXCL11 [612], FOXC1 [217], NNMT (nicotinamide N-methyltransferase) [613], TRPM2 [614], C15ORF48 [615], GAS6 [616], NAIP (NLR family apoptosis inhibitory protein) [617], NLRC4 [617], ADM (adrenomedullin) [618], E2F1 [619], GBP1 [620], CAV1 [621], HMGB2 [228], NOS1AP [622], TFF3 [623], SERPINB10 [624], STOM (stomatin) [315] and MCEMP1 [625] are active in inflammation. Altered expression of FGF9 [626], SEMA5A [627], NR3C2 [628], NPAS2 [629], DKK3 [630], EPHA4 [631], CCN2 [632], SFRP5 [633], BACH2 [634], ECRG4 [635], MTUS1 [636], ERBB3 [637], TCF7 [638], TRPC1 [639], RORA (RAR related orphan receptor A) [640], MYBL1 [245], ETS1 [641], FGFR2 [642], CCR6 [643], RBM20 [644], EBF1 [645], CCN3 [646], MEG3 [647], CAMK2N1 [648], CACNA2D2 [649], CLIC5 [650], EFHD1 [651], ZNF667 [652], ABCB1 [653], RBFOX2 [654]. DCBLD2 [655], DPP4 [656], MCF2L [657], FNDC5 [658], GATA6 [659], NFATC2 [660], MMP28 [661], GPR174 [662], DACT1 [663], ADAM12 [664], ADAMTS5 [665], ADAM23 [666], PARVA (parvin alpha) [667], CDHR3 [668], SLC4A7 [669], MMP21 [670], EPHX2 [671], MPPED2 [672], CARNS1 [673], KLF14 [674], MMP8 [675], HP (haptoglobin) [676], S100A12 [677], PCSK9 [678], GDF15 [679], PGF (placental growth factor) [680], RETN (resistin) [681], ITGA7 [682], RNASE1 [263], LHX4 [683], ANXA3 [684], S100A8 [685], RAP1GAP [686], TIMP4 [533], ELANE (elastase, neutrophil expressed) [687], ARG1 [688], SHOX2 [689], CLEC4D [690], IGFBP2 [691], UCHL1 [692], IL1R2 [693], LCN2 [694], FAP (fibroblast activation protein alpha) [695], MMP9 [696], ISG15 [697], SERPING1 [698], GCKR (glucokinase regulator) [699], VNN1 [700], ACOX2 [701], RNASE2 [544], S100A9 [546], SDC1 [702], DEFA3 [549], BPI (bactericidal permeability increasing protein) [703], MT2A [704], GPER1 [705], PBK (PDZ binding kinase) [706], SOCS3 [707], IL10 [708], NECTIN2 [709], FGF13 [710], CPE (carboxypeptidase E) [711], CEACAM1 [712], MPO (myeloperoxidase) [562], CTSG (cathepsin G) [178], ORM1 [713], VSIG4 [714], LTF (lactotransferrin) [715], CCL8 [716], PGLYRP1 [717], RBP4 [718], CAMP (cathelicidin antimicrobial peptide) [719], FCER1G [720], PFKFB2 [721], MAOB (monoamine oxidase B) [722], KCNE1 [723], CDC20 [724], PRL (prolactin) [725], CLEC5A [726], OSM (oncostatin M) [727], ECHDC3 [728], OLR1 [729], ACHE (acetylcholinesterase) [730], BIRC5 [731], IRF7 [732], CHIT1 [733], AIM2 [734], PLSCR4 [735], IL18R1 [414], EPX (eosinophil peroxidase) [736], IDO1 [737], S100P [738], CDK1 [739], FOXM1 [740], TSPO (translocator protein) [741], HTRA3 [742], SOCS1 [743], ATF3 [744], MZB1 [745], IFIT3 [599], DDAH2 [746], GRB10 [747], ALOX15B [748], MMP2 [749], IL1RN [750], ALPL (alkaline phosphatase, biomineralization associated) [751], AREG (amphiregulin) [752], TLR5 [753], TRIM55 [754], MYLK3 [755], LRG1 [756], MMP1 [757], MYBL2 [758], DCC (DCC netrin 1 receptor) [759], BEX1 [760], ALAS2 [761], CCL2 [762], CMPK2 [763], GPR4 [764], CCRL2 [765], FOXC1 [766], TRPM2 [767], PKD2L1 [768], COL1A2 [220], GAS6 [769], STAB2 [770], ADM (adrenomedullin) [771], CAV1 [621], HMGB2 [772], NOS1AP [773], CRISP2 [774], RRM2 [775] and KIF20A [776] are associated with prognosis of heart dysfunction. FGF9 [777], EHF [778], CCN2 [779], CAMK4 [780], PTCH1 [781], TPH1 [782], ZNF365 [783], IL7R [784], LEF1 [785], ECRG4 [786], TCF7 [787], WNT7A [788], CDH2 [244], ETS1 [789], FGFR2 [790], CCR6 [791], PTPN13 [792], EBF1 [793], ROR2 [794], ANGPT1 [795], CCN3 [796], MEG3 [797], DGKK (diacylglycerol kinase kappa) [248], DPP4 [656], GATA6 [798], CD28 [799], ENHO (energy homeostasis associated) [800], NFATC2 [801], TET1 [802], MMP28 [803], PIWIL2 [804], ITK (IL2 inducible T cell kinase) [805], ADAM12 [806], KLF14 [807]. MMP8 [808], GPR84 [809], HP (haptoglobin) [810], S100A12 [811], GDF15 [812], RETN (resistin) [813], OLFM4 [814], S100A8 [815], IGFBP2 [816], LCN2 [817], FAP (fibroblast activation protein alpha) [818], MMP9 [271], SIGLEC1 [819], VNN1 [820], IL27 [821], S100A9 [822], SDC1 [823], BPI (bactericidal permeability increasing protein) [824], CXCL10 [825], CXCL10 [826], SOCS3 [827], IL10 [828], CPE (carboxypeptidase E) [559], CEACAM1 [829], MPO (myeloperoxidase) [830], CCL8 [831], PGLYRP1 [402], GADD45A [571], RBP4 [832], SAMSN1 [294], CLEC5A [406], OSM (oncostatin M) [833], PTGFR (prostaglandin F receptor) [834], IRF7 [835], CHIT1 [836], AIM2 [837], IL18R1 [838], IFIT1 [839], FOXM1 [420], TSPO (translocator protein) [840], SOCS1 [841], ATF3 [301], MMP2 [842], IL1RN [843], AREG (amphiregulin) [844], CCL25 [307], CCL2 [845], CMPK2 [846], MARCO (macrophage receptor with collagenous structure) [611], TRPM2 [847], GAS6 [848], ADM (adrenomedullin) [849], CAV1 [850], HMGB2 [851], SERINC2 [314] and STOM (stomatin) [315] can contribute to lung dysfunction. FGF9 [852], SEMA5A [853], NR3C2 [854], HLF (HLF transcription factor, PAR bZIP family member) [855], GDF10 [856], CCN2 [857], CAMK4 [858], SFRP5 [859], PTCH1 [860], IL7R [861], IL23R [862], LMO7 [863], RORA (RAR related orphan receptor A) [864], ETS1 [865], CD40LG [866], FGFR2 [867], CCR6 [868], CCN3 [869], MEG3 [870], ABCB1 [871], DPP4 [872], FNDC5 [873], CD28 [874], ADAMTS5 [875], NT5E [876], COL13A1 [877], NR2E1 [878], KLF14 [879], MMP8 [880], GPR84 [881], HP (haptoglobin) [882], PCSK9 [883], PRTN3 [884], GDF15 [885], RETN (resistin) [886], OLFM4 [887], S100A8 [888], ELANE (elastase, neutrophil expressed) [889], SEPTIN4 [890], TWIST2 [891], LCN2 [892], FAP (fibroblast activation protein alpha) [537], MMP9 [893], GCKR (glucokinase regulator) [894], IL27 [273], S100A9 [274], SDC1 [895], BPI (bactericidal permeability increasing protein) [896], CXCL10 [897], GPER1 [898], SOCS3 [899], IL10 [900], CEACAM1 [901], MPO (myeloperoxidase) [902], EDNRB (endothelin receptor type B) [565], VSIG4 [903], LTF (lactotransferrin) [904], GADD45A [905], RBP4 [906], CAMP (cathelicidin antimicrobial peptide) [907], MAOB (monoamine oxidase B) [908], PRL (prolactin) [909], OSM (oncostatin M) [910], ACHE (acetylcholinesterase) [911], AIM2 [912], ASS1 [913], IDO1 [914], CDK1 [915], FOXM1 [916], TSPO (translocator protein) [299], SOCS1 [917], ATF3 [918], IFIT3 [919], MMP2 [920], AREG (amphiregulin) [921], TLR5 [922], LRG1 [923], UPP1 [924], CCL25 [925], CCL2 [926], CMPK2 [927], CXCL11 [928], NNMT (nicotinamide N-methyltransferase) [929], TRPM2 [930], GAS6 [931], NLRC4 [932], ADM (adrenomedullin) [933], E2F1 [934] and CAV1 [935] could be an early detection markers for liver dysfunction, with a potential to be utilized as individual therapy targets. Altered expression of SEMA5A [936], DKK3 [937], EPHA4 [938], IKZF2 [939], CAMK4 [940], SFRP5 [941], BCL11B [942], IL7R [943], BACH2 [944], LEF1 [945], ERBB3 [946], TCF7 [947], IL23R [948], TRPC1 [949], LMO7 [863], WNT16 [950], MYBL1 [951], ETS1 [952], RASGRP1 [953], AFF3 [954], CD40LG [955], CCR6 [956], EBF1 [957], ANGPT1 [958], CCN3 [959], MEG3 [960], SLC4A4 [961], KLRC3 [962], ZNF667 [963], ABCB1 [964], GATA3 [965], DPP4 [966], CCL28 [967], THEM4 [968], GATA6 [969], CD28 [970], ZBTB10 [971], ITK (IL2 inducible T cell kinase [506], GPR174 [972], ZNF462 [973], CD3G [974], DNASE1L3 [975], ITGA6 [968], HAVCR1 [976], NT5E [977], MMP8 [978], S100A12 [979], PCSK9 [528], GDF15 [980], RETN (resistin) [981], OLFM4 [532], S100A8 [982], ELANE (elastase, neutrophil expressed) [983], SLC6A19 [984], C1QA [370], IGFBP2 [985], UCHL1 [986], LCN2 [987], MMP9 [988], ISG15 [989], SIGLEC1 [990], GCKR (glucokinase regulator) [991], IL27 [992], RSAD2 [993], RNASE2 [994], S100A9 [995], BCL2A1 [996], SDC1 [997], IFI44L [998], BPI (bactericidal permeability increasing protein) [999], CXCL10 [1000], USP18 [1001], GPER1 [553], SOCS3 [1002], IL10 [1003], PDCD1LG2 [1004], CXCL17 [1005], MPO (myeloperoxidase) [1006], CTSG (cathepsin G) [564], C2 [1007], LTF (lactotransferrin) [1008], PGLYRP1 [1009], IL22RA2 [1010], GADD45A [1011], FCER1G [1012], PRL (prolactin) [1013], IRF7 [1014], AIM2 [1015], IDO1 [1016], S100P [1017], IFIT1 [1018], FOXM1 [1019], SOCS1 [1020], MZB1 [1021], MMP2 [1022], IL1RN [1023], AREG (amphiregulin) [1024], TLR5 [1025], LRG1 [1026], DEFB1 [1027], MMP1 [1028], DCC (DCC netrin 1 receptor) [1029], CCL25 [1030], IL18RAP [1031], OASL (2’-5’- oligoadenylate synthetase like) [1032], OAS3 [1033], CCL2 [1034], CXCL11 [1035], FOXC1 [1036], TRPM2 [1037], GAS6 [1038], NLRC4 [449], STAB2 [1039], E2F1 [1040], CAV1 [1041] and HK3 [1042] promotes autoimmune diseases. FAT4 [1043], DKK3 [1044], TMEM30B [1045], EPHA4 [1046], CCN2 [1047], CAMK4 [1048], ENPP1 [1049], SFRP5 [1050], PTCH1 [1051], TPH1 [1052], AXIN2 [1053], ECRG4 [1054], SGCD (sarcoglycan delta) [1055], HPSE2 [1056], TRPC1 [1057], MYBL1 [1058], ETS1 [1059], FGFR2 [1060], CCR6 [1061], ROR2 [1062], CCN3 [1063], MEG3 [1064], SYNPO (synaptopodin) [1065], ABCB1 [1066], GATA3 [1067], DPP4 [656], CCL28 [1068], MCF2L2 [1069], CD28 [1070], RORC (RAR related orphan receptor C) [1071], CTNND2 [1072], ITK (IL2 inducible T cell kinase) [1073], SH3YL1 [1074], HAVCR1 [253], CD207 [1075], ADAMTS5 [1076], EPHX2 [1077], GALNT12 [1078], KLF14 [1079], MMP8 [1080], HP (haptoglobin) [1081], S100A12 [1082], PCSK9 [1083], GDF15 [1084], RETN (resistin) [1085], RNASE1 [1086], OLFM4 [1087], S100A8 [1088], RAP1GAP [1089], ELANE (elastase, neutrophil expressed) [1090], SLC6A19 [1091], SEPTIN4 [1092], TWIST2 [1093], FOSL1 [1094], IGFBP2 [1095], UCHL1 [1096], LCN2 [1097], MMP9 [1098], ISG15 [1099], SERPING1 [1100], GCKR (glucokinase regulator) [1101], VNN1 [1102], SDC1 [1103], BPI (bactericidal permeability increasing protein) [1104], CXCL10 [279], MT2A [1105], GPER1 [1106], SOCS3 [282], IL10 [1107], MPO (myeloperoxidase) [1108], VSIG4 [1109], LTF (lactotransferrin) [1110], RBP4 [1111], PFKFB2 [1112], CDC20 [1113], PRL (prolactin) [1114], OSM (oncostatin M) [1115], ACHE (acetylcholinesterase) [1116], BIRC5 [1117], AIM2 [1118], IDO1 [1119], TSPO (translocator protein) [1120], SOCS1 [1121], ATF3 [1122], DDAH2 [1123], BCAM (basal cell adhesion molecule) [1124], MMP2 [1125], ALPL (alkaline phosphatase, biomineralization associated) [751], AREG (amphiregulin) [1126]. LRG1 [1127], CDC25A [1128], GRINA (glutamate ionotropic receptor NMDA type subunit associated protein 1) [1129], CCL2 [308], GPR4 [1130], TRPM2 [1131], GAS6 [309], ADM (adrenomedullin) [1132], INHBA (inhibin subunit beta A) [1133], CAV1 [1134] and APOL4 [1135] are potentially associated with higher risk of kidney dysfunction. IL7R [1136], HPSE2 [1137], CD28 [1138], HP (haptoglobin) [1139], S100A12 [1140], PCSK9 [1141], OLFM4 [1142], S100A8 [1143], MMP9 [1144], VNN1 [1145], S100A9 [1146], SDC1 [1147], BPI (bactericidal permeability increasing protein) [1148], CXCL10 [1149], IL10 [1150], MPO (myeloperoxidase) [287], ORM1 [398], C2 [1007], DDAH2 [1151], IL1RN [1152], C3AR1 [1153], GAS6 [1154] and ADM (adrenomedullin) [1155] were related to septic shock. Therefore, the occurrence and development of sepsis and its associated complications are complicated and these enriched genes in GO terms and signaling pathway might provide clues for the common underlying mechanisms between them. These findings provide a set of useful driving genes and key pathways of sepsis and its associated complications, which are worth future investigating for novel therapeutic targets, a prognostic evaluation index, and the detailed molecular pathogenesis of them in sepsis and its associated complications.

To explore the molecular pathogenesis of sepsis and its associated complications, we constructed PPI network for systematic analysis. Hub genes were obtained through the PPI network and modules, and they were all significantly regulated in sepsis group compared with the control group. PRKN (parkin RBR E3 ubiquitin protein ligase) [128], GATA3 [113], ERBB3 [89] and ETS1 [97] are mainly involved in brain dysfunction. FGFR2 [486], GATA3 [496], ERBB3 [477], CDC20 [576] and ETS1 [484] genes have been demonstrated to be responsible for the progression of inflammation. Studies have confirmed that FGFR2 [642], ERBB3 [637], CDK1 [739], CDC20 [724], ETS1 [641] and RRM2 [775] are altered expressed in heart dysfunction. FGFR2 [790] and ETS1 [789] altered expression have been closely associated with the lung dysfunction. FGFR2 [867], CDK1 [915] and ETS1 [865] have been revealed to be altered expression in liver dysfunction. FGFR2 [1060], GATA3 [1067], CDC20 [1113] and ETS1 [1059] have been identified to be involved in the development of kidney dysfunction. Studies had shown that GATA3 [341], CDK1 [418] and ETS1 [333] were associated with microbial infections. GATA3 [965], ERBB3 [946] and ETS1 [952] might be a potential therapeutic target for autoimmune diseases. Based on topological algorithms in this inveatigation, KIT (KIT proto-oncogene, receptor tyrosine kinase), PPARG (peroxisome proliferator activated receptor gamma), H2BC5, H4C4, PAX5, CDT1 and DTL (denticleless E3 ubiquitin protein ligase homolog) were identified as critical genes that might be potential novel biomarkers for sepsis and its associated complications. ROC curve analysis showed that these hub genes have diagnostic value for sepsis. This investigation might provide reference for research the connection between sepsis and its associated complications.

In this study, we also constructed a miRNA-hub gene regulatory network and TF-hub gene regulatory network for the hub genes. The primary function of miRNAs and TFs is to regulate the expression of the hub genes in the cell. In this investigation, the miRNA and TFs, were shown to be potential modulators of sepsis and its associated complications. ITGB8 [119], ETS1 [97], BIRC5 [188], E2F1 [224], CAV1 [225], hsa-mir-3145-3p [1156], hsa-mir-2113 [1157], hsa-mir-223-3p [1158], hsa-mir-652-3p [1159], hsa-mir-106b-5p [1160], YAP1 [1161] and ATF3 [1162] have been reported to encourage the development of brain dysfunction. ITGB8 [119], ETS1 [484], ITGA6 [510], LEF1 [475], FGFR2 [486], GATA3 [496], BIRC5 [582], E2F1 [619], CAV1 [621], hsa-mir-223-3p [1163], YAP1 [1164], BCL3 [1165], ATF3 [1166], RUNX2 [1167], TBX5 [1168] and TET1 [1169] are important in the development of inflammation. ETS1 [641], FGFR2 [642], BIRC5 [731], CAV1 [621], CDK1 [739], hsa-mir-182-5p [1170], hsa-mir-223-3p [1171], hsa-mir-652-3p [1172], hsa-mir-106b-5p [1173], YAP1 [1164], BCL3 [1174], ATF3 [1175], RUNX2 [1176], TBX5 [1177] and KIF20A [776] were important in the occurrence and development of heart dysfunction. Recent studies have proposed that the ETS1 [789], LEF1 [785], FGFR2 [790], CAV1 [850], hsa-mir-182-5p [1178], hsa-mir-3679-3p [1179], hsa-mir-223-3p [1180], YAP1 [1181], BCL3 [1182], ATF3 [301] and RUNX2 [1183] are associated with lung dysfunction. ETS1 [865], FGFR2 [867], E2F1 [934], CAV1 [935], CDK1 [915], hsa-mir-3679-3p [1184], YAP1 [1185], ATF3 [1186] and TET1 [1187] are a biomarkers which plays a role in diagnosis of liver dysfunction. ETS1 [1059], FGFR2 [1060], GATA3 [1067], BIRC5 [1117], CAV1 [1134], hsa-mir-223-3p [1188], hsa-mir-106b-5p [1189], YAP1 [1190], BCL3 [1191], ATF3 [1192] and CUX1 [1193] are molecular markers for the diagnosis and prognosis of kidney dysfunction. Previous studies had shown that the altered expression of ETS1 [952], ITGA6 [968], LEF1 [945], E2F1 [1040], CAV1 [1041], hsa-mir-182-5p [1194], hsa-mir-223-3p [1195] and YAP1 [1190] were closely related to the occurrence of autoimmune diseases. Altered expression of ETS1 [333], PIK3C2B [346], GATA3 [341], BIRC5 [408], CAV1 [453], CDK1 [418], CDC6 [442], hsa-mir-2116-3p [1196], hsa-mir-223-3p [1197], hsa-mir-652-3p [1198], hsa-mir-106b-5p [1199], YAP1 [1200], BCL3 [1201], ATF3 [1202], [1203] and KIF20A [459] were found to be substantially related to microbial infections. E2F1 [311], CAV1 [312], hsa-mir-223-3p [1203], YAP1 [1181], ATF3 [301] and RUNX2 [1183] expression might be regarded as an indicator of susceptibility to sepsis. These analyses led to the identification of PAK3, ANK3, PPARG (peroxisome proliferator activated receptor gamma), CDC45, hsa-mir-548ad-5p, hsa-mir-548ay-3p, PHC1 and ESRRB as key biomarkers that could be of mechanistic relevance for sepsis and its associated complications molecular pathogenesis and progression.

In conclusion, the current investigation identified key genes and pathways which might be involved in sepsis and its associated complications progression through the bioinformatics analysis of NGS dataset. These results may contribute to a better understanding of the molecular mechanisms which underlie sepsis and its associated complications, and provide a series of potential biomarkers. However, further experiments are required to verify the findings of the current.. In vivo and in vitro investigation of gene and pathway interaction is essential to delineate the specific roles of the identified genes, which might help to confirm gene functions and reveal the molecular mechanisms underlying sepsis and its associated complications.

## Acknowledgement

I Robert EW Hancock, University of British Columbia, Microbiology and Immunology, 2259 Lower Mall Research Station, Vancouver, Canada, very much, the author who deposited their NGS dataset GSE185263, into the public GEO database.

## Ethical approval

This article does not contain any studies with human participants or animals performed by any of the authors.

## Informed consent

No informed consent because this study does not contain human or animals participants.

## Availability of data and materials

The datasets supporting the conclusions of this article are available in the GEO (Gene Expression Omnibus) (https://www.ncbi.nlm.nih.gov/geo/) repository. [(GSE185263) https://www.ncbi.nlm.nih.gov/geo/query/acc.cgi?acc=GSE185263]

## Consent for publication

Not applicable.

## Author Contributions

B. V. - Writing original draft, and review and editing

C. V. - Software and investigation

